# Brain-wide projections and differential encoding of prefrontal neuronal classes underlying learned and innate threat avoidance

**DOI:** 10.1101/2022.03.31.486619

**Authors:** Michael W. Gongwer, Cassandra B. Klune, João Couto, Benita Jin, Alexander S. Enos, Rita Chen, Drew Friedmann, Laura A. DeNardo

**Author notes:** Denotes equal contribution (Michael W. Gongwer, Cassandra B. Klune). Correspondence: Dr. Laura DeNardo, 10833 Le Conte Ave, Center for Health Sciences, David Geffen School of Medicine at UCLA, Los Angeles, CA 90095.

## Abstract

To understand how the brain produces behavior, we must elucidate the relationships between neuronal connectivity and function. The medial prefrontal cortex (mPFC) is critical for complex functions including decision-making and mood. mPFC projection neurons collateralize extensively, but the relationships between mPFC neuronal activity and brain-wide connectivity are poorly understood. We performed whole-brain connectivity mapping and fiber photometry to better understand the mPFC circuits that control threat avoidance. Using tissue clearing and light sheet fluorescence microscopy we mapped the brain-wide axon collaterals of populations of mPFC neurons that project to nucleus accumbens (NAc), ventral tegmental area (VTA), or contralateral mPFC (cmPFC) in mice. We present DeepTraCE, for quantifying bulk-labeled axonal projections in images of cleared tissue, and DeepCOUNT, for quantifying cell bodies. Anatomical maps produced with DeepTraCE aligned with known axonal projection patterns and revealed class-specific topographic projections within regions. During threat avoidance, cmPFC and NAc-projectors encoded conditioned stimuli, but only when action was required to avoid threats. mPFC-VTA neurons encoded learned but not innate avoidance behaviors. Together our results present new and optimized approaches for quantitative whole-brain analysis and indicate that anatomically-defined classes of mPFC neurons have specialized roles in threat avoidance.

## Introduction

The medial prefrontal cortex (mPFC) is a vastly interconnected brain region that controls complex cognitive and emotional functions including memory, decision making, social interactions and mood^1–3^. mPFC acts through its axonal projections to exert top-down control, biasing activity in downstream brain regions to promote adaptive behavioral responses in dynamic circumstances^4–9^. With the widespread use of opto- and chemogenetics to manipulate neural populations, a large number of studies have linked individual mPFC projections to discrete behavioral functions^5,6,8,16^. However, many mPFC-dependent behaviors are known to involve multiple downstream brain regions^17^. Recent work revealed the vast anatomical heterogeneity of mPFC projection neurons and their extensive axon collaterals^18^. Thus, there is a need to integrate anatomical findings with functional work to gain a deeper understanding of how classes of mPFC projection neurons contribute to behavior.

Many studies use retrograde viral approaches to activate, inhibit, or observe the activity of projection-defined populations^19–21^. In many cases, activation of axon collaterals beyond the projection of interest may inadvertently influence animal behaviors. Interpreting results of manipulation or observation of the somatic activity of virally- or genetically-targeted projection neurons therefore requires understanding their full projection patterns.

Traditional studies of axon collaterals have established an important foundation of knowledge but have been limited by low-throughput approaches. Anatomical mapping strategies include injecting retrograde tracers into multiple target areas and measuring overlap in source neurons^5,22^. Functional mapping strategies include recording from source neurons while performing multisite antidromic stimulation in target regions^23,24^. While informative, these approaches are biased in the sense that users predefine the target areas to investigate and are limited to small numbers of regions. RNA-sequencing based approaches^25^ allow higher throughput examination of collateralization, but the spatial resolution is limited by tissue micro-dissection volumes and analyses are also often biased to experimenter-defined sets of regions. The Allen Connectivity Atlas^26^ and the recent publication of a single neuron projectome of mouse prefrontal cortex greatly improved our understanding of prefrontal projection patterns^18^. While highly informative, serial two-photon tomography and single neuron reconstruction require specialized equipment and are computationally and time-intensive, making these approaches impractical for most scientists to replicate.

On the other hand, tissue clearing^27,28^ combined with light sheet fluorescence microscopy (LSFM) is fast, easy and affordable and can preserve 3-D circuit architecture without the need to align structures. These approaches are useful for comparing the projection patterns of genetically, anatomically or behaviorally-defined neurons and generating new hypotheses that can be tested using functional methods. For instance, users may be interested in identifying novel circuits underlying behavior or in determining how genetic, environmental, or pharmacological perturbations affect structural and functional connectivity. With commercial lightsheet systems now readily available, these approaches are within reach for many labs. But we need end-to-end open-source packages for performing whole-brain quantitative analyses. To address this deficit, we developed DeepTraCE (<u>Deep</u> learning-based <u>Tra</u>cing with <u>C</u>ombined <u>E</u>nhancement) for quantifying bulk-labeled fluorescent axons and DeepCOUNT for quantifying fluorescently labeled cell bodies. These analysis pipelines build on the machine learning-based image segmentation package TrailMap^29^. Together, they comprise a user-friendly, analysis package for high-throughput, whole brain analysis based on tissue clearing and LSFM.

Advances in computer vision allow researchers to train classifiers to automatically detect axons in tissue volumes^29,30^. However, classifiers sometimes fail to identify all axons because axon appearance varies across brain regions. Axon morphology, including the extent and symmetry of branching as well as the tortuousness of branches, varies by projection type^18^. Variance in axonal appearance is also partly due to the resolution limitations of most light sheet microscopes that are capable of visualizing entire mouse brains in a timely fashion. Axons can appear thick and bright in sparsely innervated brain regions near the surface of the brain, but thin and indistinct in densely innervated regions deep in the tissue. To overcome these hurdles, DeepTraCE allows users to combine multiple models that have been separately trained using TrailMap to accurately identify axons with differing appearances. After segmenting the axons using TrailMap models, DeepTraCE registers the brains to a common coordinate framework and quantifies innervation density by brain region. Similarly, DeepCOUNT uses TrailMap to detect cells and then quantifies cell bodies by brain region.

In this study, we used DeepTraCE, DeepCOUNT and fiber photometry to better understand the mPFC circuits that control threat avoidance. Threat avoidance behavior is critical for survival. While different brain regions have been implicated in innate and learned avoidance, the prelimbic (PL) subregion of mPFC is integral to both^31–33^. But our understanding of how PL circuits control threat avoidance remain incomplete. We focused on PL neurons that project to the contralateral PL (cPL), nucleus accumbens (NAc) or ventral tegmental area (VTA). PL-cPL, PL-NAc and PL-VTA neurons have distinct distributions across cortical layers, distinct genetic profiles, and distinct functions^5,6,34^. We mapped the brain-wide collateral projections of each neuronal class. DeepTraCE accurately detected axons throughout the brain, outperforming other model combination methods. Next, we used DeepCOUNT and activity-dependent genetic labeling to map brain-wide neuronal activation patterns after mice avoided threats. We used these data to construct functional networks and found that PL was one of the most highly connected nodes. Further, PL’s functional targets were regions that are preferentially targeted by axon collaterals of cPL, NAc or VTA projectors. Using fiber photometry, we discovered class-specific activity during innate and learned avoidance. Together, our work reveals that projection-defined PL classes have specialized roles in threat avoidance. We also demonstrate the utility of combining common techniques like fiber photometry with DeepTraCE and DeepCOUNT to understand the detailed structural and functional connectivity of the cells being studied.

## Results

### DeepTraCE Workflow

We developed DeepTraCE, an open-source, end-to-end analysis pipeline for quantifying bulk axonal projection patterns in cleared brains. DeepTraCE takes in raw images of fluorescently labeled axons and then applies TrailMap^29^, a machine learning pipeline that trains a 3D U-net framework to automatically identify axons in cleared tissue (Figure 1A, Figure S1). TrailMap provides a pre-trained model that users can fine tune to fit their samples. However, it was challenging to train a model that could sufficiently generalize across the diversity of mPFC axonal structures in all of their target regions; some axons were bright and delineated (e.g. in cortical areas) while others were dense and indistinct (e.g. in the basolateral amygdala and thalamus). To surmount this hurdle, DeepTraCE allows users to combine multiple trained models to optimize axon detection throughout the brain. After importing 3-D image stacks, multiple models trained using TrailMap fully segment each brain. DeepTraCE then registers each brain to a standard atlas and generates a new image in which pixel values in each region were extracted from the assigned model. Axons in the resulting images are then thinned to single-pixel width^29^ and the number of axon-containing pixels are quantified for each brain region (Figure 1A).

**Figure 1.**
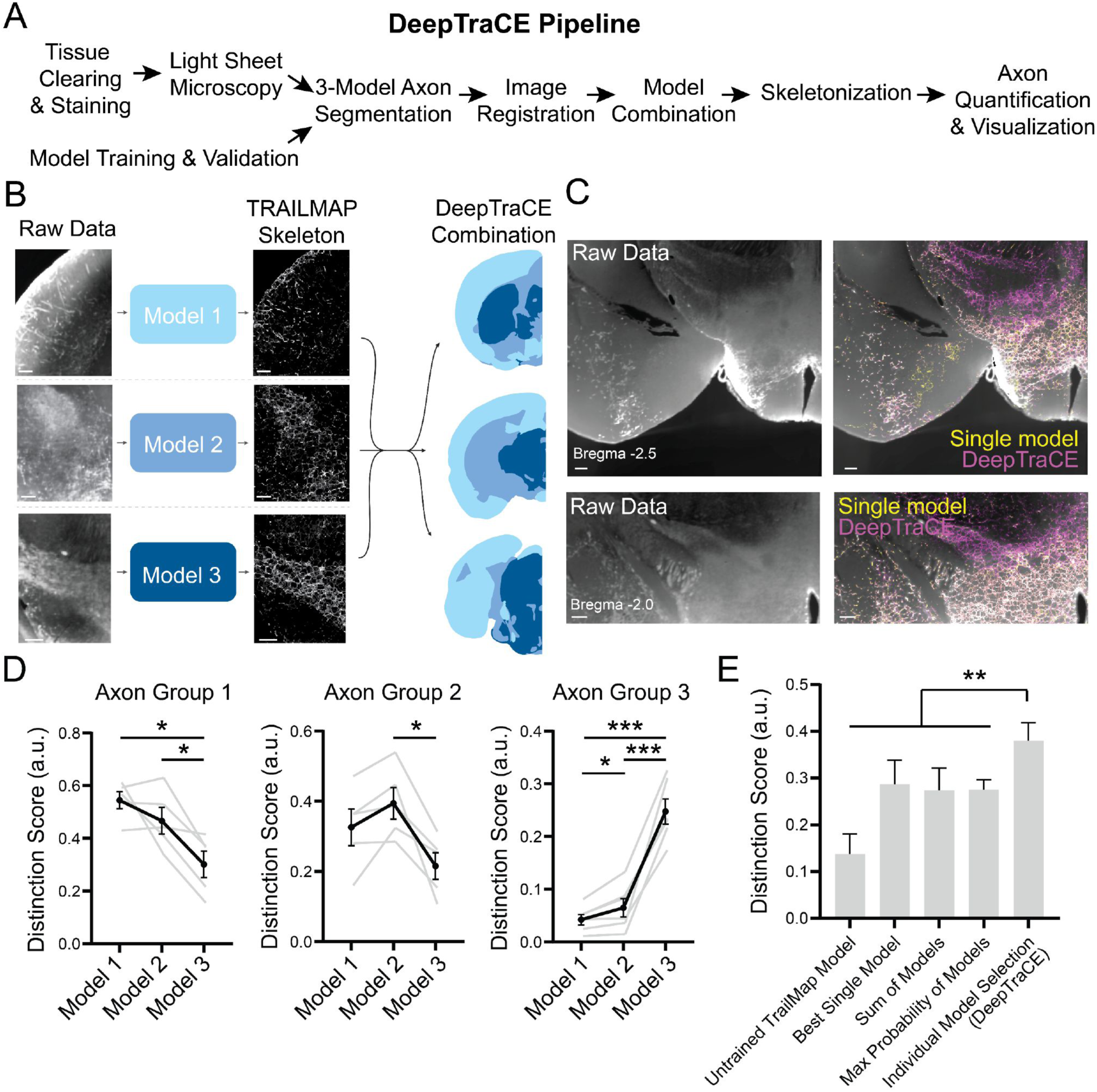
DeepTraCE workflow. (A) Overview of DeepTraCE workflow (B) Demonstration of segmentation using 3 different models followed by model combination (C) Overlay of raw data and axon segmentation using single model or DeepTraCE concatenated models (D) Distinction score produced by models 1, 2, and 3 for images from regions assigned to axon groups 1, 2, and 3. Repeated-measures one-way ANOVA with Benjamini, Krieger and Yekutieli post-hoc test. (Group 1: *F*=8.503, *P*=0.0257; Group 2: *F*=7.753, *P*=0.0309; Group 3: *F*=73.76, *P*=0.0001). (E) Comparison of DeepTraCE with alternate segmentation and model combination strategies. Repeated measures one-way ANOVA with Benjamini, Krieger and Yekutieli post-hoc test. (*F*=14.46, *P*<0.0001). See Table 3 for detailed statistics. Scale bars, 200um. Descriptive statistics: mean ± SEM. \**P*<0.05, \*\**P*<0.01, \*\*\**P*<0.001. Distinction score: (LP-LN) / (LN+(1-LP)). LP = axon+ human label. LN = axon-human label.

### Validation of model combination method

To determine how many models to combine, we examined raw images of fluorescently-labeled axons taken with LSFM. We grouped brain regions into three main categories based on the visual quality of PL axons (e.g. thick and delineated or dense and fuzzy). These categories largely segregated with the physical location of brain regions in tissue volumes (Figure 1B). We used TrailMap^29^ to train three separate models that we fine-tuned for each set of regions (Figure 1B,C; Figure S1A–C).

To validate the DeepTraCE model assignments, we performed quantitative comparisons of model accuracy in representative brain regions from each category, using human annotations as a reference. Human experts traced pixels containing axons in image stacks from each group of brain regions (Figures S1A–C). We segmented images using each of the three models in addition to the original untuned TrailMap model that was trained on serotonergic axons^29^. We then calculated the resulting axonal labeling density with reference to human labels.

In LSFM of cleared tissue, the size of small fluorescent structures is amplified such that a thin object such as an axon occupies several more pixels in the image than it does in true biological space. We account for this overrepresentation by thinning segmented axons to a single pixel width^29^. However, because rapid imaging of intact rodent brains requires lower optical resolution, it can be challenging in some regions to hand-label axons with single pixel precision. This caveat to whole-brain light sheet microscopy approach prevents the calculation of “true positive” and “true negative” rates based on human annotation. So, common metrics such as F1 Score, precision, and recall do not provide interpretable measures of segmentation accuracy. Instead, we calculated a ‘distinction score’ that measures the density of segmented axons in regions that humans estimated as containing axons vs. not containing axons. A higher distinction score indicates better overlap with human labels. When applied to our validation set, Models 1, 2, and 3 produced the highest distinction scores in images from the brain regions assigned to each model (Figure 1D).

Previous studies used alternative approaches to overcome the limitations of a single-model approach in whole-brain image segmentation. Some excluded deeper, more densely innervated areas from analysis^35^. Others combined probability maps from different segmentation methods by extracting the maximum probability for each pixel^36^. To compare DeepTraCE with these alternative approaches, we calculated the distinction score across all human-annotated images in the validation set using the original TrailMap model that was trained on serotonin neurons, our best-trained single model, the maximum probability between our three models, the sum of probabilities, and the DeepTraCE combined models. DeepTraCE had the highest distinction score (Figure 1E), indicating that it performed more accurate axon segmentation from images.

### Viral circuit mapping and brain clearing strategy

We mapped the brain-wide axon collaterals of PL neurons that project to cPL, NAc or VTA. Our goals were 1) to demonstrate the utility of DeepTraCE and 2) to provide an important anatomical resource for researchers studying mPFC function. Therefore, we used a commonly employed dual-virus approach to fluorescently label populations of PL projection neurons^37^. In contrast to multi-site retrograde tracer injections, this approach allows us to visualize the bulk brain-wide collateralization patterns of each projection-defined class of neurons. We first injected an axon-terminal-transducing adeno-associated virus^38^ expressing Cre recombinase (AAVrg-hSyn-Cre) into cPL, NAc, or VTA. Next, into PL, we injected AAV expressing Cre-dependent EYFP-tagged Channelrhodopsin-2 (AAV8-hSyn-DIO-ChR2-EYFP), which traffics efficiently to axons (Figure 2A-C). We used the iDISCO clearing variant, Adipo-Clear^39,40^ to immunostain intact brains for EYFP and render the tissue transparent. We then imaged the hemispheres ipsilateral to the PL injection with LSFM.

**Figure 2.**
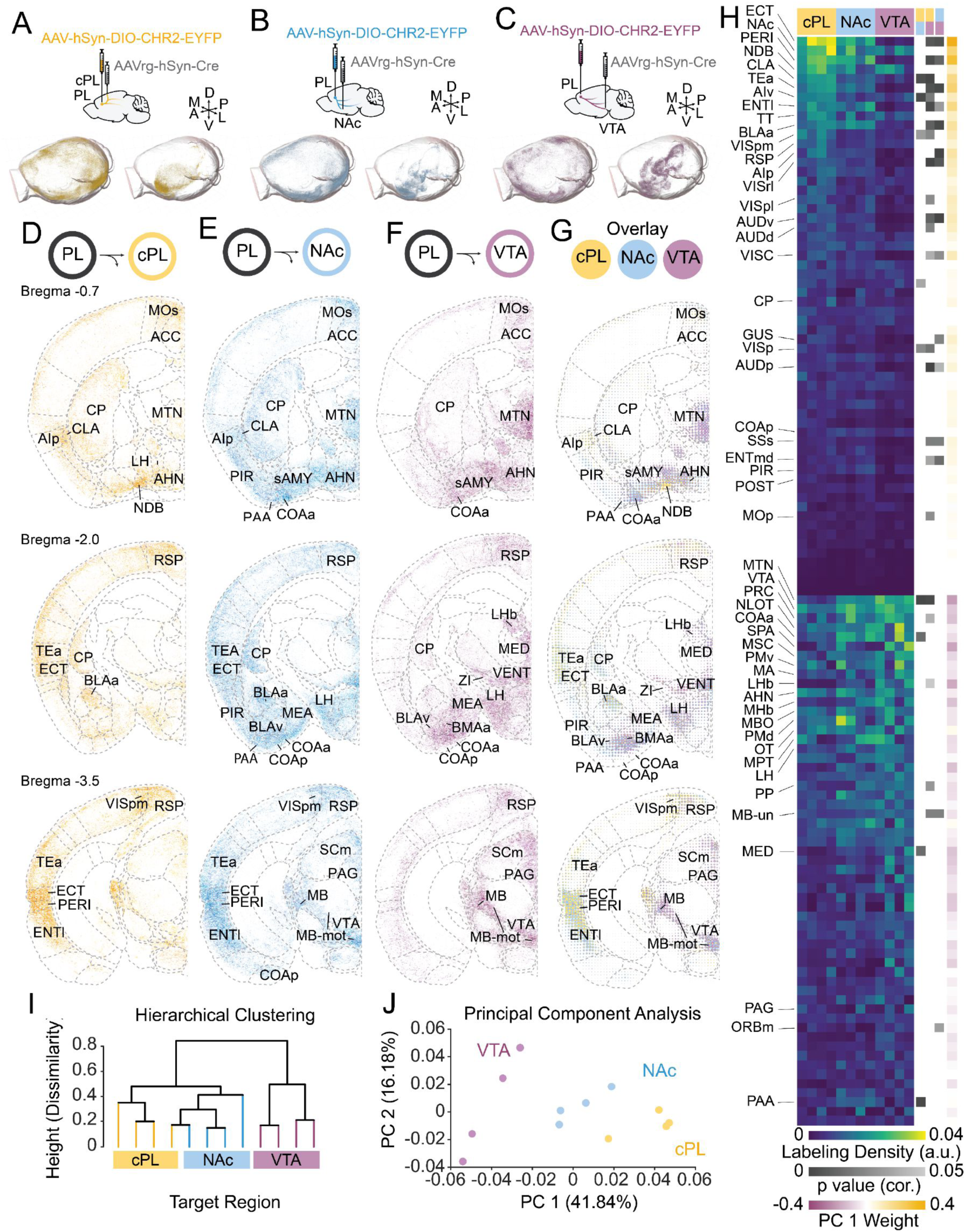
Visualization and quantification of brain-wide projection patterns of mPFC. (A–C) Viral injection strategy and DeepTraCE segmentation of superficial (left) and deep (right) brain structures for each cell type. (D–F) Single 10um coronal optical sections of thinned axons registered to the standardized brain atlas. Represents 4 overlaid brains from each neuronal class. (G) Dotogram overlay of 3 cell types. Innvervation by cPL-, NAc- and VTA-projecting PL neurons shown in yellow, blue and purple, respectively. (H) Left heatmap: relative labeling density (normalized to region volume and gross label content per brain) across 140 regions defined by the Allen Brain Atlas. Middle heatmap: P values from multiple comparisons of axon innervation density. Right heatmap: Loadings for PC1 (arbitrary PC weight units) (I) Dendrogram of hierarchical clustering of regional axon quantifications from 12 mice, colored by target region. (J) Locations of individual mice projected in principal component (PC) space defined by the first two PCs (arbitrary PC units, cPL, *n*=4; NAc, *n*=4; VTA, *n*=4). Abbreviations: Isoctx, isocortex; OLF, olfactory areas; HPF, hippocampal formation; CTXsp, cortical subplate; CNU, cerebral nuclei; HY, hypothalamus; MB, midbrain; HB, hindbrain. See Tables 1–3 and Figure S2-6 for related data.

In separate animals, we used confocal microscopy to examine the distribution of retrogradely labeled cell bodies in brain sections (Figure S2A). Different retrograde tracers can have preferences for particular neuronal types. For instance, while AAVrg favors cortical layer (L)5 over L6, rabies virus has the opposite preference^41^. Cholera toxin subunit B (CTB) is inefficient for labeling pyramidal tract neurons in L5^42^. To assess potential biases in our retrograde labeling method, we analyzed the layer distribution for AAVrg-Cre-mCherry+ cells and the combination of AAVrg-hSyn-Cre-mCherry and AAV8-hSyn-DIO-ChR2-EYFP (assessed based on EYFP fluorescence). As a comparator, we measured the layer distribution of CTB+ cells in PL following injections into cPL and NAc. Consistent with previous reports, we did not observe retrogradely labeled PL cells when we injected CTB into the VTA^42^. We determined layer boundaries based on DAPI nuclear staining in combination with immunostaining for Ctip2, a marker of subcerebral projection neurons located in L5b-6^43^. Layer boundaries and thicknesses aligned with previous literature (Figure S2B)^11,44,45^. Consistent with previous findings^45–47^, PL-cPL and PL-NAc neurons were preferentially located in L2–5a and PL-VTA neurons were restricted to L5b. We found no statistical differences in the distribution of cells labeled with AAVrg alone, AAVrg+AAV8 or CTB (Figure S2A-C). As reported previously, we observed few AAVrg-labeled cell bodies in L6^41,42^. While L6 neurons were likely underrepresented in our datasets, we could still extract meaningful differences between classes.

### Whole-brain projection patterns of PL-cPL, PL-NAc and PL-VTA neurons

We used DeepTraCE to determine the extent to which PL-cPL, PL-NAc and PL-VTA neurons collateralize to other brain regions. Examination of thin optical sections revealed widespread collateralization for all three classes. Overall, innervation patterns were consistent with reports of general mPFC projection patterns^22,45,48^; mPFC axons were prominent in cortical association areas, striatum, midline thalamus, claustrum, amygdala, hypothalamus and midbrain but largely absent from hippocampus, sensory thalamus, and relatively sparse in primary sensory areas (Supplemental Video; Figure 2D–F; Figure S3–5).

We generated dotograms^26^ to visually summarize the inter-class distinctions. To create the dotogram, we averaged axon density across all samples for a given projection class and then overlaid the samples in brain space. Dot color indicates the neuronal class and dot size corresponds to the averaged axonal density within a voxel (Figure 2G). Compared to PL-VTA neurons, PL-cPL and PL-NAc neurons more densely innervated cortical areas, especially the temporal association area (TEa), ectorhinal area, and entorhinal cortex. On the other hand, PL-VTA collaterals preferentially innervated thalamic (TH) and midbrain (MB) regions, a subset of which also received collateral input from PL-NAc neurons. All three subclasses innervated NAc and olfactory tubercle (OT), while PL-NAc neurons were the primary source of collaterals to piriform cortex (PIR).

We next quantified regional innervation densities and plotted them as a heatmap. For each brain, we normalized the number of axon-containing pixels in each region to the total number of axon-containing pixels in the brain. We then sorted regions according to their innervation density by projection classes; regions receiving more PL-cPL collaterals are on top and those receiving more PL-VTA collaterals are in the bottom half of the heatmap (Figure 2H). While there was little overlap between projection patterns of PL-cPL and PL-VTA neurons, PL-NAc neurons shared several projection targets with both other classes. Brain-wide statistical comparisons confirmed the presence of 29 significantly differentially innervated brain regions, the most notable of which were in the cortex, subplate, and thalamus (Figure 2H, S6, Table 2, Table 3).

Because PL-cPL, PL-NAc and PL-VTA projection neurons all collateralize broadly, we wondered if these neuronal classes could be separated based on their whole-brain projection patterns. To test this, we used two dimensionality reduction techniques. We first performed hierarchical clustering, in which Euclidean distance between individual brains based on axonal density in each subregion is marked by a higher branch point on the graph. PL-VTA neurons formed a distinct cluster, while PL-NAc and PL-cPL collaterals formed partially overlapping clusters (Figure 2I). This aligns with our observation that the projection patterns of PL-NAc neurons shared more overlap with PL-cPL than PL-VTA neurons (Figure 2H).

We also used principal component analysis to assess differences in whole-brain projection patterns for each cell class. We plotted the location of each brain along the axes of the first two principal components. While PL-cPL and PL-VTA projection classes could be clearly separated along the first principal component, PL-NAc brains were positioned in between, with some overlap with PL-cPL brains (Figure 2J). To assess which projection patterns distinguished the classes, we plotted the weights of each brain region in contribution to the first principal component. Positively weighted regions generally had the most innervation from PL-cPL collaterals including TEa, PERI, and ECT, and negatively weighted regions were those with most innervation from PL-VTA collaterals, including MTN and midbrain areas (Figure 2H). These data show that while cPL and NAc-projectors have substantial overlap, PL-cPL and PL-VTA neurons represent separable classes, distinguished in large part by their collaterals to TEa/PERI/ECT, olfactory and limbic areas, and thalamus and midbrain, respectively. Together, the innervation patterns we observed were consistent bulk projection mapping^26,49,50^ and single neuron reconstructions^18,51^, indicating that DeepTraCE accurately captured meaningful class-specific distinctions in collateral targeting.

### Layer-specific and topographic innervation patterns in the cortex and subplate

Whole-brain analysis of bulk labeled neuronal classes is well-suited for understanding the geometric organization of axonal projections *within* brain regions, which can have important functional implications. The neocortex is organized into layers that contain distinct neuronal types with different morphology, physiology, and connectivity^44,52^. Long-range cortico-cortical axons may innervate superficial or deep layers depending on the hierarchical relationship with the target region^44,52^. Compared to sensory and motor cortices, much less is known about the layer organization of long-range mPFC connectivity, especially for specific projection classes. To investigate this, we plotted PL-cPL, PL-NAc and PL-VTA collateral innervation density across layers in cortical target areas (Fig. 3A-B). In ORB, all classes preferentially targeted superficial layers (L1, L2/3). In AUD, TEa, ECT, and PERI, PL-cPL and PL-NAc collaterals preferentially targeted superficial layers (L1, L2/3) while PL-VTA collaterals targeted the deep layers (L5, L6) (Figure 3B, Table 3). In ENT, axon collaterals were more prominent in deep layers for all 3 classes. These findings suggest that PL-cPL, PL-NAc and PL-VTA may all participate in feedforward or feedback connectivity depending on the target region.

**Figure 3.**
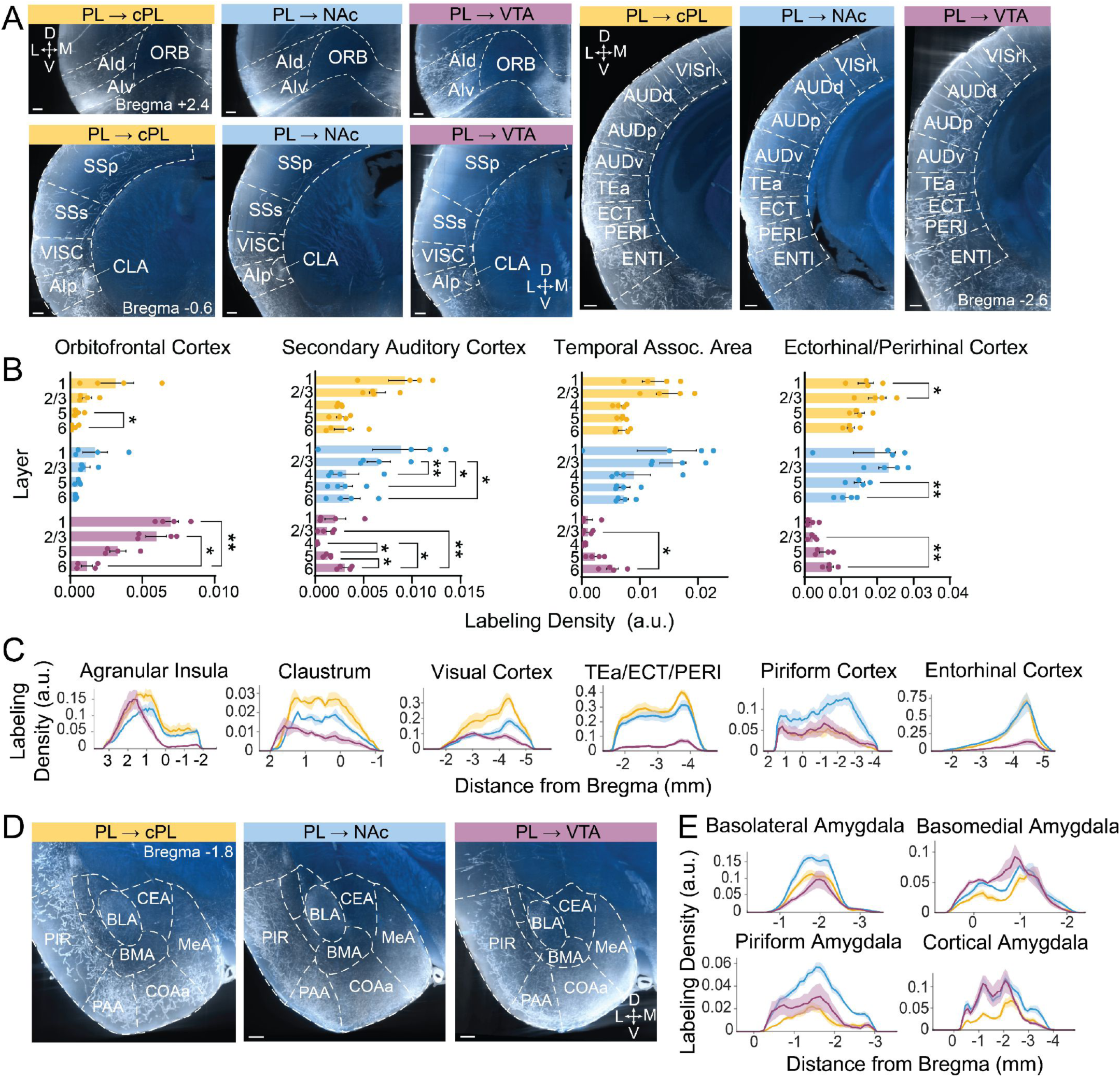
Region-specific collateralization patterns of PL-cPL, PL-NAc and PL-VTA neurons. (A) Coronal view of 100um z-projections of raw 640nm (axons, white) and 488nm (autofluorescence, blue) channels from individual brains. Images show cell-type specific innervation patterns in anterior (left) and posterior (right) cortical areas. (B) Layer distributions of axonal innervation in select cortical target regions. (C) Visualizations of axonal innervation density along the anterior-posterior axis in select regions. (D) Raw images showing axons in the amygdalar complex. (E) Quantification of axonal innervation density along the anterior-posterior axis of a given region. PL-cPL, PL-NAc and PL-VTA neurons coded in yellow, blue and purple, respectively. N=4/group. Descriptive statistics are from 2-way ANOVA with Tukey’s post-hoc multiple comparison test. See Table 1,3 for abbreviations and detailed statistics. Error bars: mean ± S.E.M. \**P*<0.05, \*\**P*<0.01.

Many PL target regions contain functional gradients that are orthogonal to layer-defined microcircuits. We therefore investigated whether PL classes project to topographically-defined locations in target regions (Figure 3C). PL-NAc collaterals were especially prominent in the posterior PIR. Compared to PL-cPL and PL-NAc, PL-VTA collaterals had a tighter distribution in the anterior portion of agranular insula (AI). PIR and AI both have contain anatomical and functional gradients^53,54^. Our data suggest that class-specific PL projections may be a determining factor in these gradients.

While all projection classes sent collaterals to the amygdalar complex, amygdalar nuclei were differentially innervated. Most notably, the anterior basolateral amygdala (BLAa) was robustly innervated by PL-cPL and PL-NAc neurons, but less so by PL-VTA neurons. Meanwhile, compared to PL-cPL neurons, PL-NAc neurons sent many more collaterals to olfactory amygdalar areas, including anterior cortical amygdalar area (COAa) and piriform amygdalar area (PAA) (Figure 3D,E). As PL-NAc collaterals were also more prominent in posterior PIR, which was recently found to play a role in spatial cognition^53^, PL-NAc neurons may have privileged control over areas dedicated to olfactory processing during cognition and emotional learning.

### Whole-Brain Cell Counting Pipeline

The application of computer vision to light sheet data extends beyond axon tracing and can be used to identify signatures of brain function. Expression of IEGs such as c-fos are frequently used as a proxy of neural activity^55,56^. For instance, recent studies have used tissue clearing, IEG-labeling, and LSFM to screen for changes neuronal activation following experiences with drugs or fear learning^28,48,57,58^. Quantifying IEG-expressing cells on brain-wide scale can serve as means to infer behaviorally-relevant changes in functional connectivity. However, currently available open-source packages for whole-brain cellular quantification can be error prone, especially in regions of particularly dense labeling.

As a companion to DeepTraCE, we developed DeepCOUNT (<u>Deep</u>-learning based <u>C</u>ounting of <u>O</u>bjects via 3D <u>U</u>-<u>N</u>et pixel <u>T</u>agging) (Figure 4A). We first trained a TrailMap model to recognize fluorescently labeled cell bodies. This produces a probability map of which pixels are most likely to contain a cell body. After thresholding the images to a desired probability cutoff, they are registered to a standard brain, and DeepCOUNT uses a 3D maxima detection strategy is used in combination with a connected component analysis to ensure each neuron is represented by a single pixel. This single-pixel output can then be used to obtain regional cell counts across the brain.

**Figure 4.**
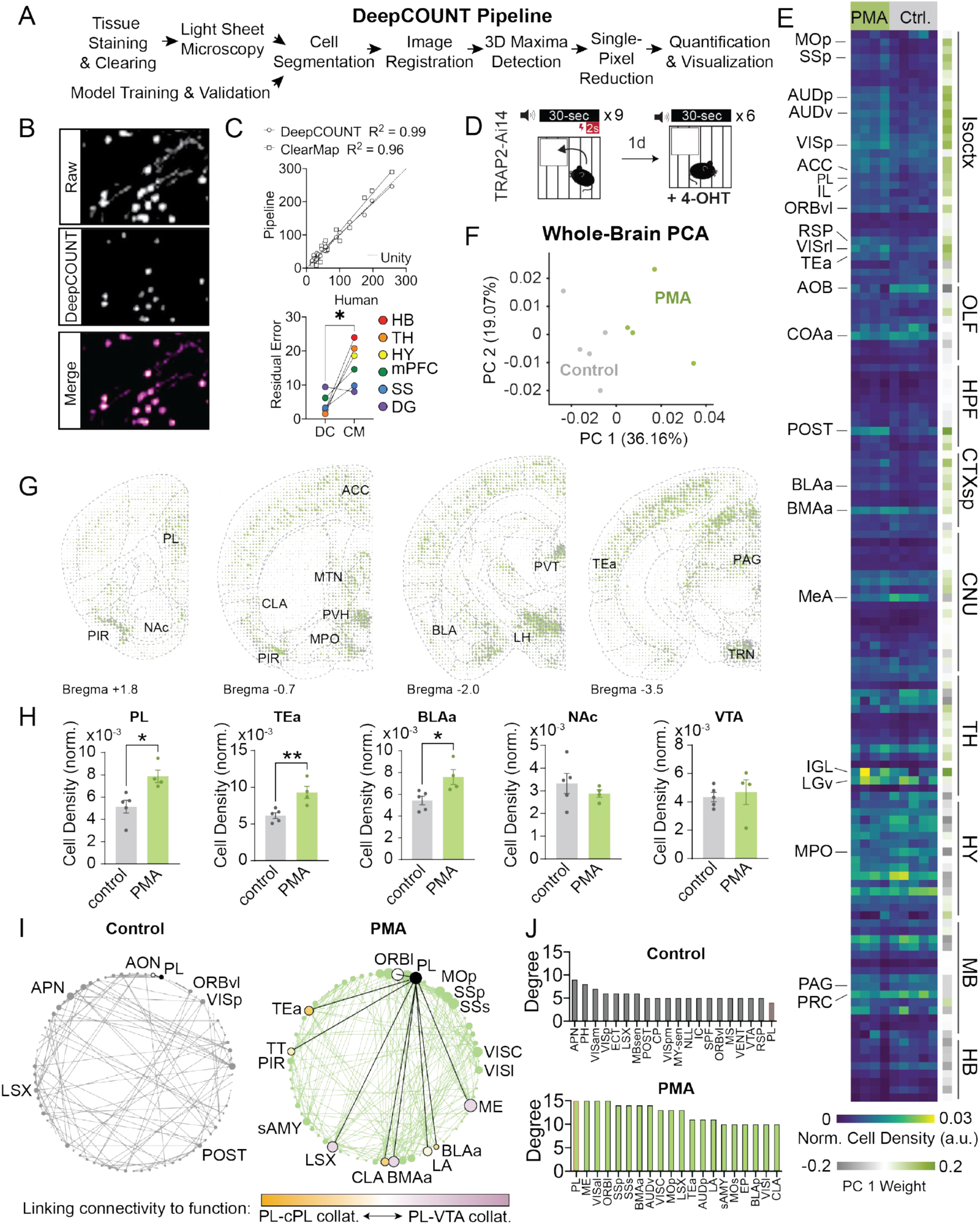
Whole-brain IEG mapping of learned avoidance with DeepCOUNT. (A) Overview of DeepCOUNT workflow (B) Overlay of raw data and axon segmentation using trained cell detection model. (C) Upper: Correlation between human vs. DeepCOUNT or ClearMap neuronal counts in image volumes (Pearson’s R^2^_DC_ = 0.99, *P*<0.0001; R2_CM_ = 0.96, *P*<0.0001; Lower: Comparison of residual error values for DeepCOUNT and ClearMap as compared to human annotations (*P*=0.022, n=6 brain regions; paired t-test). (D) Schematic of platform mediated avoidance assay. TRAP2;Ai14 mice were trained in PMA and then TRAPed during a retrieval test the next day. (E) Left heatmap: relative cell density (normalized to region volume and gross label content per brain) across 140 regions defined by the Allen Brain Atlas. Right heatmap: Loadings for PC1 (arbitrary PC weight units). (F) Locations of individual mice projected in principal component (PC) space defined by the first two PCs (arbitrary PC units, control, *n*=5; PMA, *n*=4). (G) Dotogram overlay of control (grey) and PMA (green) conditions. Dots represent the density of TRAPed cells in a given voxel. (H) Comparison of TRAPed cell density in 5 brain regions of naïve controls (grey) and PMA-trained mice (green) (Student’s t-test; *\*P*<0.05, \*\**P*<0.01; control: *n*=5; PMA: *n*=4). (I) Network diagrams for control (left) and PMA-trained (right) mice based on brain wide interregional correlations. Node size is proportional to degree. (J) Degree values for top 20 most connected regions for each condition (PL outlined in red). See Figure S6 for related data.

We compared DeepCOUNT performance to that of ClearMap^28^, a commonly used cell detection algorithm designed for tissue clearing and LSFM, and human observers. Two human observers analyzed image volumes from six different brain regions. Cell counts produced with DeepCOUNT were highly correlated with human observers and DeepCOUNT produced more accurate counts than ClearMap, using human counts as the ground truth (Figure 4B,C).

### Whole-brain threat avoidance networks

We used DeepCOUNT to map regions and functional connections involved in threat avoidance at a whole-brain level. PL is required for threat avoidance behavior^32,59^. Previous studies examining c-fos expression following active avoidance behavior analyzed only discrete regions of interest^60–62^, some of which are PL target sites. How PL coordinates threat avoidance-related neural activity on a brain-wide scale is not understood. To begin to address this question, we used DeepCOUNT and activity-dependent genetic labeling to map brain-wide neuronal activation following a platform mediated avoidance (PMA)^32^, a form of learned threat avoidance.

We labeled activated neuronal populations using TRAP2 mice (*Fos^icre-2A-ERT2^*), in which the c-fos promoter drives expression of tamoxifen-inducible Cre recombinase. We crossed TRAP2 to the Ai14 Cre reporter line^48,63,64^. We then trained the *TRAP2;Ai14* mice in PMA and TRAPed them after a threat avoidance retrieval session by pairing that experience with a tamoxifen injection (Figure 4D, S7A). We used DeepCOUNT to quantify TRAPed cell density in each brain region and plotted these data as a heatmap (Figure 4E). Principal component analysis of whole-brain TRAPed cell counts separated PMA-trained and non-shocked control animals along the first principal component (PC1) (Figure 4F). PC1 weights revealed that PMA-activated brain regions were biased toward the cortex, hippocampal formation, and cortical subplate (Figure 4E). We plotted TRAPed cell density across the brain as dotograms (Figure 4G) and compared cell counts between PMA and control animals in PL and several key targets. Our DeepCOUNT screen identified regions known to be involved in threat avoidance (e.g. PL and BLA). We also identified several regions with unknown roles in threat avoidance (e.g. the TEa and postsubiculum (POST)) that will be of interest for future functional studies (Figure 4E,H).

To better understand the organization of PL circuits that control threat avoidance, we assessed functional connectivity based on whole-brain TRAPing. We computed inter-regional correlations for groups of PMA and control mice. This allowed us to identify sets of regions where the numbers of TRAPed cells co-varied across mice (Figure S7B). Regions that co-vary may constitute elements of a network engaged during learned threat avoidance. We generated network graphs for each condition based on the strongest correlations (Pearson’s r>0.9, P<0.05, Figures 4I, S7C).

Our network analysis revealed PL as one of the mostly highly connected nodes in the PMA group (Figure 4J). PL was functionally connected to several cortical areas (TEa, ORBl, MOp, SSp, SSs), taenia tecta (TT), the lateral septal complex (LSx), claustrum (CLA), several amygdalar regions (LA, BLAa, BMAa), and the median eminence (ME). Our neuroanatomical analysis had already revealed that some these regions are preferentially innervated by PL-cPL (CLA, TEa, TT) or PL-VTA (LSx). Compared to PL-VTA, PL-NAc and PL-cPL neurons send denser projections to BLAa. Compared to PL-cPL neurons, PL-NAc and PL-VTA neurons send denser projections to BMAa. To illustrate this, we colored the PL-connected nodes based on their PCA weight from the analysis shown in Figure 2 (Figure 4I). These data suggest that all three neuronal classes likely contribute to threat avoidance behavior and that they may act via particular collateral projections.

### Fiber Photometry During Platform-Mediated Avoidance

We investigated how different PL cell classes contribute to threat avoidance by using fiber photometry to record population-level neural activity. Given the high overlap of collateral projection targets for PL-cPL and PL-NAc neurons (Figures 2I,J), we further separated these classes using an intersectional viral strategy^65^ (Figure 5A). To record neuronal activity in PL-cPL neurons that do not project to NAc, we injected AAVrg-Cre into cPL and AAVrg-Flp into NAc.

**Figure 5.**
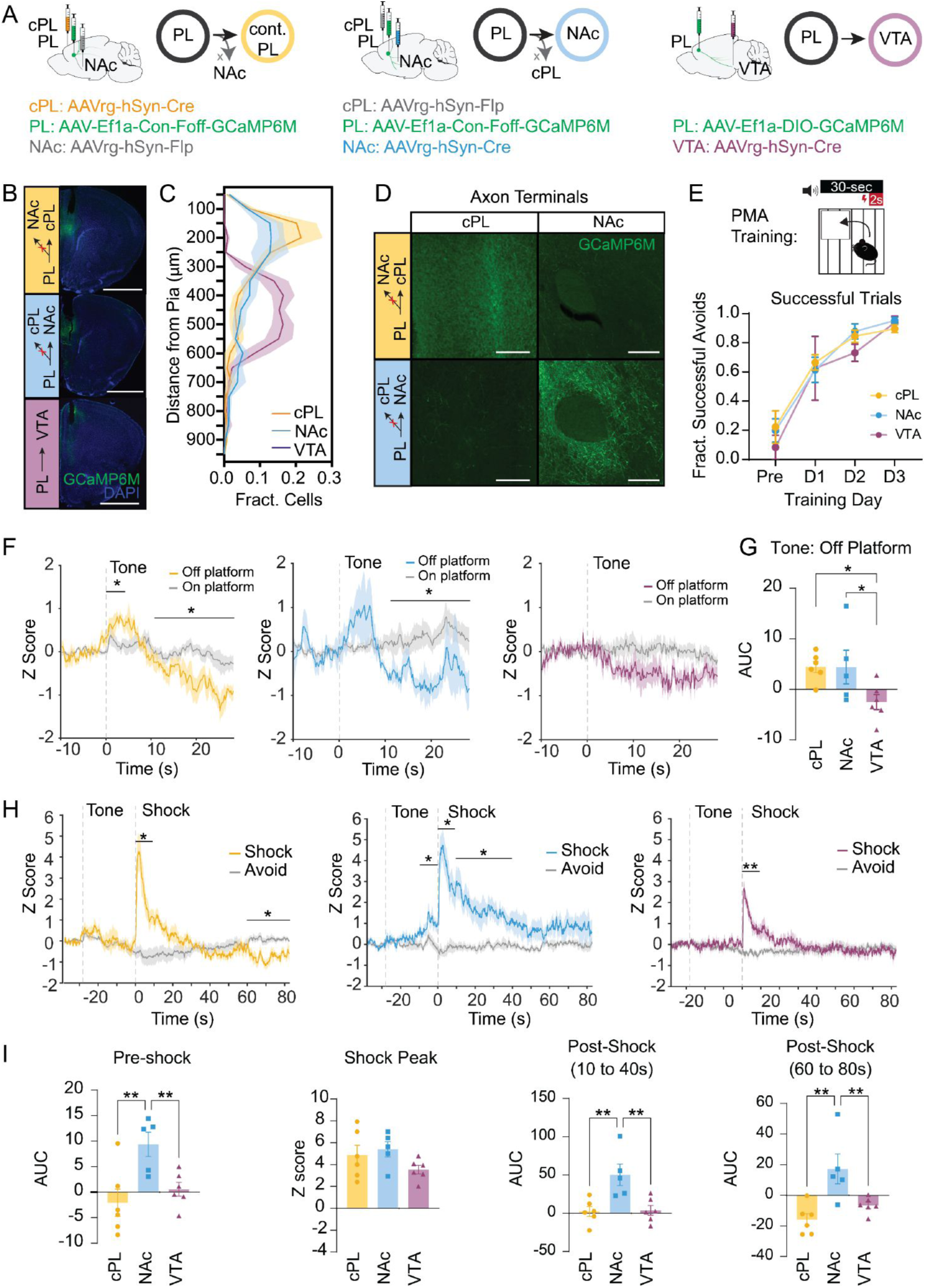
Neuronal class-specific activity during threat cues and aversive stimuli. (A) GCaMP injection strategy. Left: to target cPL-projecting neurons, mice were injected with AAVrg-Cre in cPL and AAVrg-Flp in NAc, and Con-Foff-GCaMP6M in PL. Middle: to target NAc-projecting neurons, mice were injected with AAVrg-Flp in cPL and AAVrg-Cre in NAc, and Con-Foff-GCaMP6M in PL. Right: to target VTA-projecting neurons, mice were injected with AAVrg-Cre in VTA and DIO-GCaMP6M in PL. (B) Representative images of GCaMP expression and fiber placement sites in PL. Scale bar, 1mm. (C) Distribution of cPL-, NAc- and VTA-projecting GCaMP-expressing cells across cortical layers (*F_bin_* = 28.3, *P*<0.0001, *F_binxclass_* = 34.09, *P*<0.0001; cPL: *n*=6, NAc: *n*=5, VTA: *n* =6; 2-way ANOVA). (D) Representative images of axon terminals in cPL and NAc using the intersectional viral targeting strategy shown in A. Scale bars, 100um. (E) Schematic of PMA assay and behavioral performance across sessions (*F_day_* = 65.18, P<0.0001, *F_class_* = 0.58, *P*=0.57, *F_class x day_* = 0.43, *P*=0.86, 2-way ANOVA). (F) GCaMP fluorescence in cPL-, NAc-, and VTA-projecting neurons. Signals are aligned to tone onset and separated by whether mouse is on (colored trace) or off (grey trace) the safety platform at the start of the tone. \**P*<0.05 for student’s t-test comparing on vs. off platform activity in a given time window. (G) AUC analysis of Ca^2+^ signal for tone periods (0–10s) when mice were off the platform (*F* = 4.106, *P*=0.04, cPL: *n*=6, NAc: *n*=5, VTA: *n* =6; One-way ANOVA with Benjamini, Krieger and Yekutieli post-hoc test. (H) GCaMP fluorescence in cPL-, NAc- and VTA-projecting PL cells. Signals are aligned to shock onset and separated by whether mouse is on (colored trace) or off (grey trace) the safety platform. \**P*<0.05, \*\**P*<0.01 for student’s t-test comparing on vs. off platform activity in a given time window. (I) Analysis of Ca^2+^ signals for circa-shock periods (Pre-shock AUC: *F* = 6.974, *P*=0.0079; Shock AUC: *F* = 1.868, *P*=0.19; Post-shock AUC (10–40s): *F* = 8.28, *P*=0.0042; Post-shock AUC (80–80s): *F* = 8.79, *P*=0.0034; cPL: *n*=6, NAc: *n*=5, VTA: *n* =6, One-way ANOVA with Benjamini, Krieger and Yekutieli post-hoc test).

We then injected Cre-On Flp-Off GCaMP6M into PL. We switched the Cre and Flp injection sites to record from NAc-projecting neurons that do not project to cPL. To record from VTA-projecting neurons, we injected AAVrg-Cre into VTA and DIO-GCaMP6M into PL (Figure 5B). Most GCaMP^+^ cell bodies from the intersectionally-defined cPL- and NAc-projecting neurons were in superficial layers of PL, while cell bodies from VTA-projecting neurons were restricted to the deeper layers (Figure 5C), as expected from our initial layer analysis (Figure S2A). We observed few GCaMP^+^ axon terminals in NAc from the cPL-projecting group, and few GCaMP^+^ axon terminals in cPL from the NAc-projecting group (Figure 5D), suggesting our intersectional strategy was efficient in separating these populations.

We recorded from PL while mice performed PMA across 3 days of training. By training day 3, mice successfully avoided most shocks (Figure 5E). We first analyzed responses to the conditioned tone. Interestingly, tone-evoked neural activity distinguished between epochs when mice were off vs. on the safety platform (Figure 5F). cPL-projecting neurons had significantly higher activity at the tone onset when the animal was off the platform. As the tone progressed, activity in these neurons gradually decreased. We observed a similar trend in NAc-projecting neurons. In contrast, PL-VTA cells lacked a tone onset response, but did have a gradual decrease in activity during the conditioned tone (Figure 5G). These findings suggest neural activity in response to a conditioned stimulus is modulated by whether threat requires action, and that these patterns vary across projection classes.

Activity in all classes sharply increased upon foot shock onset (Figure 5H), and we found no differences in peak shock response between the three classes (Figure 5I). NAc-projecting neurons had significantly higher activity than the other two classes in the ten seconds preceding a shock (Figure 5I), consistent with a larger role for those cells in risky exploration. Further, while activity in PL-VTA neurons rapidly returned to baseline levels following the foot shock, activity in PL-cPL and PL-NAc neurons had prolonged deviations from baseline for up to a minute after the shock. Activity in NAc-projecting neurons remained elevated above baseline for the forty seconds following a foot shock, while activity in cPL-projecting neurons decreased steadily during the same period (Figure 5I,H). These data suggest that while all PL classes encode aversive stimuli, prolonged post-shock activity in PL-cPL and PL-NAc neurons may play specialized roles in action-outcome learning.

### Prefrontal Neuron Classes Distinguish between Learned and Innate Avoidance

To further classify the effects of aversive learning on neural activity in PL cell classes, we compared their activity during learned vs. innate threat avoidance. Using the same mice from the PMA experiments, we recorded neural activity in the Elevated Zero Maze (EZM), in which two quarters of an elevated ring are protected by walls and the other two quarters are open. Mice innately avoid the open arms of this apparatus, where there is more perceived threat potential.

We compared activity during entries and exits from the safe zone in each assay. In PMA, population activity in all three neuronal classes increased prior to platform entries (Figure 6A). In contrast, only NAc- and cPL-projecting neurons increased their activity prior to entry into the closed arm of the EZM (Figure 6B). During exits from the safe zone, a form of risky exploration, all three PL classes had similar increases in activity during PMA (Figure 6C). In innate avoidance, however, exits from the closed arm were marked by a significantly larger response in NAc-projecting neurons compared to the other two cell populations (Figure 6D). Together, these data indicate that aversive learning engages VTA-projecting neurons to behavioral circuits for threat avoidance and that PL-NAc neurons preferentially encode risky exploration.

**Figure 6.**
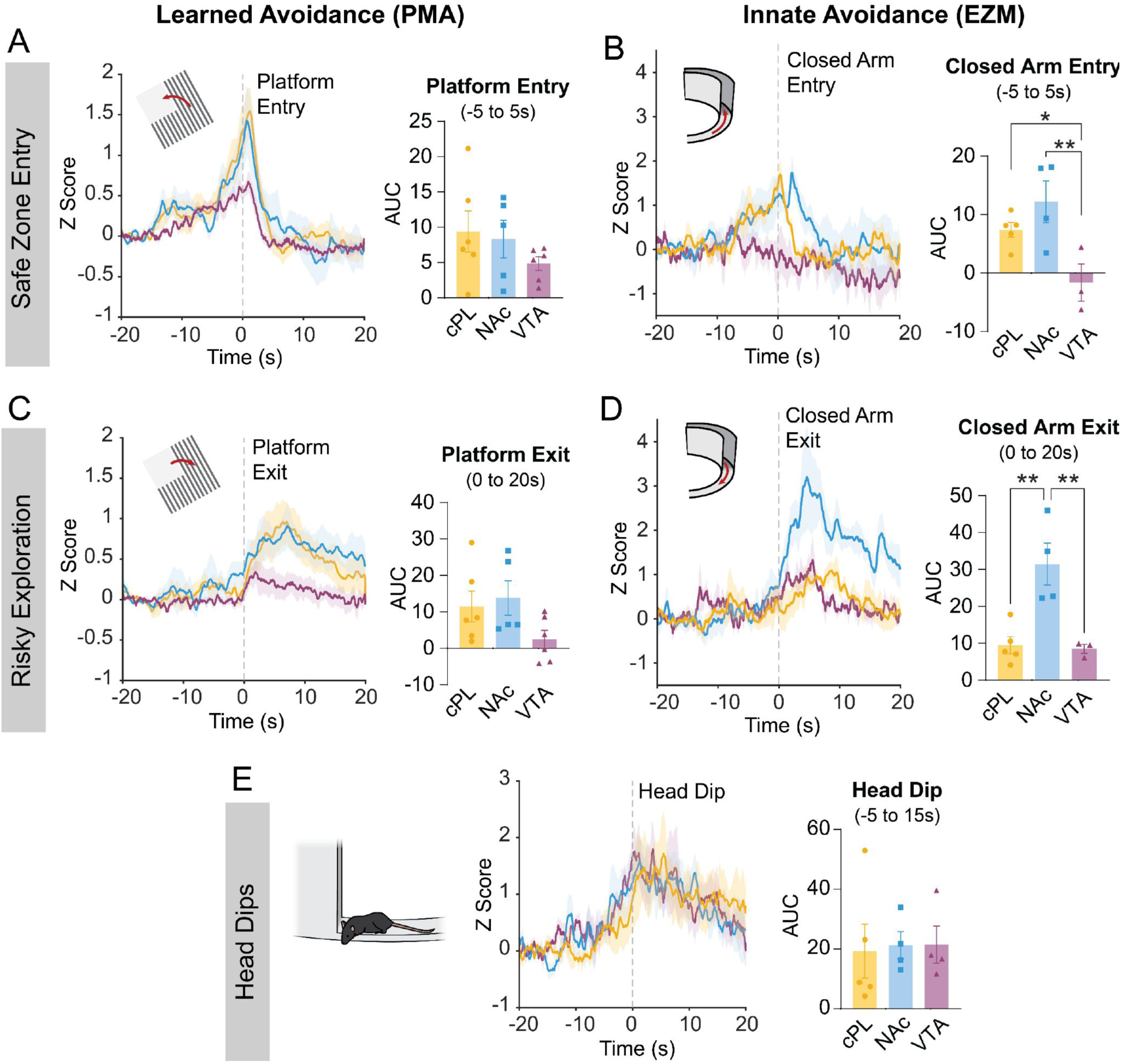
Neuronal class-specific activity during learned vs. innate threat avoidance behavior. (A,B) GCaMP fluorescence aligned to safe-zone entry in learned (A) vs. innate (B) avoidance for cPL-projecting (yellow), NAc-projecting (blue) and VTA-projecting (purple) neurons. Inset plots show AUC analysis of Ca^2+^ signal aligned to avoidance (-5–5s) (*F* = 4.106, *P*=0.04, One-way ANOVA with Benjamini, Krieger and Yekutieli post-hoc test). (C,D) GCaMP fluorescence aligned to onset of risky exploration in learned (C) vs. innate (D) avoidance for cPL-projecting (yellow), NAc-projecting (blue) and VTA-projecting (purple) neurons. Inset plots show AUC analysis of Ca^2+^ signal aligned to exploration onset (0–20s) (*F* = 4.106, *P*=0.04, One-way ANOVA with Benjamini, Krieger and Yekutieli post-hoc test). (E) Analysis of Ca^2+^ signals during head dips (*F* = 0.64, *P*=0.55; PMA: cPL *n*=6, NAc *n*=5, VTA *n*=6; EZM: cPL *n*=5, NAc *n*=4, VTA *n*=3; One-way ANOVA with Benjamini, Krieger and Yekutieli post-hoc test.).

A concern with comparing fiber photometry recordings across different classes of neurons is that differences in signal between the classes could lead to exaggerated results in one direction. While PL-VTA neurons had lower overall activity during risky exploration of the shock bars and open arm, all three PL classes had similar activity during head dips on the EZM, another form of exploratory behavior (Figure 6E). This suggests that PL-VTA neurons did not simply have lower activity than the other classes during behavior, but instead, that these differences are specific to situations when animals are navigating the environment. To further corroborate this, we performed a separate analysis of signals normalized to the peak of the average shock response for an animal, as the shock provides a consistent time-locked response in each animal. All between-group statistical comparisons remained consistent upon normalization to the shock response (Figure S8), suggesting the effects we observe are not driven by differences in signal quality between the three classes.

## Discussion

In this study, we integrated whole brain mapping with the observation of prefrontal neural activity to better understand how mPFC controls threat avoidance behavior. We introduce DeepTraCE and DeepCOUNT, two new open-source analysis pipelines that extend the utility of TrailMap to quantify bulk axonal projection patterns and IEG labelling, respectively. We used DeepTraCE to produce whole brain structural projection maps of three populations of mPFC projection neurons: PL-cPL, PL-NAc, and PL-VTA. We next combined activity-dependent genetic labeling with DeepCOUNT to relate the structural and functional organization of brain-wide networks for threat avoidance. Finally, with whole-brain projection maps as a foundation, we used intersectional viral targeting to separate the overlapping populations of PL-cPL and PL-NAc neurons and then recorded the activity of PL-cPL, PL-NAc and PL-VTA neuronal populations during learned and innate threat avoidance. Our study reveals PL class-specific roles in threat avoidance and demonstrates the utility of DeepTraCE and DeepCOUNT for linking high-throughput neuroanatomy with functional techniques to explore the reveal mechanisms of complex behavior (Figure 7).

**Figure 7.**
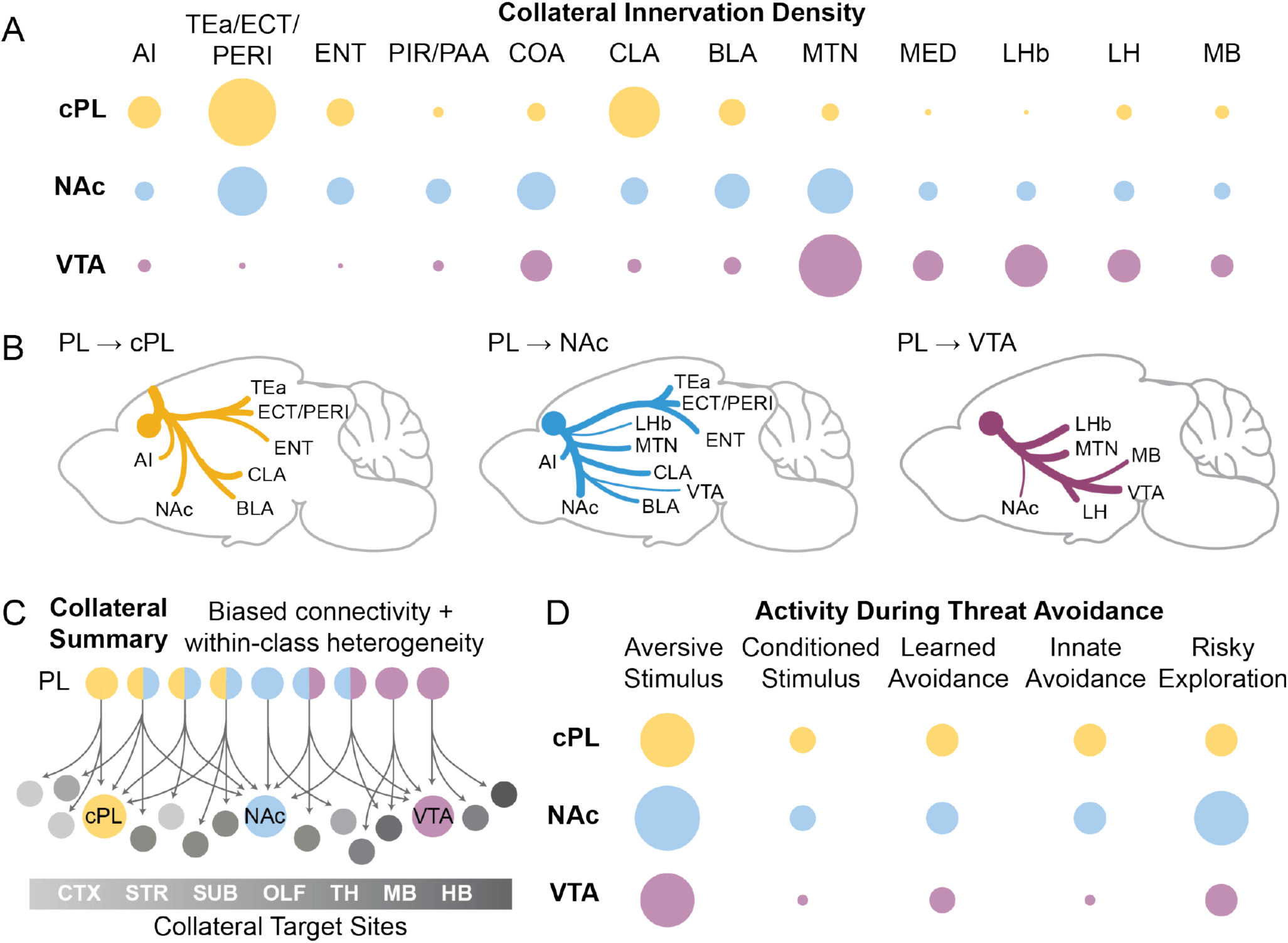
Summary of Findings. (A) Visualization of collateralization density in key targets of PL–cPL, PL–NAc, and PL–VTA neurons. Dot radius correlates with average normalized labeling density within a region. (B) Schematic of whole-brain collateralization patterns of PL–cPL, PL–NAc, and PL–VTA neurons. (C) Summary of PL efferent connectivity patterns. (D) Visualization of activity levels in PL–cPL, PL–NAc, and PL–VTA neurons during aspects of threat avoidance. Dot radius correlates with average normalized signal intensity. See Table 1 for abbreviations.

While we have gained invaluable insights into the complex anatomy of cortical neurons from single neuron reconstructions^18,26,66^, the equipment, computing power, expertise and time required to reconstruct individual neurons are out of reach for many labs. Other high-throughput techniques such as MAPseq^67^ provide single cell resolution, but are costly and limited to a predefined number of target regions. While bulk labelling approaches lose granularity of individual neuron differences, DeepTraCE’s high-throughput workflow combined with its reliance on common viral techniques makes it a useful tool to tackle a variety of unanswered questions. Capitalizing on mouse Cre-driver lines and floxed alleles, DeepTraCE can be used to analyze the projection patterns of defined neuronal populations and how they can be altered by genetic manipulations. Similarly, DeepTraCE and DeepCOUNT can be used to define differences in structural and functional connectivity resulting from environmental insults such as chronic stress. Given the sample size needed to pursue these questions, our pipeline’s high-throughput processing puts these experiments within practical reach when compared to single neuron reconstruction. With DeepTraCE, in addition to simply quantifying relative densities of axonal projections in target regions, users can assess within-region changes in layer-specific or topographic targeting. This information can guide the design of subregion-targeted functional studies. Experimenters can use the same viral-genetic strategies to drive expression of opsins or GCaMP proteins in neuronal classes of interest and then implant optic fibers with targeting informed by the anatomical maps.

It is important to also note the caveats associated with retrograde viruses, brain clearing and LSFM. Compounds for retrograde labeling can have biases against projection classes^68^. AAVrg viruses have been shown to have preferential labelling of corticothalamic (CT) neurons in L5, and biases against CT neurons in L6^41,42^, while CTB showed the opposite pattern of layer bias^42^. In line with this, we observed few AAVrg+ neurons in L6, but some CTB+ neurons (Figure S2A). So we may have underestimated the contributions of L6 cells to the population data for PL-cPL and PL-NAc neurons^22,44^. Like other retrograde labeling methods, the full range of neuronal types that may be resistant to AAVrg is not well understood. Also, tissue clearing does not always achieve complete transparency. This issue, together with resolution limits of light sheet microscopes that can visualize entire mouse brains in a timely fashion, can produce blurring of structures deep in the tissue. Therefore, estimates of axonal density may be less accurate in deep brain structures. However, estimates of axon density are likely to be biased to a similar degree for different neuronal classes in any given target structure. So, high-throughput mapping using viral tracing, brain clearing and LSFM is well-suited for comparing inter-class differences.

In our fiber photometry experiments, the intersectional viral strategy we used to parse apart PL-cPL and PL-NAc populations relied on an AAVrg injection expressing Flp recombinase to block GCaMP expression in projection populations. Our AAVrg-Flp injections did not span the entirety of cPL or NAc. Therefore, we do not claim to have completely segregated these populations but rather biased them towards each projection type. Finally, like most mapping studies, we visualized axons but not synapses. Therefore, our DeepTraCE quantifications may also include axons of passage. To mitigate this, our models were trained to exclude major white matter tracts.

Despite these caveats, our brain wide projections maps of PL-cPL, PL-VTA, and PL-NAc neurons aligned with known projection patterns^18,22,26,48,51^ – underscoring the accuracy of TrailMap^29^ and DeepTraCE – and revealed unappreciated topographic innervation patterns. Consistent with bulk tracing studies^48–50^, we observed the densest PL projections in temporal association cortices, claustrum, striatum, BLA, lateral hypothalamus, polymodal thalamic nuclei, hypothalamus and midbrain. These bulk projection patterns are the sum of multiple classes. Like sensory cortices, mPFC contains intratelencephalic (IT) neurons that project within the cortex and pyramidal tract (PT) neurons, that project to subcortical areas. PL-cPL neurons fall in the IT class, PL-VTA neurons are from the PT class, and PL-NAc contain members of both^44^. In line with these classifications and single neuron reconstructions^18,44,51^, we observed that PL-cPL neurons were enriched in superficial layers (L2-5a) and distinguished by their collaterals to other cortical association areas, subplate (CLA and BLA), and striatum. Consistent with the well-described features of PT neurons^18,44,66^, we observed retrogradely labeled PL-VTA neurons with somas localized to L5b and axon collaterals in the thalamus, pons, PAG, striatum, and neuromodulatory centers. Also consistent with prior work, PL-NAc neurons collateralized broadly to both IT and PT targets ^44^. The diversity of the collateral targets of PL-NAc neurons is supported by single neuron reconstructions which further show a variety of subclasses targeting only a few structures^18^. Overall, there was excellent alignment between our data and data collected using methods that have higher spatial resolution. This underscores the accuracy of DeepTraCE and its utility for capturing meaningful class-specific differences in connectivity in a high-throughput manner.

We also observed unappreciated topography in how each class innervated target regions. PL-NAc collaterals preferentially innervated the posterior PIR, which was recently found to have a specialized role in spatial cognition^53^. Compared to PL-cPL and PL-NAc, PL-VTA collaterals were restricted to the AI. PIR and AI are both complex anatomical hubs with distinct functional zones along the rostro-caudal axis^53,69,70^. Preferential innervation by PL projection neurons may underlie some of the functional specializations within these regions.

On a brain wide scale, counts of IEG-expressing cells can serve as means to identify behaviorally-relevant activity and functional connectivity^48,57,58^. Previous studies examining c-fos expression following active avoidance behavior analyzed only discrete regions of interest^60–62^. We used DeepCOUNT and TRAP2 mice to measure brain-wide neuronal activation following threat avoidance. DeepCount recapitulated previous findings of activation in PL, BLA and OFC following PMA^60^. We also found high numbers of TRAPed cells in POST and TEa, whose roles in threat avoidance are unknown. POST is important for spatial learning^71–73^. During PMA learning, prefrontal connections to this area may facilitate learning of the location of the safety platform. Moreover, PL projection to TEa may facilitate long-term storage of the avoidance memory as TEa is necessary for long term retrieval of cued fear^74^. Functional connectivity analysis revealed the major functional targets of PL. Of these, TEa, CLA, and TT are highly innervated by the collaterals of PL-cPL projection neurons, while LSx and BMAa are more densely innervated by PL-NAc and PL-VTA collaterals. Only PL-NAc neurons have been previously studied in active avoidance^9^. Our study highlights how whole brain anatomical and functional maps can be used together to identify novel behaviorally-relevant pathways.

We used fiber photometry to determine how each class of neuron encodes threat avoidance. We found that intersectionally-defined PL-NAc and PL-cPL neurons increased their activity in response to the conditioned tone in PMA, but only when mice were off the safety platform, suggesting those populations encode the predictability of threat. In contrast, PL-VTA neurons decreased their activity in response to the threat predictive cue. Single unit recordings haven shown that distinct populations are excited and inhibited by a conditioned tone during PMA^59^. Our results indicate that the projection targets we studied may be a factor that separates these populations.

PL-cPL connections are important for both hemispheric synchronization and lateralization of mPFC activity. While lateralization has been associated with anxiety-like behavior^75^ and stress responses in rodents^76–78^, little is known about the behavioral functions of PL-cPL neurons, especially in aversive behaviors. Our data show that PL-cPL neurons persistently tracked threat, threat predictive cues, and threat avoidance behaviors. In the future, studies manipulating these neurons will be required to understand whether these activity patterns are necessary for learned or innate avoidance.

We found that PL-NAc neurons encoded risky exploration. This is consistent with previous work showing that excitation of PL-NAc projections decreases avoidance behaviors^9^. However, individual neurons within this projection class can have heterogeneous functions. For instance, only some mPFC-NAc encoded foot shocks and promoted reward seeking under threat of punishment^6^. mPFC neurons projecting to the NAc shell play a larger role in suppression of reward seeking^79^. Given that our AAVrg targeted the NAc core, we likely captured a larger proportion of mPFC-NAc neurons involved in risk engagement rather than suppression. Our observation of PL-NAc neurons resembles the activity patterns seen in mPFC neurons projecting to dorsomedial striatum (DMS), which also increase their activity during conditioned tones and during risky exploration ^80,81^. Indeed, we see that mPFC-NAc neurons send some axon collaterals in the DMS suggesting these populations overlap.

PL-VTA populations encoded learned but not innate threat avoidance behavior. Associative learning depends on the mPFC to encode predictive cues and on VTA to encode prediction errors^82^. PL-VTA neurons may regulate the separable actions of these two regions. To our knowledge, we are the first to directly study the role of mPFC-VTA neurons in threat avoidance. However, dopaminergic VTA cells projecting to mPFC have been widely studied and shown to robustly respond to aversive stimuli and facilitate associative learning^13,83,85^. mPFC-VTA neurons form synapses onto VTA cells projecting back to mPFC, including dopaminergic neurons^86^ and may form part of a feedback loop influencing aversive learning.

In sum, we demonstrated the utility of pairing a standard approach like fiber photometry with whole-brain mapping to gain a detailed understanding of the relationships between function and connectivity. We show that PL-cPL, PL-NAc and PL-VTA neurons encode distinct features of innate and learned avoidance. Further, we mapped PL avoidance networks that that include discrete collateral targets of cPL, NAc and VTA projecting neurons. These whole-brain maps are an important resource for researchers studying the behavioral functions of mPFC. They can inform new experiments manipulating different PL pathways during threat avoidance and other disease-relevant behaviors. Further, our open-source analysis packages make brain wide anatomical approaches more accessible for integration into a variety of future research.

## Methods

### Animals

Female and male C57B16/J mice (JAX Stock No. 000664) were group housed (2–5 per cage) and kept on a 12 hr light cycle. All animal procedures followed animal care guidelines approved by the University of California, Los Angeles Chancellor’s Animal Research Committee.

### Surgery

Mice were induced in 5% isoflurane in oxygen until loss of righting reflex and transferred to a stereotaxic apparatus where they were maintained under 2% isoflurane in oxygen. Mice were warmed with a circulating water heating pad throughout surgery and eye gel was applied to the animal’s eyes. The mouse’s head was shaved and prepped with three scrubs of alternating betadine and then 70% ethanol. Following a small skin incision, a dental drill was used to drill through the skulls above the injection targets. A syringe pump (Kopf, 693A) with a hamilton syringe was used for injections. Injections were delivered at a rate 75nL/min and the syringe was left in the brain for 7 minutes following injection. For collateralization mapping, 300uL of AAVrg-Ef1a-mCherry-IRES-Cre-WPRE (Addgene 55632-AAVrg, 1.7x10^13^ vg/mL) was injected unilaterally into either left NAc (AP: 1.3, ML: -1.0, DV: 4.7), left VTA (AP: -3.3, ML: 0.4, DV: -4.5), or right PL (AP: 1.8, ML: -0.5, DV: -2.3). 200uL of AAV8-hSyn-DIO-hCHR2(H134R)-EYFP-WPRE (1.7x10^13^ vg/mL) was then injected into left PL (AP: 1.8, ML: -0.4, DV: -2.3). To control for Cre-independent expression of EYFP, control mice received a single injection of 200uL of AAV8-hSyn-DIO-hCHR2(H134R)-EYFP-WPRE into left PL. For fiber photometry from cPL and NAc-projectors, we injected AAVrg-Ef1a-mCherry-IRES-Cre-WPRE (same as above) or AAVrg-EF1a-mCherry-IRES-Flpo-WPRE (Addgene 55634-AAVrg, 1.7x10^13^ vg/mL) into either cPL or NAc, and after 2 weeks of expression injected AAV8-EF1a-Con-Foff 2.0-GCamp6m-WPRE into PL (Addgene 137120-AAV8). For recordings from PL-VTA neurons, we injected AAVrg-Ef1a-mCherry-IRES-Cre-WPRE (same as above) into VTA and AAV5-CAG-GCamp6m-WPRE (Addgene 100839-AAV5) into PL. 400uM optic fibers (Doric) were implanted into left PL (AP: 1.8, ML: -0.4, DV: -2.3) and sealed in place using Metabond (Patterson Dental Company, 5533559, 5533492, S371). For topographical mapping of the origin of each projection type, mice were injected with 300uL of cholera toxin subunit B (ThermoFisher, C34775, C34776, C34778) or Fluorogold (SCBT, C223769) at the same coordinates for cPL, NAc and VTA listed above. For pain management mice received 5mg/kg carprofen diluted in 0.9% saline subcutaneously. Mice received one injection during surgery and daily injections for two days following surgery. Samples with mistargeted injection sites were excluded from analysis. Samples with obviously poor antibody penetration or distribution following tissue clearing were also excluded.

### Brain Slice Histology and Immunostaining

Mice were transcardially perfused with phosphate-buffered saline (PBS) followed by 4% paraformaldehyde (PFA) in PBS. Brains were dissected, post-fixed in 4% PFA for 12–24h and placed in 30% sucrose for 24–48 hours. They were then embedded in Optimum Cutting Temperature (OCT, Tissue Tek) and stored at -80°C until sectioning. 60um floating sections were collected into PBS. Sections were washed 3x10min in PBS and then blocked in 0.3% PBST containing 10% normal donkey serum (JacksonImmunoresearch, 17-000-121) for 2h. Sections were then stained with chicken anti-GFP (AVES 1020 at 1:2000), rabbit anti-RFP (Rockland 600-401-379 at 1:2000) or rat anti-CTIP2 (Abcam ab18465, 1:200) in 0.3% PBST containing 3% donkey serum overnight at 4°C. The following day, sections were washed 3x5min in PBS and then stained with secondary antibody (JacksonImmunoresearch Cy2 donkey anti-chicken IgG(H+L) 703-225-155, 1:1000, Cy3 donkey anti-rabbit IgG(H+L) 711-005-152, 1:1000 or Alexa647 donkey anti-rat IgG(H+L) 702-605-150, 1:500) in 0.3% PBST containing 5% donkey serum for 2 hours at room temperature. Sections were then washed 5 min with PBS, 15 min with PBS+DAPI (Thermofisher Scientific, D1306, 1:4000), and then 5 min with PBS. Sections were mounted on glass slides using FluoroMount-G (ThermoFisher, 00-4958-02) and then imaged at 10x with a Leica STELLARIS confocal microscope.

### Brain Clearing

Mouse brains were collected and processed based on the published Adipo-Clear protocol^39^ with slight modifications. Mice were perfused intracardially with 20mL of PBS (Gibco) followed by 4% paraformaldehyde (PFA, Electron Microscopy Sciences) on ice. Brains were hemisected approximately 1mm past midline and postfixed overnight in 4% PFA at 4°C. The following day, samples were dehydrated with a gradient of methanol (MeOH, Fisher Scientific):B1n buffer (1:1,000 Triton X-100, 2% w/v glycine, 1:10,000 NaOH 10N, 0.02% sodium azide) for 1 hour for each step (20%, 40%, 60%, 80%) on a nutator (VWR). Samples were then washed with 100% MeOH 2x for 1hr each and then incubated in a 2:1 dicholoromethane (DCM):MeOH solution overnight. The following day, two washes of 1hr in 100% DCM were performed followed by three washes of 100% MeOH for 30min, 45min then 1hr. Samples were bleached for 4 hours in 5:1 H2O2/MeOH buffer. A cascade of MeOH/B1n washes (80%, 60%, 40%, 20%) for 30min each rehydrated the samples followed by a 1h wash in B1n buffer. 5%DMSO/0.3M Glycine/PTxWH permeabilized tissue for one hour and then again for 2h with fresh solution. Samples were washed with PTxwH for 30min and then incubated in fresh PTxwH overnight. The following day two more PTxwH washes lasted 1h then 2h. Samples were incubated in primary GFP antibody (AVES Labs GFP 1020) at 1:2000 in PTxwH shaking at 37°C for 11, washed in PTxwH 2x1h and then 2x2h, then for two days with at least one PTxwH change per day while shaken at 37°C. Samples were then incubated in secondary antibody (AlexaFluor 647, ThermoFisher Scientific) for 8 days shaken at 37°C. Samples were washed in PTxwH 2x1h, then 2x2h, then 2 days with at least one PTxwH change per day while shaken at 37°C. Samples were then washed in 1x PBS twice 1x1hr, 2x2hr and then overnight. To dehydrate samples, a gradient of washes in MeOH:H2O (20%, 40%, 60% and 80%) were conducted for 30min each, followed by 3x100% MeOH for 30min, 1h, then 1.5h. Samples were incubated overnight in 2:1 DCM:MeOH on a nutator. The next day, samples were washed in 100% DCM 2x1h each. Samples were then cleared in 100% DBE. DBE was changed after 4h. Samples were stored in DBE in a dark place at room temperature. Imaging took place at least 24h after clearing.

### Whole Brain Imaging

Brain samples were imaged on a light-sheet microscope (Ultramicroscope II, LaVision Biotec) equipped with a sCMOS camera (Andor Neo) and a 2x/0.5 NA objective lens (MVPLAPO 2x) equipped with a 6 mm working distance dipping cap. Image stacks were acquired at 0.8x optical zoom using Imspector Microscope v285 controller software. For axons, we imaged using 488-nm (laser power 20%) and 640-nm (laser power 50%) lasers. The samples were scanned with a step-size of 3 µm using the continuous light-sheet scanning method with the included contrast adaptive algorithm for the 640-nm channel (20 acquisitions per plane), and without horizontal scanning for the 488-nm channel. For TRAP2-Ai14 brains, the 640-nm channel was imaged at 20% laser power without the contrast adaptive algorithm.

### Model Training

Light sheet images of fluorescently labeled axons were segmented using the 3D U-net based machine learning pipeline TRAILMAP (Friedmann et al., 2020). We trained new models for segmentation of cortical axons using inference learning as described in the TRAILMAP pipeline (https://github.com/AlbertPun/TRAILMAP). Briefly, 120x120 pixel stacks of 100 images were selected for training, and axons and artifacts (pixels that could be mistakenly interpreted as axons) were hand-labeled in 3 or 4 images from each stack. 1-pixel-wide edges of axons were labeled using python to be given less weight in training of the 3D convolutional network, which accounts for slight variability in human annotation patterns.

Models 1, 2, and 3 were designed for axon segmentation in regions with delineated, marginally distinguished, and indistinct axons, respectively. Beginning with the published TRAILMAP model weights, trainings were iteratively performed. Model 2 was trained first for 7 sessions and represents our best-trained single model for use across the entire brain. From this, Model 1 for use in superficial regions was trained for 2 sessions and Model 3 was trained for 4 sessions for use in deep regions.

Each training session used between 6 and 20 hand-labeled image stacks, of which approximately 10-20% were set aside for use in model validation. Training was performed using between 20 and 120 steps per epoch and between 5 and 20 epochs per session. The best model from each session was selected by plotting loss in validation data across epoch, which was minimized to prevent overtraining. The weaknesses of the best model from each session were assessed to guide selection of training data for the next session. Sessions were continued until all visible axons in the desired regions were reliably detected by the model.

The TRAP2-Ai14 model was trained in the same manner. Initial weights were derived from model that had been trained on fos^+^ cells. From this, 6 sessions of training with 20 steps per epoch and between 4 and 150 epochs per session were performed to generate the TRAP2-Ai14 model.

### Axon Model Validation

We validated our analysis pipeline including image segmentation, scaling and axon thinning by selecting 120x120 pixel stacks of 100 images from regions assigned to the ‘delineated’, ‘moderately distinguished’, and ‘indistinct’ groups in brains that were not included in initial training or validation data sets. Two human experts annotated 2–4 images from each stack, marking pixels likely to contain axons. In stacks with difficult-to-distinguish axons, such as deep regions, experts used larger brush strokes to label broader groups of pixels likely to contain axons (Figure S1B,C). One-pixel edges were then added to human annotations in python.

The raw image stacks were segmented as explained below. In brief, each stack was segmented with Models 1, 2 and 3 using TRAILMAP and scaled in ImageJ to a 10um space. Scaled images were skeletonized in python at 8 different thresholds for binarization, combined in MATLAB, and pixel values were adjusted in ImageJ. Using the same quantification threshold as used in all analyses (number of skeletonized pixels with intensity above 64 divided by total number of pixels in the region), labeling density was calculated in human-annotated “axon-positive” pixels (LP) and “axon-negative” pixels (LN), excluding edges to account for slight variability in human labels. Distinction score was calculated from these values as (LP-LN)/LN+(1-LP). While direct true/false positive and true/false negative rates cannot be calculated due to the axon thinning process, which reduces the number of labeled pixels, LP correlates with the true positive rate, LN correlates with the false positive rate, and 1-LP correlates with the false negative rate. Thus, LP-LN in the numerator will increase distinction score when better separation is obtained between true and false positives, and LN and 1-LP in the denominator will decrease the distinction score if there is a higher rate of false positives or false negatives.

Alternate model combination methods were performed in FIJI. Maximum probability projections between all models were calculated by generating images with maximum pixel values across all models for each cube then processing the cubes for validation as above. Summation of probability maps between models was calculated by adding probability calculated by each model, then dividing by 3 prior to processing the cubes for validation as above. Distinction scores from alternate approaches were compared using repeated measures one-way ANOVA with Dunnett’s multiple comparisons test, comparing individual model selection (DeepTraCE) with each other method. Statistical comparisons of distinguishment scores were performed using GraphPad PRISM.

### TRAP2-Ai14 Model Validation

To validate that cell counts produced by DeepCOUNT were accurate, we selected 120x120x100 image cubes from several brain regions across multiple brains not included in the training dataset. Cubes were scaled and processed in the same manner as whole-brain data to produce a raw cell count from each cube. Two human experts then manually counted cells in each cube. Average counts from the two human experts were then compared to counts produced by DeepCOUNT as shown in Figure 4.

### DeepTraCE Analysis Pipeline

#### Image Segmentation & Registration

Whole-brain image stacks from the 640nm channel from each brain were segmented three times in TRAILMAP, once each using models 1, 2, and 3. Following segmentation, the 488nm autofluorescence channel and axon segmentations from each model were converted to 8-bit and scaled to 10um resolution in FIJI with scaling values of 0.40625 in the x and y directions and 0.3 in the z direction using a bilinear interpolation algorithm. To improve image registration, each scaled 488nm image was manually rotated in the x, y, and z planes using the TransformJ ImageJ plugin (https://imagescience.org/meijering/software/transformj/) such that the midline blood vessels visible were all visible in the same z plane. The same manual rotation parameters were then applied to the three scaled model segmentation images from the corresponding brain.

The scaled and manually rotated 488nm autofluorescence channel was registered using elastix to the Gubra Lab LSFM atlas average template, which has annotations based on the Allen CCF^87–89^. The same transformation was applied to the scaled and manually rotated model segmentations using transformix. Image registration quality was manually verified by overlaying the atlas image and the registered 488nm channels in ImageJ. Following segmentation and registration of each model, the transformix images were converted to 8-bit .tif format in ImageJ. When combining multiple probability maps, it is important that minimum and maximum pixel values are comparable between images. Model 1 produced slightly lower maximum probability values compared to the other two models. We corrected for this by brightening segmentations from Model 1 by 34% to match pixel values obtained by the other models prior to combination and thinning. We provide a unified python pipeline for automating these steps in the supporting software repository.

#### Model Combination & Thinning

Converted segmentations from the three models were then combined by generating a new image in which pixel values in each region were extracted from the model with best performance in that region (Figure 2B). The same regional model assignments for combination were used for all brains in the data set and are provided in Table 2. Following model combination, axon segmentations were thinned (or ‘skeletonized’) as described in the TRAILMAP pipeline^29^. In brief, images were binarized at 8 different thresholds from 20 to 90% of the maximum intensity value using python. Skeletons were combined in MATLAB by summing values from each skeleton. Small objects unlikely to be axons were removed by calculating connected components within the combined skeleton and removing objects less than 90 voxels in size. Combined skeletons with small objects removed were optimized for visualization and quantification using an ImageJ macro that multiplied each pixel value by 17.

#### Axon Quantification

Regional axon innervation was quantified in MATLAB (Mathworks) by counting the number of skeletonized pixels in each brain region above a threshold (64), then dividing this pixel count by the total number of pixels in a region. This regional pixel count was then divided by the total number of labeled pixels across the brain to normalize for differences in total fluorescence and viral expression. Regions were defined by a collapsed version of the LSFM atlas in which maximum granularity was balanced with the need to account for slight differences in registration which would lead to inaccurate quantification of small brain regions. This atlas was cropped on the anterior and posterior ends to match the amount of tissue visible in our data. Fiber tracts, ventricular systems, cerebellum, and olfactory bulb were excluded from analysis.

All statistical comparisons were performed in MATLAB and GraphPad Prism v9. For comparison of regional axon labeling between cell types, Two-way ANOVA with Tukey’s multiple comparison correction was performed on all brain regions. Anterior-posterior axon distributions within regions were calculated in MATLAB by binning the whole-brain image into 100um voxels and calculating the percentage of segmented pixels within each voxel. Voxels falling within a given region were summed across the medial-lateral and dorsal-ventral axis and normalized for total fluorescence as above. The averaged summation of axon counts from a given cell class was then averaged and plotted along with the standard error of the mean.

#### Axon Visualization

To visualize axons as shown in Figures 1 and 3, Z-projections of raw light sheet data were created in FIJI by scaling images to a 4.0625um space, virtually reslicing images in the coronal plane, and performing maximum intensity z-projections of 100um depth followed by local contrast enhancement. Axon segmentations were created by overlaying skeletonized and registered axon segmentations from each sample of a cell type in a slightly different color and virtually reslicing in the coronal plane. 3D projections were created in Imaris using a representative sample from each cell type. Dotogram overlays (Figures 2 and 4) were created using MATLAB. Images were binned into 100um voxels and the percentage of segmented pixels within each voxel was calculated. Area of visible dot in the overlay corresponds with the averaged labeling intensity within a voxel across a condition. Outer dots represent the cell type with the highest labeling intensity within that voxel.

#### Computational Analyses

Hierarchical clustering of regional axon quantifications was performed as described in Kebschull et al., 2020^36^.

### DeepCOUNT Analysis Pipeline

#### Image Segmentation & Registration

Whole-brain image stacks from the 640nm channel from each brain were segmented in TRAILMAP using the TRAP2-Ai14 trained model. Images were registered using the 488nm autofluorescence channel as described above. Transformed images were converted to 8-bit in ImageJ.

#### 3D Maxima Detection & Single-Pixel Reduction

MATLAB was used to identify 3D maxima of the transformed probability map. Connected component analysis was then used to reduce any maxima that consisted of multiple pixels into a single pixel per cell.

#### Cell Quantification

Regional TRAP^+^ cell density was quantified in MATLAB (Mathworks) by counting the number of labeled pixels (i.e. cells) in each brain region, then dividing this pixel count by the total number of pixels in a region. This regional pixel count was then divided by the total number of detected cells across the brain to normalize for differences in tamoxifen-induced recombination. Regions were defined in the same way as described for the DeepTraCE pipeline.

### Functional Network Construction

Networks were constructed by thresholding inter-regional correlations (Pearson’s r≥0.9, *P*<0.05) for PMA and control groups. The nodes represent brain region and the correlatio^≥^ns above threshold were considered connections. Cytoscape software was used to visualize networks. Node size was proportional to degree (number of connections).

### 4-Hydroxytamoxifen Preparation

4-hydroxytamoxifen (4-OHT; Sigma, Cat# H6278) was dissolved at 20□mg mL^-1^ in ethanol by shaking at 37°C for 15□min and was then aliquoted and stored at –20°C for up to several weeks. Before use, 4-OHT was redissolved in ethanol by shaking at 37°C for 15□min, a 1:4 mixture of castor oil:sunflower seed oil (Sigma, Cat #s 259853 and S5007) was added to give a final concentration of 10□mg mL^-1^ 4-OHT, and the ethanol was evaporated by vacuum under centrifugation. The final 10□mg mL^-1^ 4-OHT solutions were always used on the day they were prepared. All injections were delivered intraperitoneally (i.p.).

### Behavioral assays

For PMA experiments, mice were placed in an operant chamber with a shock floor. The quarter of the floor furthest from the door of the chamber was covered in a white plexiglass platform. Two odor pods with novel scents (vanilla, almond, coconut, or peanut butter) were placed underneath the part of the shock floor not covered by the platform to promote exploration.

For TRAP2 experiments, on training day, mice received 3 baseline tones (30s, 4000Hz, 75dB) followed by 9 tone-shock pairings (0.13mA shock, 2s, co-terminating), where mice could learn to avoid the shock by entering the safety platform. Tones were separated by a random interval between 80 and 150 seconds. The next day, mice received 6 tones with no shock. Mice were injected with 4-OHT solution immediately following the retrieval session. Non-shock control animals were placed in the operant chamber with no platform for five minutes on day 1, and on day 2 were placed in the operant chamber for five minutes and injected with 4-OHT solution immediately after this session.

For fiber photometry recordings during PMA, mice received 3 baseline tones on day 1, followed by 12 tone-shock pairings on day 1 and 16 tone-shock pairings on days 2, and 3. For EZM experiments, mice were placed on a custom-built elevated zero maze 24 inches in diameter for 15 minutes. Both assays were recorded using a Point Grey Chameleon3 USB camera (Teledyne FLIR).

### Fiber Photometry

Mice were habituated to the operant chamber and optic fiber for at least two days prior to recording. A TDT RZ10x processor in combination with the TDT Synapse software was used to simultaneously record the 405nm isosbestic channel and the 465nm signal channel during behavior. For each mouse, the light output was adjusted such that the 465nm and 405nm channel produced a signal of approximately 80mV as reported by the Synapse software.

### Fiber Photometry Analysis

Point-tracking of PMA videos were performed in DeepLabCut^90^ and behavior was analyzed using BehaviorDEPOT^91^. Elevated Zero Maze behavioral epochs were annotated manually. Fiber photometry analysis was performed in MATLAB in a modified version of an example provided by TDT. To align fiber photometry and behavioral data, TTL pulses marking the beginning and end of each tone were aligned between the fiber photometry signal and video frames. A lookup table was generated using linear interpolation between each TTL pulse to identify which behavior frame lines up with each photometry frame. For alignment of EZM fiber photometry data, a silent TTL pulse was generated every 30 seconds to be used for alignment in the same manner.

Fiber photometry signal was down-sampled by a factor of 10 prior to analysis. To account for potential movement artifacts and bleaching, the 405nm isosbestic control channel was fit to the 465nm signal using the polyfit function, and this curve was then subtracted from the 465nm signal. Z scores of this signal were calculated using a baseline period of -10 to 0 seconds relative to the tone for tone-aligned responses (i.e. tone and shock responses) and -20 to -15 seconds relative to epoch onset for all other behaviors (i.e. platform entries and exits, closed arm entries and exits, head dips). The average of all traces for an individual animal was calculated and used for analysis. To generate plots, each animal’s average trace was smoothed by averaging values from every 0.5 seconds (for time-locked tone and shock responses) or using a moving average of 0.5 seconds (for all other traces), and the mean ± SEM of smoothed traces across animals was displayed. To generate shock-normalized AUC values (provided in Figure S8), the maximum shock response from an individual was calculated from the averaged shock trace. All values in a trace were then divided by this value, and the resulting trace was used to calculate the normalized AUC.

## CODE AVAILABILITY

Code, instructions and sample data available at https://github.com/DeNardoLab/DeepTraCE and https://github.com/jcouto/DeepTraCE/tree/gui

## ACKNOWLEDGEMENTS

We thank Justus Kebschull and Xiaoyin Chen for helpful comments on the manuscript. This work was funded by K01MH116264 (L.A.D.), 1R01MH127214-01A1 (L.A.D.), a Whitehall Foundation Research Grant (L.A.D), a Klingenstein-Simons Foundation Grant (L.A.D.), a NARSAD Young Investigator Award (L.A.D.), a Fay/Frank Seed Grant (L.A.D.), an NSERC Postgraduate Fellowship (C.B.K.), T32MH073526-14 (M.W.G), and the UCLA Medical Scientist Training Program (M.W.G.).

## AUTHOR CONTRIBUTIONS

C.B.K, M.W.G. and L.A.D designed the research; C.B.K. performed the stereotaxic AAV injections and PMA fiber photometry; C.B.K. and B.J. performed tissue clearing; C.B.K., M.W.G., and B.J. performed imaging and data analysis; D.F. and M.W.G. wrote DeepTraCE and DeepCOUNT code; J.C. ported the pipeline to python and wrote the jupyter examples; C.B.K, R.C., and A.S.E. performed histology and image analysis, A.S.E. and M.W.G performed EZM fiber photometry; C.B.K, M.W.G. and L.A.D. wrote the manuscript.

