## Supplemental Figures for "Brain-wide projections and differential encoding of prefrontal neuronal classes underlying learned and innate threat avoidance"

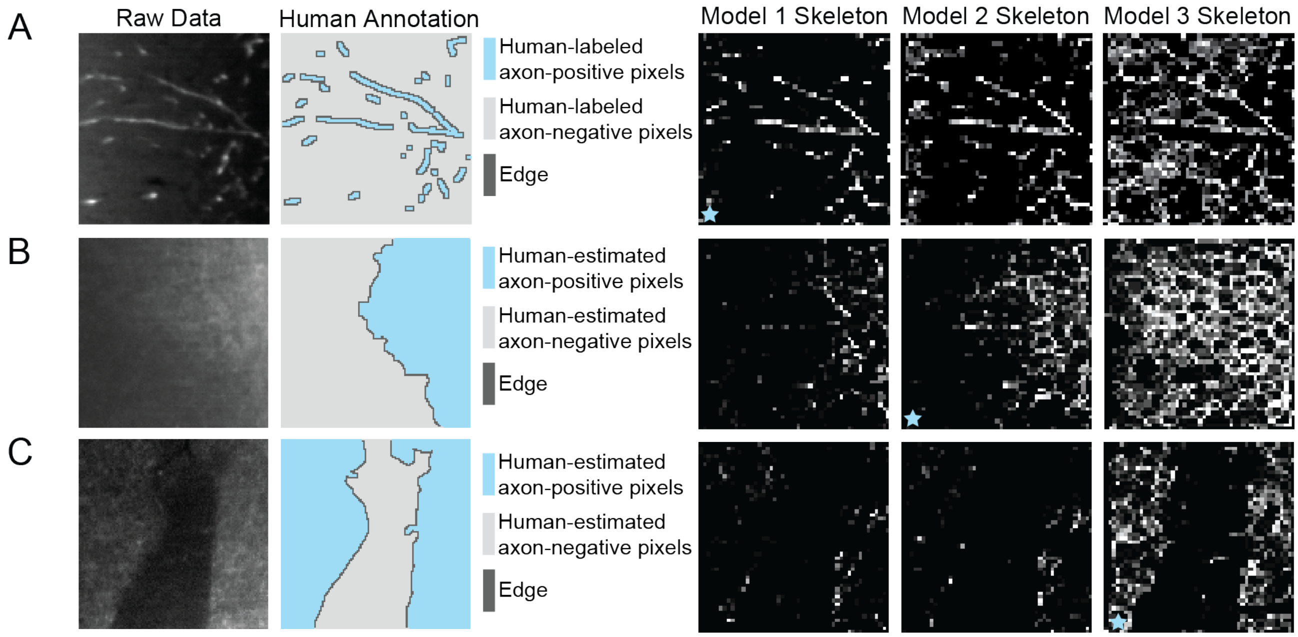


**Supplemental Figure S1. Validation of DeepTraCE Pipeline. Related to Figure 1.**

(A-C) Single 500x500um planes of raw axon data, human annotation, and skeletons from Models 1, 2, and 3 in images of axons from each group. Blue stars indicate model fine-tuned for segmentation in corresponding brain region.

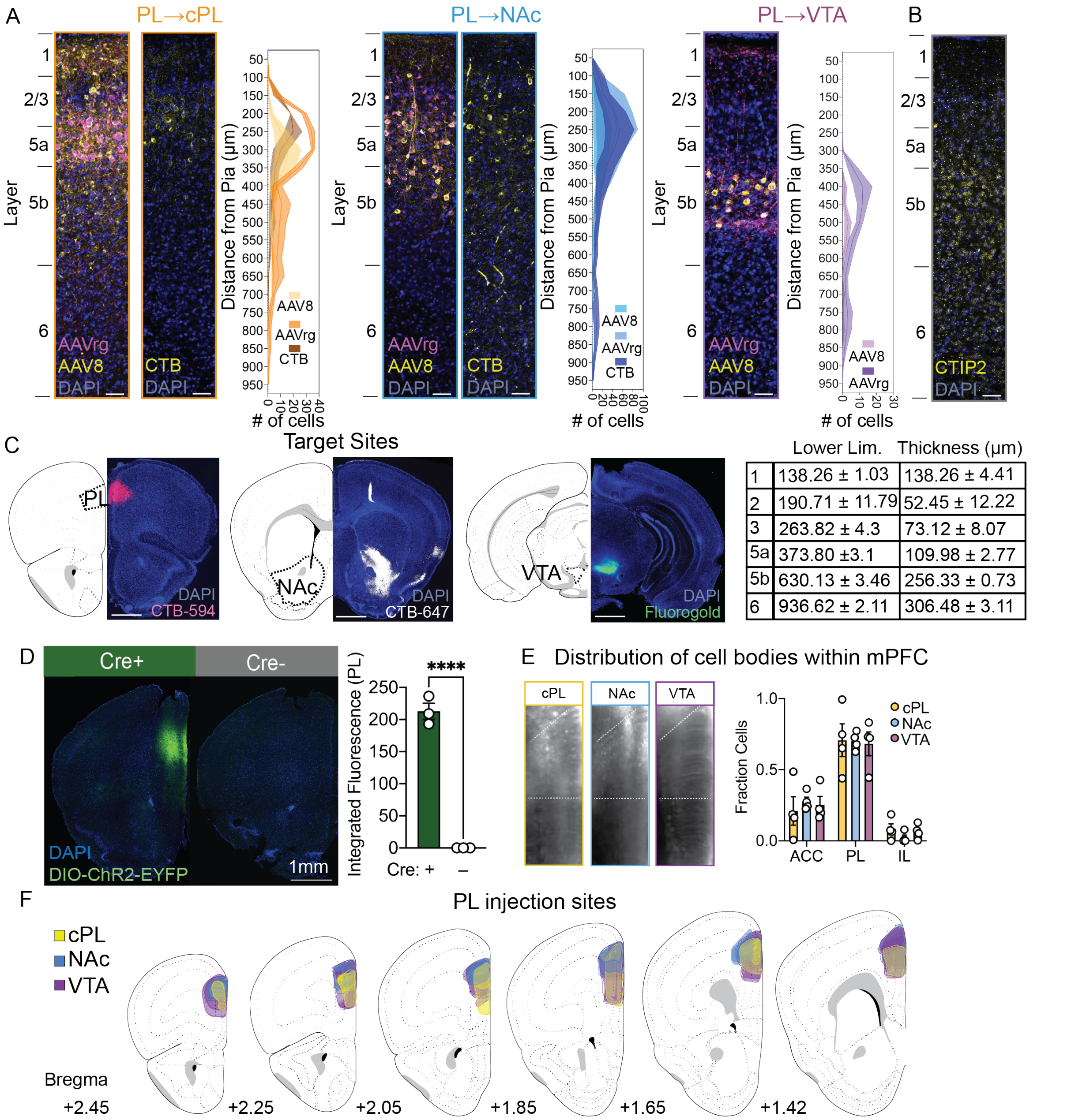


| | Lower Lim. | Thickness (µm) |
|---|---|---|
| 1 | 138.26 ± 1.03 | 138.26 ± 4.41 |
| 2 | 190.71 ± 11.79 | 52.45 ± 12.22 |
| 3 | 263.82 ± 4.3 | 73.12 ± 8.07 |
| 5a | 373.80 ±3.1 | 109.98 ± 2.77 |
| 5b | 630.13 ± 3.46 | 256.33 ± 0.73 |
| 6 | 936.62 ± 2.11 | 306.48 ± 3.11 |

**Supplemental Figure 2. Validation of Cre-Dependent Virus and Injection Site Mapping. Related to Figure 2.**

(A) Layer distributions of prelimbic neurons retrogradely labeled via injections of AAVrg, AAVrg+AAV8, CTB-594, CTB-647, and AAVrg-EGFP in cPL, NAc, and VTA, respectively (cPL: $F_{tracer}$=9.69, *P*=0.05; $F_{bin}$=52.14, *P*<0.0001, n=3/group; NAc: $F_{tracer}$=3.71, *P*=0.62; $F_{bin}$ =30.48, *P*=0.07, n=3/group; VTA: $F_{tracer}$=8.52, *P*=0.0015; $F_{bin}$ =5.84, *P*=0.02, n=3–6/group 2-way ANOVA). Scale bars, 100um.

(B) Representative stain of layer marker CTIP2 in PL (top) and table of layer lower limits and thickness (bottom). Scale bars, 100um.

(C) Representative images of target sites (cPL, NAc and VTA) visualized with Cholera Toxin Subunit B and Fluorogold. Scale bars, 500um.

(D) Representative Coronal Sections sections of brains injected with AAVrg-Cre and AAV-DIO-ChR2-EYPF (left) or AAV-DIO-ChR2-EFYP in the absence of Cre and quantification of fluorescence in PL.

(E) Raw fluorescence images from LSFM showing mPFC and quantifications of cell body numbers within ACC, PL and IL.

(F) Maps of starter cells in mPFC for each cell type

Error bars, S.E.M., ****$P$<0.001, Student's t-test.

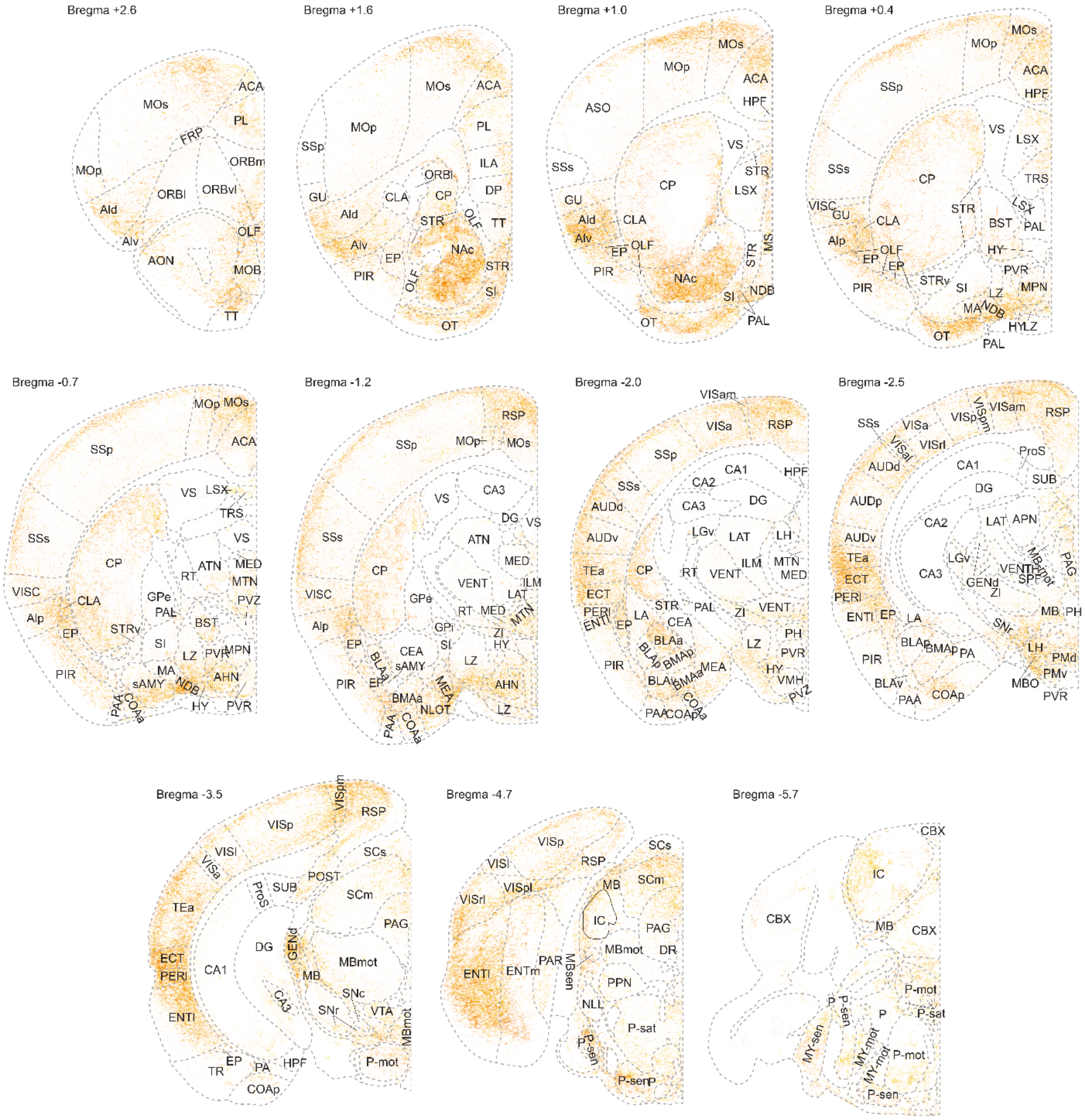


**Supplemental Figure S3. Whole-brain PL–cPL collateralization patterns. Related to Figure 3.**

Eleven fully annotated 10um coronal optical sections of skeletonized images registered to the standardized brain atlas. Each image represents 4 overlaid PL–cPL brains. Sections from Bregma values +0.7, -2.0, and -3.5 are the same as shown in Figure 3.

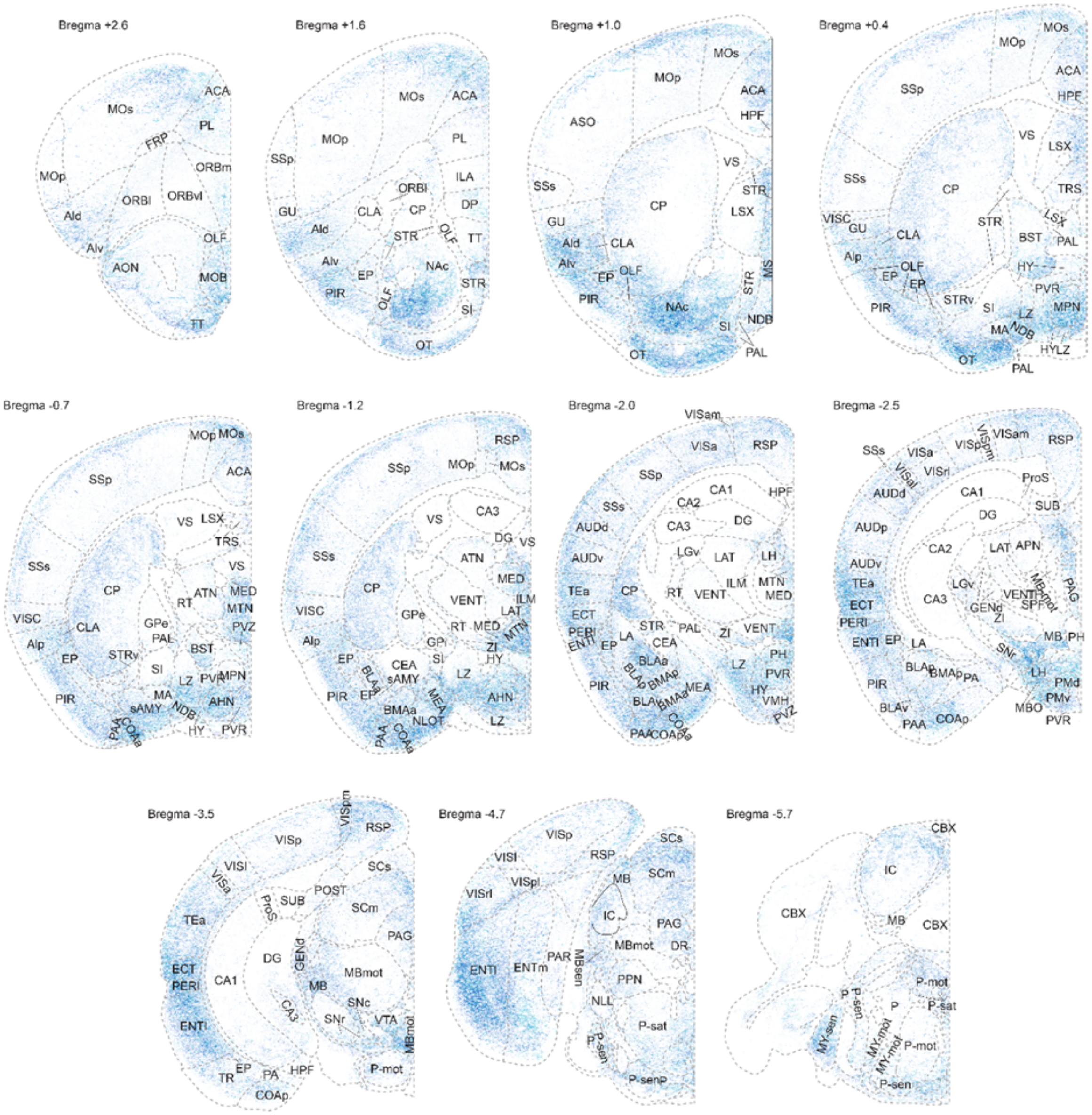


**Supplemental Figure S4. Whole-brain PL–NAc collateralization patterns. Related to Figure 3.**

Eleven fully annotated 10um coronal optical sections of skeletonized images registered to the standardized brain atlas. Each image represents 4 overlaid PL–NAc brains. Sections from Bregma values +0.7, -2.0, and -3.5 are the same as shown in Figure 3.

.

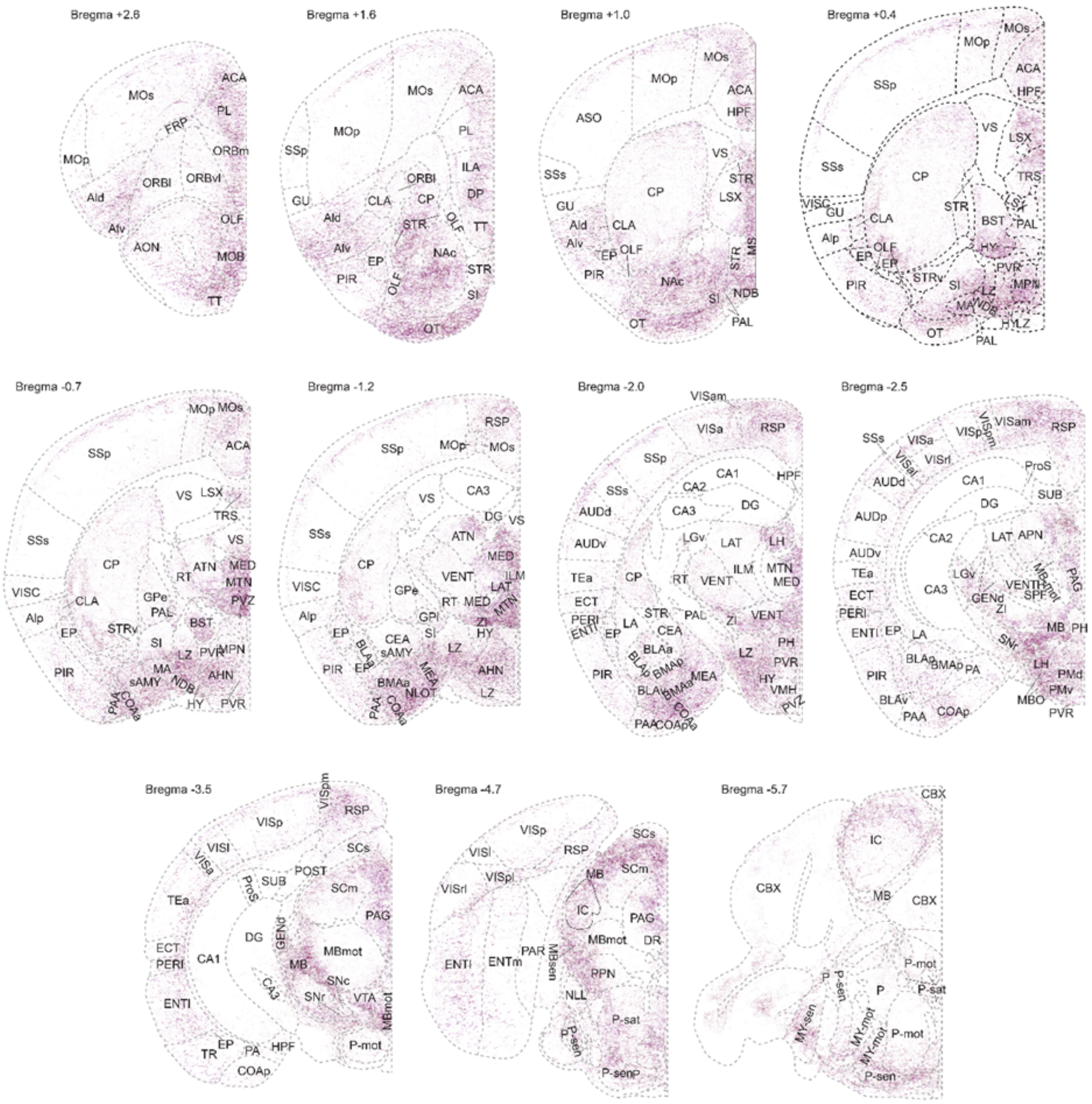


**Supplemental Figure S5. Whole-brain PL–VTA collateralization patterns. Related to Figure 3.**

Eleven fully annotated 10um coronal optical sections of skeletonized images registered to the standardized brain atlas. Each image represents 4 overlaid PL–VTA brains. Sections from Bregma values +0.7, -2.0, and -3.5 are the same as shown in Figure 2.

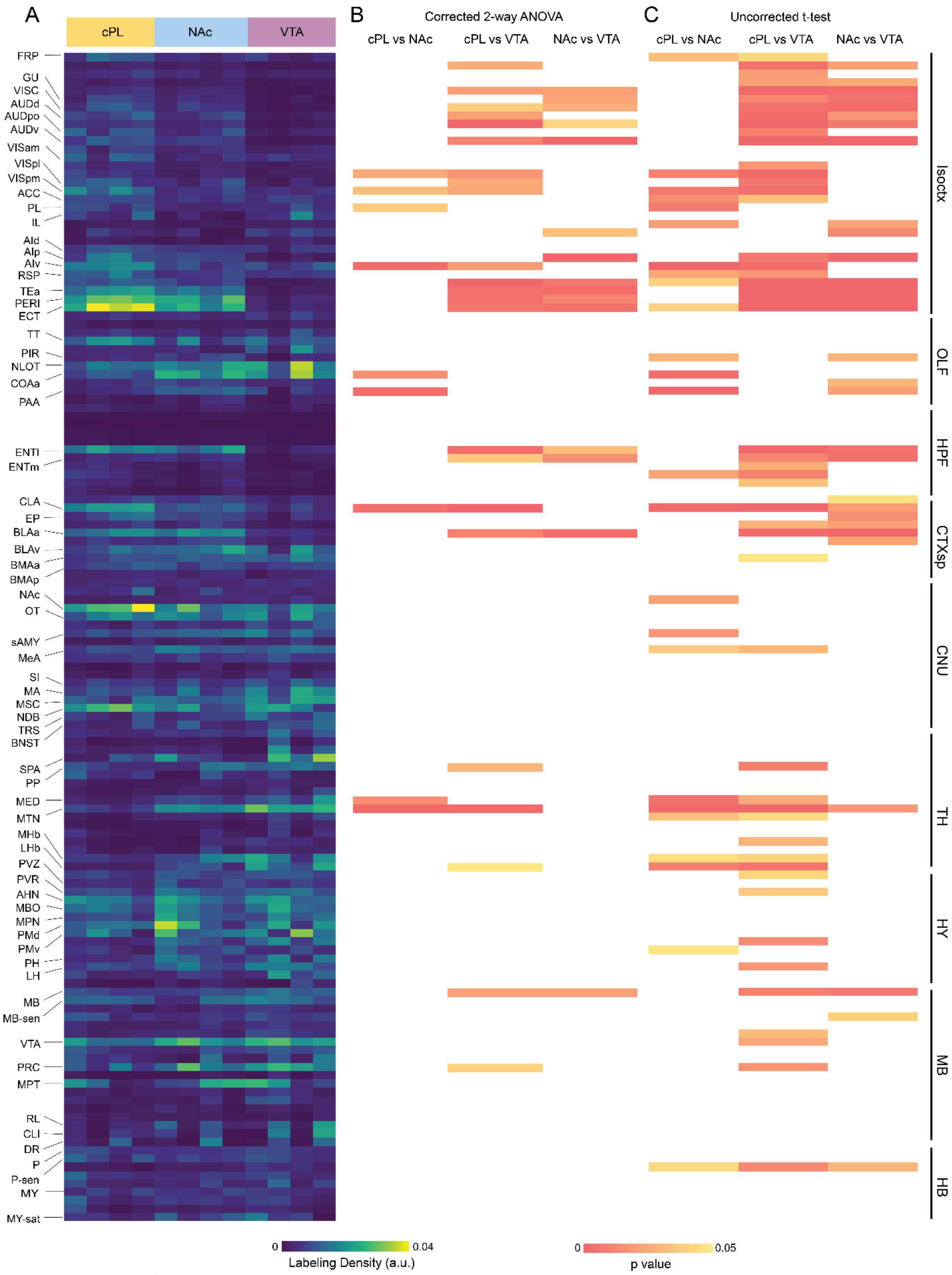


**Supplemental Figure S6. Whole-brain collateralization data and statistics ordered by Allen Brain Atlas ontology. Related to Figure 3. See also Table 2.**

(A) Heatmap from Figure 3 ordered by Allen Brain Atlas ontology.
(B) Heatmap of p values from 2-way ANOVA with Tukey's multiple comparison correction for all brain regions.
(C) Heatmap of uncorrected p values from unpaired t-tests between cPL and NAc, cPL and VTA, and NAc and VTA.

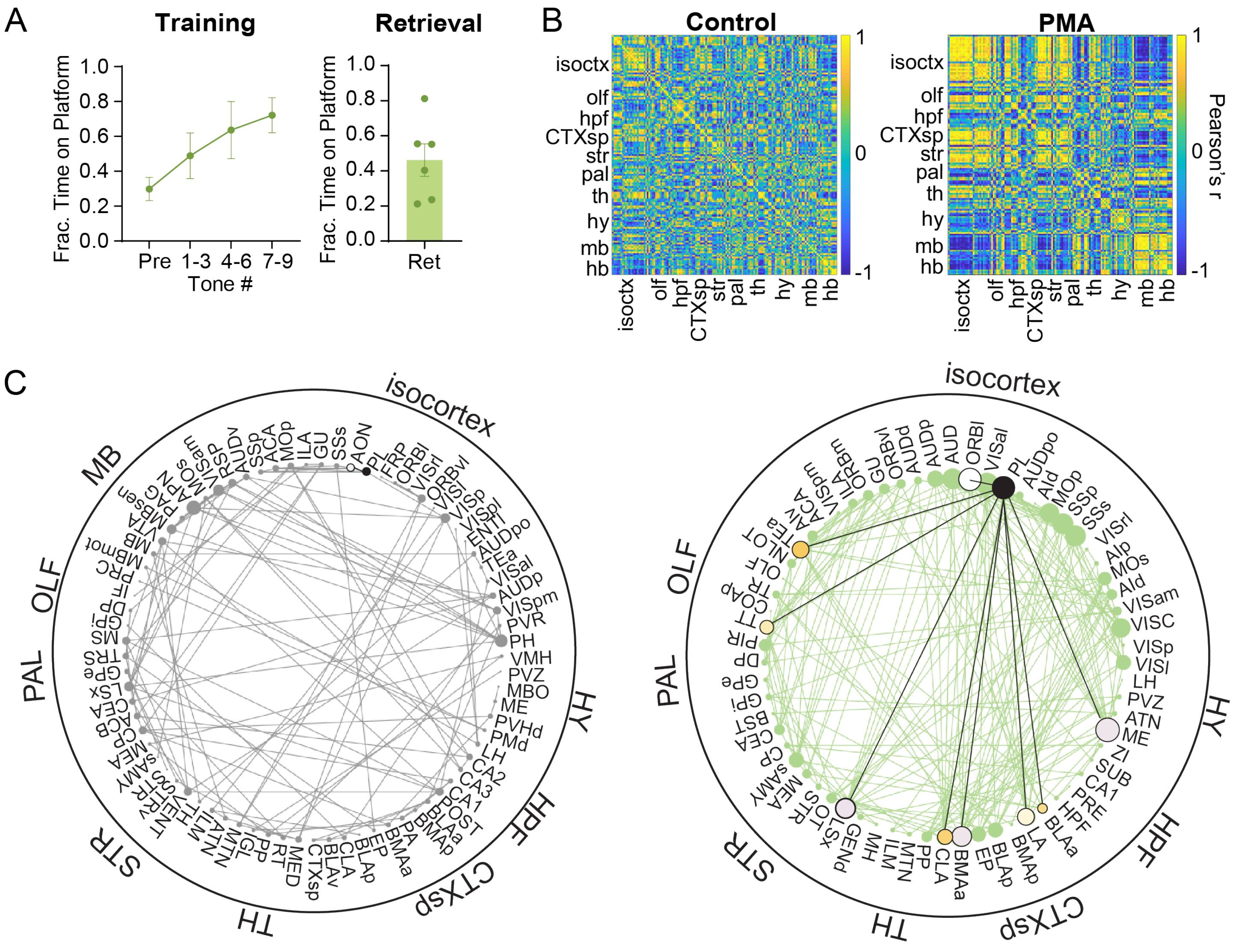


**Supplemental Figure S7. Whole-brain TRAP during PMA**

(A) Behavioral data for PMA training and retrieval (PMA train: $F_{time}$=22.78, $P$=0.02, n=7mice, 2-way ANOVA).

(B) Inter-regional correlations based on whole-brain TRAP. Colormap represents Pearson's r value.

(C) Detailed network diagrams built from thresholded whole-brain correlations (Pearson's r≤0.9, $P$<0.05) shown in B.

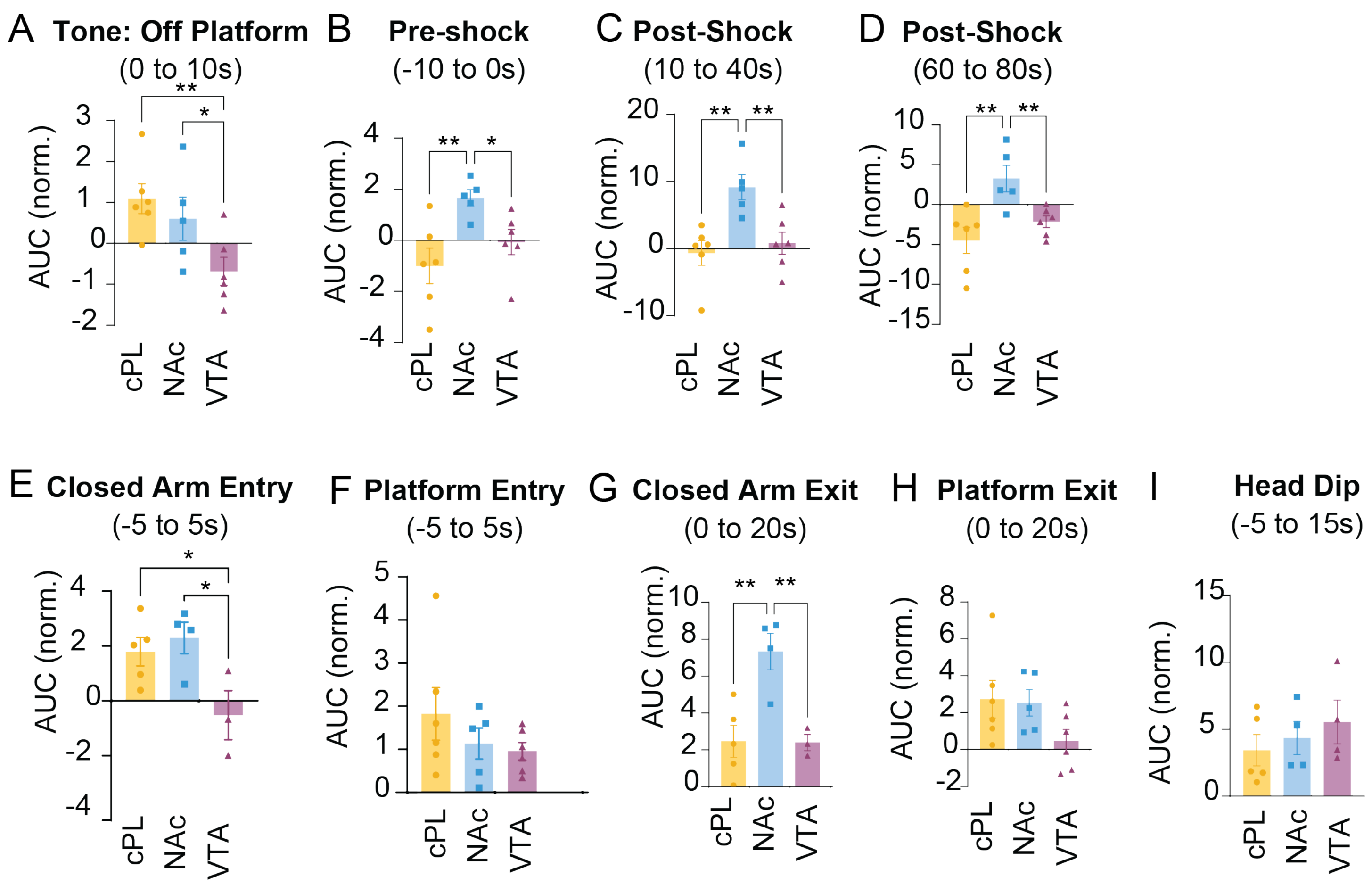


**Supplemental Figure S8. Shock-normalized activity (related to figures 5 and 6)**
(A) Normalized tone responses when mice were off the safety platform (*F*=5.37, *P*=0.02, one-way ANOVA with Benjamini, Krieger and Yekutieli post-hoc test).
(B) Normalized pre-shock activity (*F*=5.76, *P*=0.02, one-way ANOVA with Benjamini, Krieger and Yekutieli post-hoc test).
(C) Normalized post-shock activity (*F*=8.28, *P*=0.004, one-way ANOVA with Benjamini, Krieger and Yekutieli post-hoc test).
(D) Normalized post-shock activity (*F*=7.8, *P*=0.005, one-way ANOVA with Benjamini, Krieger and Yekutieli post-hoc test).
(E) Normalized closed arm entry activity (*F*=4.8, *P*=0.038, one-way ANOVA with Benjamini, Krieger and Yekutieli post-hoc test).
(F) Normalized platform entry activity (*F*=1.16, *P*=0.34, one-way ANOVA with Benjamini, Krieger and Yekutieli post-hoc test).
(G) Normalized closed arm exit activity (*F*=10.23, *P*=0.005, one-way ANOVA with Benjamini, Krieger and Yekutieli post-hoc test).
(H) Normalized platform exit activity (*F*=2.42, *P*=0.12, one-way ANOVA with Benjamini, Krieger and Yekutieli post-hoc test).
(H) Normalized platform exit activity (*F*=2.42, *P*=0.12, one-way ANOVA with Benjamini, Krieger and Yekutieli post-hoc test).
(I) Normalized head dip activity (*F*=0.64, *P*=0.55, one-way ANOVA with Benjamini, Krieger and Yekutieli post-hoc test).
*$P<0.05$, **$P<0.01$
