## Supplementary material for "Brain-wide projections and differential encoding of prefrontal neuronal classes underlying learned and innate threat avoidance": Table 1

| | |
|---|---|
| ACC | Anterior Cingulate Cortex |
| AH | Anterior Hypothalamic Area |
| AHN | Anterior Hypothalamic Nucleus |
| AId | Agranular Insula, dorsal part |
| AIp | Agranular Insula, posterior part |
| AIv | Agranular Insula, ventral part |
| ATN | Anterior Group of the dorsal thalamus |
| AUDd | Dorsal auditory area |
| AUDp | Primary auditory area |
| AUDv | Ventral auditory area |
| BLA | Basolateral amygdala |
| BLAa | Basolateral amygdala, anterior part |
| BLAv | Basolateral amygdala, ventral part |
| BNST | Bed nucleus of the stria terminalis |
| CLA | Claustrum |
| cPL | Contralateral prelimbic area |
| CNU | Cerebral nuclei |
| COA | Cortical amygdalar area |
| COAa | Cortical amygdalar area, anterior part |
| COAp | Cortical amygdalar area, posterior part |
| CP | Caudate putamen |
| CTXsp | Cortical subplate |
| DP | Dorsal peduncular area |
| ECT | Ectorhinal cortex |
| ENTl | Entorhinal area, lateral part |
| ENTm | Entorhinal area, medial part |
| EP | Endopiriform nucleus |
| GENd | Geniculate group, dorsal thalamus |
| GU | Gustatory areas |
| HB | Hindbrain |
| HPF | Hippocampal formation |
| IC | Inferior colliculus |
| IGL | Intergeniculat leaflet of the lateral geniculate complex |
| IL | Infralimbic cortex |
| ILM | Intralaminar nucleus of the dorsal thalamus |
| Isoctx | Isocortex |
| LAT | Lateral geniculate nucleus |
| LGv | Ventral part of the lateral geniculate nucleus |
| LH | Lateral hypothalamus |
| LHb | Lateral habenula |
| MA | Magnocellular Nucleus |
| MB | Midbrain |
| MB-un | Midbrain-unlabeled |
| MBO | Mammillary body |
| MeA | Medial amygdala |
| MED | Medial group of the dorsal thalamus |
| MHb | Medial habenula |
| MOp | Primary motor area |
| MOs | Secondary motor area |
| mPFC | medial prefrontal cortex |
| MPN | Medial preoptic nucleus |
| MTN | Midline group of the dorsal thalamus |
| MY-mot | Medulla, motor related |
| MY-sen | Medulla, sensory related |
| NAc | Nucleus accumbens |
| NDB | Nucleus of the diagonal band |
| NLOT | Nucleus of the lateral olfactory tract |
| OLF | Olfactory tract |
| ORBm | Orbital area, medial part |
| OT | Olfactory tubercle |
| P-mot | Pons, motor related |
| P-Sat | Pons, behavioral state related |
| P-Sen | Pons, sensory related |
| PAA | Piriform-amygdalar area |
| PAG | Periaqueductal grey |
| PERI | Perirhinal area |
| PH | Posterior hypothalamic nucleus |
| PIR | Piriform area |
| PL | Prelimbic area |
| POST | Postrhinal area |
| PP | Peripeduncular nucleus |
| PPN | Pedunculopontine nucleus |
| PRC | Precommisural nucleus |
| PVHd | Paraventricular hypothalamic nucleus, descending division |
| PVR | Periventricular region |
| RE | Nucleus of reuniens |
| RSP | Retrosplenial cortex |
| RT | Reticular nucleus of the thalamus |
| sAMY | Striatum-like amygdala |
| SCm | Superior colliculus, motor related |
| SCs | Superior colliculus, senosry related |
| SNr | Substantia nigra, reticular part |
| SPA | Subparafascicular area |
| SPF | Subparafascicular nucleus |
| SSp | Primary somatosensory area |
| SSs | Secondary somatosensory area |
| SubG | Subgeniculate nucleus |
| TEa | Temporal association area |
| TH | Thalamus |
| TR | Postpiriform transition area |
| VENT | Ventral group of the dorsal thalamus |
| VISC | Visceral area |
| VISp | Primary visual area |
| VISpl | Posterolateral visual area |
| VISpm | Posteromedial visual area |
| VISrl | Rostrolateral visual area |
| VTA | Ventral tegmental area |
| ZI | Zona incerta |

**Table 1.** List of abbreviations
