## Supplementary material for "Brain-wide projections and differential encoding of prefrontal neuronal classes underlying learned and innate threat avoidance": Table 2

**Table 2.** Quantification of axon density by brain region. Related to Figures 2,3.

| Abbreviation | Allen Brain Atlas ID | Mean cPL | Mean NAc | Mean VTA | cPL Brain 1 | cPL Brain 2 | cPL Brain 3 | cPL Brain 4 | NAc Brain 1 | NAc Brain 2 | NAc Brain 3 | NAc Brain 4 | VTA Brain 1 | VTA Brain 2 | VTA Brain 3 | VTA Brain 4 | PC1 Weight | Corrected 2-way ANOVA: cPL vs NAc | Corrected 2-way ANOVA: cPL vs VTA | Corrected 2-way ANOVA: NAc vs VTA | Uncorrected p value: cPL vs NAc | Uncorrected p value: cPL vs VTA | Uncorrected p value: NAc vs VTA | Model Used |
|---|---|---|---|---|---|---|---|---|---|---|---|---|---|---|---|---|---|---|---|---|---|---|---|---|
| FRP | 'Frontal pole, cerebral cortex' | 0.01032 | 0.00481 | 0.00452 | 0.00550 | 0.01398 | 0.01145 | 0.01035 | 0.00472 | 0.00597 | 0.00237 | 0.00620 | 0.00811 | 0.00304 | 0.00172 | 0.00518 | 0.0750 | 0.0952 | 0.0972 | 0.9817 | 0.0322 | 0.0425 | 0.8619 | 1 |
| Mop | 'Primary motor area' | 0.00270 | 0.00183 | 0.00089 | 0.00226 | 0.00214 | 0.00258 | 0.00381 | 0.00126 | 0.00206 | 0.00156 | 0.00245 | 0.00067 | 0.00061 | 0.00105 | 0.00123 | 0.0229 | 0.2381 | 0.0266 | 0.0621 | 0.1111 | 0.0045 | 0.0208 | 1 |
| Mos | 'Secondary motor area' | 0.00556 | 0.00388 | 0.00251 | 0.00354 | 0.00541 | 0.00534 | 0.00795 | 0.00382 | 0.00534 | 0.00194 | 0.00442 | 0.00146 | 0.00268 | 0.00309 | 0.00281 | 0.0405 | 0.3783 | 0.0757 | 0.301 | 0.1967 | 0.0206 | 0.1393 | 1 |
| SSp | 'Primary somatosensory area' | 0.00254 | 0.00190 | 0.00076 | 0.00277 | 0.00102 | 0.00261 | 0.00375 | 0.00098 | 0.00168 | 0.00220 | 0.00275 | 0.00060 | 0.00067 | 0.00089 | 0.00089 | 0.0215 | 0.6433 | 0.0996 | 0.1073 | 0.3865 | 0.0210 | 0.0252 | 1 |
| SSs | 'Supplemental somatosensory area' | 0.00479 | 0.00343 | 0.00053 | 0.00300 | 0.00422 | 0.00544 | 0.00649 | 0.00192 | 0.00352 | 0.00397 | 0.00431 | 0.00062 | 0.00030 | 0.00039 | 0.00080 | 0.0540 | 0.3732 | 0.0207 | 0.0209 | 0.1907 | 0.0014 | 0.0017 | 1 |
| GU | 'Gustatory areas' | 0.00683 | 0.00435 | 0.00079 | 0.00219 | 0.00935 | 0.00894 | 0.00685 | 0.00271 | 0.00613 | 0.00372 | 0.00483 | 0.00097 | 0.00012 | 0.00168 | 0.00038 | 0.0814 | 0.4279 | 0.0646 | 0.0225 | 0.2158 | 0.0113 | 0.0047 | 1 |
| VISC | 'Visceral area' | 0.00900 | 0.00577 | 0.00056 | 0.00418 | 0.01124 | 0.01248 | 0.00811 | 0.00324 | 0.00819 | 0.00579 | 0.00586 | 0.00072 | 0.00010 | 0.00055 | 0.00086 | 0.1073 | 0.3588 | 0.039 | 0.0265 | 0.1766 | 0.0039 | 0.0022 | 1 |
| AUDd | 'Dorsal auditory area' | 0.00916 | 0.00652 | 0.00162 | 0.00851 | 0.00647 | 0.00846 | 0.01319 | 0.00418 | 0.00415 | 0.00753 | 0.01021 | 0.00134 | 0.00109 | 0.00136 | 0.00269 | 0.0918 | 0.4494 | 0.0222 | 0.0818 | 0.2436 | 0.0022 | 0.0176 | 1 |
| AUDp | 'Primary auditory area' | 0.00639 | 0.00440 | 0.00098 | 0.00652 | 0.00552 | 0.00597 | 0.00753 | 0.00309 | 0.00302 | 0.00505 | 0.00644 | 0.00076 | 0.00029 | 0.00109 | 0.00177 | 0.0640 | 0.1875 | 0.0002 | 0.0412 | 0.0771 | 0.0001 | 0.0082 | 1 |
| AUDpo | 'Posterior auditory area' | 0.01004 | 0.00598 | 0.00236 | 0.01315 | 0.00414 | 0.01124 | 0.01164 | 0.00384 | 0.00275 | 0.00613 | 0.01120 | 0.00233 | 0.00119 | 0.00259 | 0.00332 | 0.0887 | 0.3652 | 0.0582 | 0.2769 | 0.1902 | 0.0097 | 0.1090 | 1 |
| AUDv | 'Ventral auditory area' | 0.00967 | 0.00685 | 0.00064 | 0.00631 | 0.01255 | 0.00966 | 0.01015 | 0.00582 | 0.00769 | 0.00742 | 0.00646 | 0.00072 | 0.00018 | 0.00062 | 0.00105 | 0.1091 | 0.2182 | 0.0111 | 0.0004 | 0.0828 | 0.0004 | 0.0000 | 1 |
| VISal | 'Anterolateral visual area' | 0.00698 | 0.00444 | 0.00280 | 0.01018 | 0.00179 | 0.00790 | 0.00806 | 0.00296 | 0.00280 | 0.00551 | 0.00649 | 0.00184 | 0.00164 | 0.00326 | 0.00445 | 0.0438 | 0.4822 | 0.1953 | 0.3828 | 0.2570 | 0.0724 | 0.1975 | 1 |
| VISam | 'Anteromedial visual area' | 0.01042 | 0.00439 | 0.00541 | 0.01345 | 0.00328 | 0.01371 | 0.01123 | 0.00234 | 0.00526 | 0.00283 | 0.00714 | 0.00497 | 0.00742 | 0.00462 | 0.00461 | 0.0523 | 0.172 | 0.2481 | 0.7329 | 0.0660 | 0.0954 | 0.4668 | 1 |
| VISl | 'Lateral visual area' | 0.00532 | 0.00276 | 0.00183 | 0.00720 | 0.00250 | 0.00539 | 0.00618 | 0.00231 | 0.00197 | 0.00359 | 0.00317 | 0.00161 | 0.00087 | 0.00231 | 0.00255 | 0.0363 | 0.1594 | 0.0711 | 0.2677 | 0.0555 | 0.0180 | 0.1324 | 1 |
| VISp | 'Primary visual area' | 0.00655 | 0.00255 | 0.00152 | 0.00887 | 0.00527 | 0.00681 | 0.00526 | 0.00160 | 0.00126 | 0.00361 | 0.00371 | 0.00106 | 0.00151 | 0.00171 | 0.00179 | 0.0515 | 0.0252 | 0.0174 | 0.3816 | 0.0097 | 0.0012 | 0.1751 | 1 |
| VISpl | 'Posterolateral visual area' | 0.01018 | 0.00620 | 0.00227 | 0.00974 | 0.01249 | 0.01301 | 0.00547 | 0.00570 | 0.00196 | 0.00632 | 0.01083 | 0.00163 | 0.00059 | 0.00504 | 0.00180 | 0.0976 | 0.3215 | 0.026 | 0.237 | 0.1640 | 0.0071 | 0.1043 | 1 |
| VISpm | 'posteromedial visual area' | 0.01562 | 0.00630 | 0.00488 | 0.01909 | 0.01145 | 0.01953 | 0.01241 | 0.00397 | 0.00621 | 0.00591 | 0.00909 | 0.00429 | 0.00563 | 0.00332 | 0.00628 | 0.1124 | 0.0322 | 0.0243 | 0.536 | 0.0079 | 0.0030 | 0.2997 | 1 |
| ACC | 'Anterior cingulate area' | 0.00889 | 0.00599 | 0.00649 | 0.00911 | 0.00832 | 0.00942 | 0.00870 | 0.00424 | 0.00662 | 0.00517 | 0.00794 | 0.00502 | 0.00827 | 0.00756 | 0.00512 | 0.0288 | 0.0676 | 0.1188 | 0.905 | 0.0142 | 0.0330 | 0.6832 | 2 |
| PL | 'Prelimbic area' | 0.00833 | 0.00361 | 0.00626 | 0.00983 | 0.00911 | 0.00500 | 0.00939 | 0.00386 | 0.00286 | 0.00276 | 0.00497 | 0.00538 | 0.00474 | 0.01011 | 0.00481 | 0.0248 | 0.0367 | 0.4899 | 0.2532 | 0.0087 | 0.2706 | 0.1059 | 2 |
| IL | 'Intralimbic area' | 0.00730 | 0.00156 | 0.00871 | 0.00833 | 0.00387 | 0.00315 | 0.01387 | 0.00083 | 0.00437 | 0.00073 | 0.00031 | 0.00462 | 0.00568 | 0.01835 | 0.00621 | -0.0088 | 0.1939 | 0.9366 | 0.2132 | 0.0728 | 0.7404 | 0.0775 | 2 |
| ORBl | 'Orbital area, lateral part' | 0.00185 | 0.00101 | 0.00220 | 0.00122 | 0.00213 | 0.00168 | 0.00236 | 0.00099 | 0.00088 | 0.00126 | 0.00092 | 0.00160 | 0.00156 | 0.00317 | 0.00248 | -0.0027 | 0.081 | 0.7356 | 0.1016 | 0.0206 | 0.4717 | 0.0236 | 1 |
| ORBm | 'Orbital area, medial part' | 0.00432 | 0.00196 | 0.00734 | 0.00086 | 0.00794 | 0.00508 | 0.00342 | 0.00176 | 0.00550 | 0.00018 | 0.00041 | 0.00792 | 0.00531 | 0.00958 | 0.00653 | -0.0257 | 0.4838 | 0.2851 | 0.0332 | 0.2668 | 0.1350 | 0.0128 | 1 |
| ORBvl | 'Orbital area, ventrolateral part' | 0.00152 | 0.00066 | 0.00234 | 0.00074 | 0.00237 | 0.00175 | 0.00121 | 0.00021 | 0.00141 | 0.00017 | 0.00086 | 0.00152 | 0.00154 | 0.00505 | 0.00126 | -0.0041 | 0.231 | 0.6965 | 0.2998 | 0.1115 | 0.4275 | 0.1276 | 1 |
| AId | 'Agranular insular area, dorsal part' | 0.00911 | 0.00647 | 0.00565 | 0.00561 | 0.01027 | 0.01129 | 0.00925 | 0.00657 | 0.00598 | 0.00692 | 0.00642 | 0.00661 | 0.00183 | 0.00714 | 0.00703 | 0.0490 | 0.2307 | 0.2082 | 0.8131 | 0.0800 | 0.1001 | 0.5490 | 1 |
| AIp | 'Agranular insular area, posterior part' | 0.01245 | 0.00844 | 0.00136 | 0.00639 | 0.01595 | 0.01851 | 0.00894 | 0.00798 | 0.00965 | 0.00818 | 0.00795 | 0.00144 | 0.00028 | 0.00250 | 0.00121 | 0.1394 | 0.4458 | 0.0579 | 0.0001 | 0.2141 | 0.0086 | 0.0000 | 1 |
| AIv | 'Agranular insular area, ventral part' | 0.01792 | 0.00801 | 0.00869 | 0.01654 | 0.01733 | 0.01863 | 0.01918 | 0.00803 | 0.01064 | 0.00506 | 0.00831 | 0.01009 | 0.00459 | 0.00738 | 0.01269 | 0.1059 | 0.0021 | 0.0195 | 0.9441 | 0.0003 | 0.0024 | 0.7573 | 1 |
| RSP | 'Retrosplenial area' | 0.01266 | 0.00652 | 0.00656 | 0.01183 | 0.01119 | 0.01794 | 0.00969 | 0.00442 | 0.00631 | 0.00630 | 0.00907 | 0.00556 | 0.00702 | 0.00622 | 0.00742 | 0.0702 | 0.0735 | 0.0812 | 0.9995 | 0.0243 | 0.0169 | 0.9769 | 1 |
| VISrl | 'Rostrolateral visual area' | 0.01076 | 0.00761 | 0.00122 | 0.01044 | 0.00882 | 0.01256 | 0.01123 | 0.00955 | 0.00592 | 0.00620 | 0.00879 | 0.00073 | 0.00115 | 0.00075 | 0.00223 | 0.1092 | 0.0893 | 0.0007 | 0.0069 | 0.0395 | 0.0000 | 0.0006 | 1 |
| TEa | 'Temporal association areas' | 0.01926 | 0.01368 | 0.00171 | 0.01369 | 0.01957 | 0.02109 | 0.02271 | 0.01250 | 0.01669 | 0.01120 | 0.01431 | 0.00205 | 0.00051 | 0.00222 | 0.00207 | 0.2168 | 0.1263 | 0.0048 | 0.0022 | 0.0509 | 0.0001 | 0.0001 | 1 |
| PERI | 'Perirhinal area' | 0.02741 | 0.02306 | 0.00318 | 0.02034 | 0.03022 | 0.03142 | 0.02765 | 0.02414 | 0.02394 | 0.01485 | 0.02934 | 0.00337 | 0.00139 | 0.00415 | 0.00378 | 0.3069 | 0.543 | 0.0032 | 0.0121 | 0.3086 | 0.0001 | 0.0006 | 1 |
| ECT | 'Ectorhinal area' | 0.03475 | 0.02265 | 0.00307 | 0.02323 | 0.04126 | 0.03622 | 0.03829 | 0.02182 | 0.02649 | 0.01640 | 0.02591 | 0.00377 | 0.00138 | 0.00435 | 0.00277 | 0.3937 | 0.1031 | 0.0074 | 0.0048 | 0.0393 | 0.0002 | 0.0002 | 1 |
| OLF | 'Olfactory areas' | 0.00543 | 0.00473 | 0.00557 | 0.00392 | 0.00521 | 0.00729 | 0.00528 | 0.00543 | 0.00630 | 0.00283 | 0.00435 | 0.00697 | 0.00257 | 0.00841 | 0.00431 | 0.0089 | 0.7817 | 0.9951 | 0.8488 | 0.5184 | 0.9282 | 0.5981 | 1 |
| AOB | 'Accessory olfactory bulb' | 0.00145 | 0.00150 | 0.00189 | 0.00261 | 0.00083 | 0.00182 | 0.00054 | 0.00141 | 0.00143 | 0.00134 | 0.00180 | 0.00132 | 0.00113 | 0.00338 | 0.00174 | -0.0048 | 0.9953 | 0.8076 | 0.7475 | 0.9250 | 0.5490 | 0.4784 | 1 |
| AON | 'Anterior olfactory nucleus' | 0.00507 | 0.00369 | 0.00619 | 0.00399 | 0.00493 | 0.00748 | 0.00389 | 0.00267 | 0.00625 | 0.00184 | 0.00399 | 0.00709 | 0.00177 | 0.01145 | 0.00444 | 0.0020 | 0.5561 | 0.8749 | 0.5615 | 0.3183 | 0.6351 | 0.3144 | 1 |
| TT | 'Taenia tecta' | 0.01730 | 0.01019 | 0.01122 | 0.01100 | 0.02028 | 0.02117 | 0.01675 | 0.00648 | 0.01989 | 0.00452 | 0.00987 | 0.01559 | 0.00646 | 0.01230 | 0.01054 | 0.0929 | 0.281 | 0.1877 | 0.9626 | 0.1352 | 0.0882 | 0.8004 | 1 |
| DP | 'Dorsal peduncular area' | 0.00404 | 0.00327 | 0.01098 | 0.00374 | 0.00450 | 0.00410 | 0.00382 | 0.00370 | 0.00606 | 0.00169 | 0.00162 | 0.01197 | 0.00211 | 0.02059 | 0.00924 | -0.0614 | 0.7655 | 0.3051 | 0.2557 | 0.4951 | 0.1194 | 0.0995 | 2 |
| PIR | 'Piriform area' | 0.00391 | 0.00785 | 0.00318 | 0.00270 | 0.00337 | 0.00574 | 0.00380 | 0.00782 | 0.00455 | 0.00889 | 0.01017 | 0.00261 | 0.00031 | 0.00545 | 0.00435 | 0.0196 | 0.0803 | 0.8479 | 0.0663 | 0.0279 | 0.5954 | 0.0294 | 1 |
| NLOT | 'Nucleus of the lateral olfactory tract' | 0.01308 | 0.01788 | 0.02135 | 0.00846 | 0.01715 | 0.01303 | 0.01365 | 0.01838 | 0.01471 | 0.01458 | 0.02387 | 0.02413 | 0.00929 | 0.03616 | 0.01582 | -0.0444 | 0.2801 | 0.4455 | 0.8478 | 0.1388 | 0.2213 | 0.5956 | 1 |
| COAa | 'Cortical amygdalar area, anterior part' | 0.00976 | 0.02289 | 0.02058 | 0.00732 | 0.01161 | 0.01186 | 0.00826 | 0.02531 | 0.02412 | 0.01576 | 0.02636 | 0.02079 | 0.00931 | 0.03453 | 0.01768 | -0.0757 | 0.0148 | 0.2449 | 0.9173 | 0.0027 | 0.0907 | 0.7031 | 1 |
| COAp | 'Cortical amygdalar area, posterior part' | 0.00503 | 0.00796 | 0.00482 | 0.00246 | 0.00561 | 0.00434 | 0.00770 | 0.00961 | 0.00625 | 0.00848 | 0.00751 | 0.00340 | 0.00350 | 0.00694 | 0.00544 | 0.0104 | 0.1549 | 0.9878 | 0.0687 | 0.0669 | 0.8854 | 0.0298 | 1 |
| PAA | 'Piriform-amygdalar area' | 0.00468 | 0.01235 | 0.00570 | 0.00236 | 0.00505 | 0.00452 | 0.00678 | 0.01221 | 0.01153 | 0.01111 | 0.01453 | 0.00945 | 0.00027 | 0.00786 | 0.00520 | 0.0121 | 0.0018 | 0.8922 | 0.0798 | 0.0007 | 0.6603 | 0.0213 | 1 |
| TR | 'Postpiriform transition area' | 0.00106 | 0.00379 | 0.00174 | 0.00049 | 0.00222 | 0.00088 | 0.00064 | 0.00292 | 0.00079 | 0.00542 | 0.00600 | 0.00079 | 0.00028 | 0.00326 | 0.00262 | -0.0016 | 0.2033 | 0.7021 | 0.3827 | 0.0741 | 0.4350 | 0.1933 | 1 |
| HPF | 'Hippocampal formation' | 0.00256 | 0.00205 | 0.00135 | 0.00310 | 0.00389 | 0.00198 | 0.00128 | 0.00178 | 0.00076 | 0.00294 | 0.00273 | 0.00125 | 0.00057 | 0.00280 | 0.00078 | 0.0147 | 0.7906 | 0.3253 | 0.6089 | 0.5287 | 0.1646 | 0.3598 | 2 |
| CA1 | 'Field CA1' | 0.00026 | 0.00039 | 0.00012 | 0.00042 | 0.00033 | 0.00006 | 0.00021 | 0.00017 | 0.00007 | 0.00082 | 0.00050 | 0.00003 | 0.00012 | 0.00017 | 0.00016 | 0.0012 | 0.7647 | 0.3444 | 0.3788 | 0.4916 | 0.1609 | 0.1647 | 2 |
| CA2 | 'Field CA2' | 0.00012 | 0.00011 | 0.00007 | 0.00019 | 0.00015 | 0.00002 | 0.00011 | 0.00003 | 0.00002 | 0.00023 | 0.00016 | 0.00002 | 0.00013 | 0.00007 | 0.00006 | 0.0003 | 0.9922 | 0.5502 | 0.7683 | 0.9263 | 0.3421 | 0.5105 | 2 |
| CA3 | 'Field CA3' | 0.00048 | 0.00052 | 0.00038 | 0.00066 | 0.00053 | 0.00047 | 0.00025 | 0.00036 | 0.00015 | 0.00081 | 0.00078 | 0.00088 | 0.00026 | 0.00034 | 0.00003 | 0.0014 | 0.9638 | 0.8742 | 0.8204 | 0.8056 | 0.6463 | 0.5767 | 2 |
| DG | 'Dentate gyrus' | 0.00054 | 0.00063 | 0.00035 | 0.00068 | 0.00089 | 0.00037 | 0.00021 | 0.00054 | 0.00026 | 0.00070 | 0.00101 | 0.00062 | 0.00028 | 0.00028 | 0.00023 | 0.0024 | 0.9122 | 0.5854 | 0.3604 | 0.7123 | 0.3273 | 0.1795 | 2 |
| ENTl | 'Entorhinal area, lateral part' | 0.01786 | 0.01652 | 0.00298 | 0.01434 | 0.02296 | 0.01590 | 0.01826 | 0.01573 | 0.01002 | 0.01589 | 0.02442 | 0.00223 | 0.00049 | 0.00434 | 0.00485 | 0.1888 | 0.923 | 0.003 | 0.0315 | 0.7149 | 0.0004 | 0.0050 | 1 |
| ENTm | 'Entorhinal area, medial part, dorsal zone' | 0.00444 | 0.00550 | 0.00109 | 0.00245 | 0.00550 | 0.00387 | 0.00593 | 0.00631 | 0.00308 | 0.00587 | 0.00675 | 0.00052 | 0.00016 | 0.00186 | 0.00184 | 0.0448 | 0.6447 | 0.036 | 0.0149 | 0.3907 | 0.0105 | 0.0033 | 1 |
| PAR | 'Parasubiculum' | 0.00360 | 0.00230 | 0.00057 | 0.00434 | 0.00602 | 0.00276 | 0.00128 | 0.00174 | 0.00020 | 0.00260 | 0.00466 | 0.00048 | 0.00034 | 0.00036 | 0.00111 | 0.0338 | 0.6366 | 0.1127 | 0.2953 | 0.3829 | 0.0265 | 0.1174 | 1 |
| POST | 'Postsubiculum' | 0.00526 | 0.00162 | 0.00101 | 0.00742 | 0.00633 | 0.00504 | 0.00225 | 0.00138 | 0.00056 | 0.00201 | 0.00252 | 0.00130 | 0.00096 | 0.00148 | 0.00029 | 0.0433 | 0.0828 | 0.058 | 0.4896 | 0.0224 | 0.0099 | 0.2659 | 1 |
| PRE | 'Presubiculum' | 0.00332 | 0.00244 | 0.00085 | 0.00546 | 0.00298 | 0.00362 | 0.00122 | 0.00177 | 0.00047 | 0.00221 | 0.00530 | 0.00089 | 0.00086 | 0.00134 | 0.00032 | 0.0278 | 0.7965 | 0.1255 | 0.3924 | 0.5353 | 0.0337 | 0.1804 | 1 |
| SUB | 'Subiculum' | 0.00191 | 0.00097 | 0.00025 | 0.00383 | 0.00191 | 0.00170 | 0.00018 | 0.00034 | 0.00016 | 0.00073 | 0.00265 | 0.00024 | 0.00018 | 0.00044 | 0.00016 | 0.0170 | 0.6095 | 0.216 | 0.5095 | 0.3579 | 0.0701 | 0.2605 | 1 |
| CTXsp | 'Cortical subplate' | 0.00645 | 0.00733 | 0.00430 | 0.00529 | 0.00467 | 0.00631 | 0.00954 | 0.00627 | 0.00576 | 0.00944 | 0.00786 | 0.00518 | 0.00232 | 0.00624 | 0.00347 | 0.0332 | 0.8028 | 0.3397 | 0.1008 | 0.5438 | 0.1733 | 0.0458 | 2 |
| CLA | 'Claustrum' | 0.02057 | 0.01108 | 0.00573 | 0.01686 | 0.02129 | 0.02119 | 0.02294 | 0.00904 | 0.01206 | 0.00999 | 0.01322 | 0.00765 | 0.00328 | 0.00865 | 0.00333 | 0.1849 | 0.0034 | 0.0006 | 0.0534 | 0.0011 | 0.0002 | 0.0202 | 2 |
| EP | 'Endopiriform nucleus' | 0.01060 | 0.00856 | 0.00404 | 0.00481 | 0.00830 | 0.01422 | 0.01505 | 0.00892 | 0.00751 | 0.00953 | 0.00830 | 0.00462 | 0.00056 | 0.00656 | 0.00441 | 0.0919 | 0.7182 | 0.1418 | 0.0636 | 0.4444 | 0.0543 | 0.0144 | 1 |
| LA | 'Lateral amygdalar nucleus' | 0.00465 | 0.00539 | 0.00119 | 0.00299 | 0.00298 | 0.00463 | 0.00799 | 0.00229 | 0.00483 | 0.00883 | 0.00559 | 0.00151 | 0.00085 | 0.00168 | 0.00071 | 0.0460 | 0.9121 | 0.1152 | 0.1004 | 0.6943 | 0.0284 | 0.0220 | 2 |
| BLAa | 'Basolateral amygdalar nucleus, anterior part | 0.01635 | 0.01832 | 0.00516 | 0.01210 | 0.01462 | 0.01916 | 0.01953 | 0.01445 | 0.02162 | 0.01771 | 0.01948 | 0.00635 | 0.00491 | 0.00623 | 0.00316 | 0.1496 | 0.6986 | 0.0101 | 0.0023 | 0.4371 | 0.0012 | 0.0002 | 2 |
| BLAp | 'Basolateral amygdalar nucleus, posterior pa | 0.00540 | 0.00750 | 0.00335 | 0.00379 | 0.00745 | 0.00476 | 0.00561 | 0.00448 | 0.00739 | 0.00930 | 0.00883 | 0.00353 | 0.00182 | 0.00558 | 0.00246 | 0.0351 | 0.3332 | 0.2435 | 0.0553 | 0.1669 | 0.1192 | 0.0226 | 2 |
| BLAv | 'Basolateral amygdalar nucleus, ventral part' | 0.01065 | 0.01646 | 0.01265 | 0.00657 | 0.00923 | 0.01543 | 0.01139 | 0.01277 | 0.01343 | 0.01600 | 0.02362 | 0.01502 | 0.00292 | 0.02161 | 0.01106 | 0.0211 | 0.2346 | 0.8922 | 0.7073 | 0.1117 | 0.6612 | 0.4426 | 1 |
| BMAa | 'Basomedial amygdalar nucleus, anterior par | 0.00816 | 0.01107 | 0.01350 | 0.00589 | 0.00710 | 0.00999 | 0.00966 | 0.00938 | 0.01257 | 0.00991 | 0.01242 | 0.01271 | 0.01005 | 0.01892 | 0.01231 | -0.0416 | 0.1435 | 0.1259 | 0.5264 | 0.0658 | 0.0472 | 0.2858 | 1 |
| BMAp | 'Basomedial amygdalar nucleus, posterior pa | 0.00906 | 0.00909 | 0.00746 | 0.00613 | 0.00862 | 0.00913 | 0.01239 | 0.01112 | 0.00547 | 0.01071 | 0.00907 | 0.00457 | 0.00744 | 0.00837 | 0.00944 | 0.0203 | 0.9999 | 0.6197 | 0.6111 | 0.9877 | 0.3695 | 0.3612 | 1 |
| PA | 'Posterior amygdalar nucleus' | 0.00341 | 0.00423 | 0.00281 | 0.00257 | 0.00252 | 0.00366 | 0.00490 | 0.00456 | 0.00205 | 0.00488 | 0.00544 | 0.00058 | 0.00295 | 0.00285 | 0.00485 | 0.0077 | 0.6753 | 0.8348 | 0.477 | 0.4148 | 0.5814 | 0.2619 | 1 |
| STR | 'Striatum' | 0.00418 | 0.00384 | 0.00478 | 0.00291 | 0.00416 | 0.00410 | 0.00554 | 0.00333 | 0.00570 | 0.00321 | 0.00313 | 0.00560 | 0.00390 | 0.00619 | 0.00343 | -0.0007 | 0.9136 | 0.7691 | 0.585 | 0.6993 | 0.5046 | 0.3412 | 2 |
| CP | 'Caudoputamen' | 0.00751 | 0.00468 | 0.00356 | 0.00565 | 0.00384 | 0.00568 | 0.01487 | 0.00196 | 0.00376 | 0.00918 | 0.00383 | 0.00330 | 0.00504 | 0.00261 | 0.00330 | 0.0441 | 0.629 | 0.3799 | 0.7873 | 0.3728 | 0.1715 | 0.5223 | 3 |
| STRv | 'Striatum ventral region' | 0.00222 | 0.00370 | 0.00421 | 0.00216 | 0.00152 | 0.00197 | 0.00325 | 0.00318 | 0.00354 | 0.00462 | 0.00346 | 0.00505 | 0.00256 | 0.00628 | 0.00294 | -0.0171 | 0.0527 | 0.21 | 0.8563 | 0.0226 | 0.0830 | 0.6076 | 2 |
| NAc | 'Nucleus accumbens' | 0.03172 | 0.01904 | 0.01628 | 0.02104 | 0.02803 | 0.02880 | 0.04903 | 0.01726 | 0.03056 | 0.01164 | 0.01670 | 0.01852 | 0.00857 | 0.02228 | 0.01577 | 0.2146 | 0.2737 | 0.1578 | 0.8488 | 0.1312 | 0.0604 | 0.5993 | 3 |
| OT | 'Olfactory tubercle' | 0.01534 | 0.01624 | 0.01574 | 0.00947 | 0.01682 | 0.02063 | 0.01442 | 0.01926 | 0.01738 | 0.01047 | 0.01786 | 0.02075 | 0.00659 | 0.02357 | 0.01205 | 0.0330 | 0.9528 | 0.9957 | 0.9928 | 0.7767 | 0.9325 | 0.9123 | 1 |
| LSx | 'Lateral septal complex' | 0.00467 | 0.00483 | 0.00858 | 0.00587 | 0.00367 | 0.00165 | 0.00749 | 0.00373 | 0.00672 | 0.00422 | 0.00465 | 0.00972 | 0.00526 | 0.00719 | 0.01214 | -0.0459 | 0.9932 | 0.1982 | 0.1655 | 0.9145 | 0.0943 | 0.0620 | 3 |
| sAMY | 'Striatum-like amygdalar nuclei' | 0.00878 | 0.01287 | 0.01276 | 0.00685 | 0.01106 | 0.00700 | 0.01020 | 0.01122 | 0.01304 | 0.01276 | 0.01447 | 0.01850 | 0.00835 | 0.01552 | 0.00868 | -0.0243 | 0.0522 | 0.4012 | 0.999 | 0.0182 | 0.1975 | 0.9690 | 2 |
| CEA | 'Central amygdalar nucleus' | 0.00197 | 0.00215 | 0.00389 | 0.00130 | 0.00107 | 0.00092 | 0.00459 | 0.00129 | 0.00243 | 0.00350 | 0.00137 | 0.00358 | 0.00250 | 0.00733 | 0.00216 | -0.0158 | 0.9835 | 0.4472 | 0.4431 | 0.8637 | 0.2394 | 0.2270 | 2 |
| MEA | 'Medial amygdalar nucleus' | 0.00868 | 0.01360 | 0.01262 | 0.00633 | 0.01050 | 0.00981 | 0.00808 | 0.01671 | 0.01585 | 0.01011 | 0.01172 | 0.00979 | 0.01287 | 0.01485 | 0.01298 | -0.0381 | 0.0986 | 0.0699 | 0.8692 | 0.0374 | 0.0308 | 0.6283 | 1 |
| PAL | 'Pallidum' | 0.00643 | 0.00651 | 0.00757 | 0.00573 | 0.00617 | 0.00599 | 0.00785 | 0.00522 | 0.00925 | 0.00504 | 0.00654 | 0.00671 | 0.01072 | 0.00634 | 0.00650 | -0.0130 | 0.9972 | 0.6253 | 0.7526 | 0.9457 | 0.3649 | 0.4885 | 3 |
| GPe | 'Globus pallidus, external segment' | 0.00108 | 0.00103 | 0.00191 | 0.00116 | 0.00021 | 0.00027 | 0.00267 | 0.00083 | 0.00047 | 0.00255 | 0.00025 | 0.00136 | 0.00364 | 0.00094 | 0.00170 | -0.0142 | 0.9975 | 0.6005 | 0.5405 | 0.9490 | 0.3551 | 0.3092 | 3 |
| GPi | 'Globus pallidus, internal segment' | 0.00179 | 0.00205 | 0.00380 | 0.00232 | 0.00091 | 0.00059 | 0.00334 | 0.00130 | 0.00372 | 0.00238 | 0.00079 | 0.00351 | 0.00642 | 0.00119 | 0.00409 | -0.0302 | 0.9572 | 0.3259 | 0.4099 | 0.7857 | 0.1586 | 0.2122 | 3 |
| SI | 'Substantia innominata' | 0.00666 | 0.00669 | 0.00983 | 0.00458 | 0.00864 | 0.00638 | 0.00705 | 0.00519 | 0.01064 | 0.00459 | 0.00636 | 0.01099 | 0.00622 | 0.01407 | 0.00802 | -0.0206 | 0.9998 | 0.3209 | 0.3908 | 0.9851 | 0.1498 | 0.2040 | 2 |
| MA | 'Magnocellular nucleus' | 0.00934 | 0.00964 | 0.01713 | 0.00694 | 0.01143 | 0.00837 | 0.01061 | 0.00586 | 0.01698 | 0.00592 | 0.00979 | 0.01817 | 0.00789 | 0.02403 | 0.01845 | -0.0636 | 0.9937 | 0.1941 | 0.2645 | 0.9187 | 0.0684 | 0.1288 | 1 |
| MS | 'Medial septal nucleus' | 0.01031 | 0.01251 | 0.01892 | 0.01269 | 0.00925 | 0.00371 | 0.01559 | 0.01471 | 0.01134 | 0.01094 | 0.01307 | 0.02035 | 0.00879 | 0.02416 | 0.02238 | -0.0888 | 0.7149 | 0.1992 | 0.299 | 0.4451 | 0.0925 | 0.1232 | 3 |
| NDB | 'Diagonal band nucleus' | 0.02438 | 0.01392 | 0.01706 | 0.01910 | 0.02705 | 0.03107 | 0.02028 | 0.01546 | 0.02259 | 0.00757 | 0.01005 | 0.02147 | 0.02375 | 0.01704 | 0.00596 | 0.0895 | 0.119 | 0.3593 | 0.8217 | 0.0539 | 0.1830 | 0.5659 | 1 |
| TRS | 'Triangular nucleus of septum' | 0.00607 | 0.00625 | 0.01072 | 0.00810 | 0.00415 | 0.00150 | 0.01053 | 0.00307 | 0.00546 | 0.00827 | 0.00820 | 0.01588 | 0.00777 | 0.00570 | 0.01352 | -0.0564 | 0.9968 | 0.3617 | 0.3145 | 0.9422 | 0.1870 | 0.1479 | 3 |
| BST | 'Bed nuclei of the stria terminalis' | 0.00482 | 0.00590 | 0.00980 | 0.00419 | 0.00247 | 0.00267 | 0.00996 | 0.00456 | 0.01223 | 0.00434 | 0.00248 | 0.00518 | 0.01031 | 0.00937 | 0.01433 | -0.0609 | 0.9214 | 0.2096 | 0.4174 | 0.7110 | 0.1010 | 0.2227 | 3 |
| TH | 'Thalamus' | 0.00416 | 0.00595 | 0.00981 | 0.00589 | 0.00395 | 0.00315 | 0.00367 | 0.00661 | 0.00581 | 0.00725 | 0.00415 | 0.00667 | 0.01578 | 0.00532 | 0.01147 | -0.0840 | 0.1956 | 0.1862 | 0.3734 | 0.0934 | 0.0615 | 0.1708 | 3 |
| VENT | 'Ventral group of the dorsal thalamus' | 0.00135 | 0.00111 | 0.00403 | 0.00271 | 0.00031 | 0.00079 | 0.00159 | 0.00126 | 0.00111 | 0.00158 | 0.00047 | 0.00037 | 0.01055 | 0.00037 | 0.00484 | -0.0472 | 0.9067 | 0.5792 | 0.5254 | 0.6873 | 0.3192 | 0.2731 | 3 |
| SPF | 'Subparafascicular nucleus' | 0.00283 | 0.00297 | 0.00909 | 0.00384 | 0.00222 | 0.00241 | 0.00285 | 0.00522 | 0.00247 | 0.00373 | 0.00045 | 0.00024 | 0.02081 | 0.00267 | 0.01265 | -0.1037 | 0.991 | 0.4772 | 0.4953 | 0.9014 | 0.2361 | 0.2536 | 3 |
| SPA | 'Subparafascicular area' | 0.00574 | 0.00990 | 0.01948 | 0.00167 | 0.00213 | 0.01133 | 0.00782 | 0.02195 | 0.00855 | 0.00663 | 0.00248 | 0.00296 | 0.02732 | 0.01384 | 0.03379 | -0.1912 | 0.6841 | 0.2643 | 0.5108 | 0.4201 | 0.1081 | 0.2808 | 3 |
| PP | 'Peripeduncular nucleus' | 0.00696 | 0.00961 | 0.01347 | 0.01080 | 0.00767 | 0.00420 | 0.00517 | 0.00421 | 0.00385 | 0.01514 | 0.01523 | 0.01155 | 0.01349 | 0.01612 | 0.01273 | -0.0747 | 0.7505 | 0.0299 | 0.5444 | 0.4837 | 0.0103 | 0.2944 | 3 |
| GENd | 'Geniculate group, dorsal thalamus' | 0.00754 | 0.00406 | 0.00352 | 0.01348 | 0.00544 | 0.00432 | 0.00694 | 0.00024 | 0.00006 | 0.01096 | 0.00499 | 0.00325 | 0.00809 | 0.00136 | 0.00139 | 0.0249 | 0.5715 | 0.3381 | 0.9826 | 0.3301 | 0.1712 | 0.8626 | 3 |
| LAT | 'Lateral group of the dorsal thalamus' | 0.00148 | 0.00214 | 0.00282 | 0.00294 | 0.00131 | 0.00077 | 0.00090 | 0.00131 | 0.00126 | 0.00401 | 0.00200 | 0.00339 | 0.00438 | 0.00166 | 0.00184 | -0.0209 | 0.7088 | 0.3084 | 0.753 | 0.4468 | 0.1539 | 0.4894 | 3 |
| ATN | 'Anterior group of the dorsal thalamus' | 0.00131 | 0.00192 | 0.00508 | 0.00208 | 0.00091 | 0.00071 | 0.00152 | 0.00131 | 0.00267 | 0.00242 | 0.00127 | 0.00261 | 0.00684 | 0.00127 | 0.00960 | -0.0550 | 0.4581 | 0.2665 | 0.3576 | 0.2501 | 0.1004 | 0.1567 | 3 |
| MED | 'Medial group of the dorsal thalamus' | 0.00260 | 0.00765 | 0.01230 | 0.00358 | 0.00144 | 0.00160 | 0.00379 | 0.00821 | 0.00959 | 0.00776 | 0.00504 | 0.00957 | 0.01269 | 0.00593 | 0.02101 | -0.1269 | 0.0145 | 0.1079 | 0.4372 | 0.0045 | 0.0253 | 0.2151 | 3 |
| MTN | 'Midline group of the dorsal thalamus' | 0.00703 | 0.01827 | 0.02518 | 0.00925 | 0.00693 | 0.00426 | 0.00769 | 0.01861 | 0.01940 | 0.01736 | 0.01769 | 0.03057 | 0.02262 | 0.02115 | 0.02639 | -0.2064 | 0.0011 | 0.0023 | 0.0869 | 0.0001 | 0.0002 | 0.0185 | 3 |
| ILM | 'Intralaminar nuclei of the dorsal thalamus' | 0.00130 | 0.00339 | 0.00583 | 0.00155 | 0.00067 | 0.00047 | 0.00251 | 0.00403 | 0.00383 | 0.00411 | 0.00160 | 0.00327 | 0.00794 | 0.00261 | 0.00952 | -0.0629 | 0.078 | 0.1435 | 0.449 | 0.0332 | 0.0427 | 0.2259 | 3 |
| RT | 'Reticular nucleus of the thalamus' | 0.00068 | 0.00088 | 0.00293 | 0.00110 | 0.00043 | 0.00033 | 0.00084 | 0.00091 | 0.00063 | 0.00178 | 0.00019 | 0.00085 | 0.00703 | 0.00127 | 0.00259 | -0.0347 | 0.8598 | 0.373 | 0.4296 | 0.6127 | 0.1643 | 0.2070 | 3 |
| IGL | 'Intergeniculate leaflet of the lateral genicula | 0.00235 | 0.00204 | 0.00345 | 0.00634 | 0.00166 | 0.00085 | 0.00055 | 0.00070 | 0.00047 | 0.00539 | 0.00159 | 0.00076 | 0.00713 | 0.00043 | 0.00549 | -0.0326 | 0.983 | 0.8692 | 0.7762 | 0.8656 | 0.6276 | 0.5127 | 3 |
| LGv | 'Ventral part of the lateral geniculate comple | 0.00121 | 0.00217 | 0.00472 | 0.00304 | 0.00098 | 0.00036 | 0.00046 | 0.00074 | 0.00030 | 0.00416 | 0.00349 | 0.00714 | 0.00468 | 0.00505 | 0.00199 | -0.0400 | 0.6992 | 0.0795 | 0.2565 | 0.4349 | 0.0290 | 0.1262 | 3 |
| SubG | 'Subgeniculate nucleus' | 0.00209 | 0.00259 | 0.00307 | 0.00563 | 0.00070 | 0.00034 | 0.00168 | 0.00072 | 0.00054 | 0.00798 | 0.00112 | 0.00425 | 0.00328 | 0.00197 | 0.00277 | -0.0231 | 0.9711 | 0.7496 | 0.9648 | 0.8244 | 0.4802 | 0.8061 | 3 |
| MHb | 'Medial habenula' | 0.00545 | 0.01275 | 0.01638 | 0.00659 | 0.00731 | 0.00412 | 0.00380 | 0.00799 | 0.00800 | 0.01713 | 0.01785 | 0.02449 | 0.01618 | 0.00520 | 0.01963 | -0.1293 | 0.1428 | 0.1435 | 0.7536 | 0.0447 | 0.0403 | 0.4893 | 3 |
| LHb | 'Lateral habenula' | 0.00186 | 0.00796 | 0.01700 | 0.00202 | 0.00209 | 0.00159 | 0.00172 | 0.00763 | 0.01032 | 0.01049 | 0.00340 | 0.02252 | 0.01277 | 0.00906 | 0.02367 | -0.1838 | 0.0681 | 0.0491 | 0.165 | 0.0103 | 0.0057 | 0.0627 | 3 |
| HY | 'Hypothalamus' | 0.00675 | 0.00902 | 0.01004 | 0.00727 | 0.00898 | 0.00605 | 0.00468 | 0.01100 | 0.01032 | 0.00692 | 0.00787 | 0.01024 | 0.01138 | 0.01106 | 0.00747 | -0.0376 | 0.2767 | 0.0919 | 0.7355 | 0.1384 | 0.0415 | 0.4707 | 2 |
| PVZ | 'Periventricular zone' | 0.00456 | 0.00829 | 0.00640 | 0.00492 | 0.00560 | 0.00484 | 0.00287 | 0.01253 | 0.00471 | 0.00599 | 0.00992 | 0.00637 | 0.00484 | 0.00819 | 0.00620 | -0.0162 | 0.2437 | 0.1861 | 0.6264 | 0.0958 | 0.0878 | 0.3639 | 1 |
| PVR | 'Periventricular region' | 0.00808 | 0.01182 | 0.01257 | 0.00811 | 0.00979 | 0.00933 | 0.00511 | 0.01623 | 0.01362 | 0.00877 | 0.00866 | 0.01219 | 0.01389 | 0.01507 | 0.00911 | -0.0504 | 0.2842 | 0.0834 | 0.9432 | 0.1322 | 0.0363 | 0.7548 | 1 |

| | | | | | | | | | | | | | | | | | | | | | | | | |
|---|---|---|---|---|---|---|---|---|---|---|---|---|---|---|---|---|---|---|---|---|---|---|---|---|
| AHN | 'Anterior hypothalamic nucleus' | 0.01526 | 0.01678 | 0.01698 | 0.02051 | 0.01631 | 0.01564 | 0.00859 | 0.02385 | 0.01946 | 0.01383 | 0.00996 | 0.02004 | 0.02256 | 0.01167 | 0.01362 | -0.0472 | 0.9228 | 0.8838 | 0.9987 | 0.7134 | 0.6484 | 0.9618 | 1 |
| MBO | 'Mammillary body' | 0.01285 | 0.01370 | 0.01591 | 0.01407 | 0.01684 | 0.01336 | 0.00712 | 0.02143 | 0.01526 | 0.01090 | 0.00722 | 0.01416 | 0.02398 | 0.01368 | 0.01182 | -0.0573 | 0.9708 | 0.664 | 0.8561 | 0.8240 | 0.4049 | 0.6097 | 1 |
| MPN | 'Medial preoptic nucleus' | 0.00810 | 0.01465 | 0.01282 | 0.00795 | 0.00949 | 0.00981 | 0.00515 | 0.02361 | 0.01476 | 0.01080 | 0.00941 | 0.00784 | 0.01875 | 0.01006 | 0.01464 | -0.0707 | 0.2505 | 0.2839 | 0.8947 | 0.0997 | 0.1251 | 0.6656 | 1 |
| PMd | 'Dorsal premammillary nucleus' | 0.01371 | 0.01921 | 0.01581 | 0.01097 | 0.01499 | 0.01536 | 0.01351 | 0.03642 | 0.02678 | 0.01017 | 0.00346 | 0.01256 | 0.02410 | 0.00599 | 0.02061 | -0.0582 | 0.7682 | 0.8747 | 0.9188 | 0.4972 | 0.6332 | 0.7060 | 1 |
| PMv | 'Ventral premammillary nucleus' | 0.01126 | 0.01673 | 0.01799 | 0.01082 | 0.02042 | 0.01211 | 0.00168 | 0.02832 | 0.01008 | 0.01269 | 0.01585 | 0.01970 | 0.00534 | 0.03258 | 0.01436 | -0.0532 | 0.6128 | 0.6172 | 0.9823 | 0.3637 | 0.3645 | 0.8627 | 1 |
| PVHd | 'Paraventricular hypothalamic nucleus, desce | 0.00418 | 0.01020 | 0.01369 | 0.00478 | 0.00479 | 0.00238 | 0.00479 | 0.01523 | 0.01259 | 0.00907 | 0.00392 | 0.01227 | 0.01963 | 0.00692 | 0.01595 | -0.1307 | 0.1708 | 0.0732 | 0.6285 | 0.0539 | 0.0141 | 0.3762 | 2 |
| VMH | 'Ventromedial hypothalamic nucleus' | 0.00460 | 0.00998 | 0.00574 | 0.00613 | 0.00752 | 0.00345 | 0.00130 | 0.01399 | 0.00717 | 0.00736 | 0.01142 | 0.00475 | 0.00417 | 0.00969 | 0.00437 | -0.0067 | 0.1052 | 0.8264 | 0.1961 | 0.0471 | 0.5731 | 0.0922 | 1 |
| PH | 'Posterior hypothalamic nucleus' | 0.00500 | 0.01043 | 0.01144 | 0.00503 | 0.00454 | 0.00382 | 0.00663 | 0.01648 | 0.01287 | 0.00754 | 0.00483 | 0.00868 | 0.01832 | 0.00312 | 0.01563 | -0.0969 | 0.2428 | 0.2911 | 0.9706 | 0.0897 | 0.1147 | 0.8233 | 2 |
| LH | 'Hypothalamic lateral zone' | 0.00797 | 0.01147 | 0.01460 | 0.00632 | 0.01044 | 0.00854 | 0.00658 | 0.01216 | 0.01728 | 0.00866 | 0.00779 | 0.01711 | 0.01645 | 0.01558 | 0.00926 | -0.0693 | 0.3853 | 0.0567 | 0.5426 | 0.1886 | 0.0177 | 0.3079 | 2 |
| ZI | 'Zona incerta' | 0.00442 | 0.00514 | 0.01197 | 0.00849 | 0.00294 | 0.00291 | 0.00334 | 0.00327 | 0.00760 | 0.00666 | 0.00305 | 0.00480 | 0.02182 | 0.00818 | 0.01309 | -0.1141 | 0.9148 | 0.2533 | 0.3021 | 0.7002 | 0.1038 | 0.1286 | 3 |
| ME | 'Median eminence' | 0.00089 | 0.00240 | 0.00298 | 0.00310 | 0.00034 | 0.00012 | 0.00001 | 0.00325 | 0.00073 | 0.00397 | 0.00166 | 0.00056 | 0.00008 | 0.00853 | 0.00273 | -0.0226 | 0.3781 | 0.6146 | 0.9593 | 0.1979 | 0.3539 | 0.7915 | 1 |
| MB-un | 'Midbrain' | 0.00797 | 0.00832 | 0.01271 | 0.00985 | 0.00902 | 0.00627 | 0.00673 | 0.00708 | 0.00735 | 0.00990 | 0.00895 | 0.01248 | 0.01516 | 0.01078 | 0.01242 | -0.0632 | 0.9452 | 0.0216 | 0.0218 | 0.7605 | 0.0092 | 0.0080 | 3 |
| MB-sen | 'Midbrain, sensory related' | 0.01225 | 0.00951 | 0.01401 | 0.01252 | 0.01330 | 0.01256 | 0.01063 | 0.00446 | 0.00466 | 0.01453 | 0.01441 | 0.01822 | 0.01526 | 0.01391 | 0.00863 | -0.0156 | 0.6546 | 0.7039 | 0.4576 | 0.3842 | 0.4322 | 0.2458 | 3 |
| SCs | 'Superior colliculus, sensory related' | 0.00293 | 0.00343 | 0.00481 | 0.00443 | 0.00249 | 0.00427 | 0.00055 | 0.00243 | 0.00106 | 0.00553 | 0.00471 | 0.00659 | 0.00675 | 0.00348 | 0.00243 | -0.0255 | 0.9309 | 0.4372 | 0.649 | 0.7284 | 0.2357 | 0.3948 | 1 |
| IC | 'Inferior colliculus' | 0.00744 | 0.00504 | 0.00784 | 0.01098 | 0.01000 | 0.00300 | 0.00577 | 0.00363 | 0.00390 | 0.00671 | 0.00592 | 0.01000 | 0.00710 | 0.00655 | 0.00772 | -0.0139 | 0.5171 | 0.978 | 0.0878 | 0.2781 | 0.8468 | 0.0399 | 3 |
| MB-mot | 'Midbrain, motor related' | 0.00277 | 0.00322 | 0.00518 | 0.00520 | 0.00209 | 0.00197 | 0.00179 | 0.00197 | 0.00203 | 0.00647 | 0.00242 | 0.00546 | 0.00730 | 0.00387 | 0.00408 | -0.0378 | 0.9394 | 0.1643 | 0.3797 | 0.7487 | 0.0776 | 0.1958 | 3 |
| SNr | 'Substantia nigra, reticular part' | 0.00330 | 0.00406 | 0.00703 | 0.00419 | 0.00206 | 0.00292 | 0.00403 | 0.00446 | 0.00532 | 0.00452 | 0.00193 | 0.00959 | 0.00854 | 0.00589 | 0.00409 | -0.0465 | 0.6898 | 0.1059 | 0.1988 | 0.4285 | 0.0325 | 0.0865 | 2 |
| VTA | 'Ventral tegmental area' | 0.01588 | 0.02282 | 0.02446 | 0.02214 | 0.01372 | 0.01406 | 0.01359 | 0.02389 | 0.02954 | 0.02026 | 0.01757 | 0.02620 | 0.02905 | 0.02007 | 0.02252 | -0.1166 | 0.1765 | 0.0559 | 0.8721 | 0.0823 | 0.0247 | 0.6323 | 3 |
| SCm | 'Superior colliculus, motor related' | 0.00683 | 0.00580 | 0.01010 | 0.01162 | 0.00759 | 0.00419 | 0.00390 | 0.00319 | 0.00311 | 0.01020 | 0.00670 | 0.00900 | 0.01185 | 0.00816 | 0.01139 | -0.0551 | 0.9109 | 0.3287 | 0.1631 | 0.6928 | 0.1553 | 0.0658 | 3 |
| PAG | 'Periaqueductal gray' | 0.00646 | 0.00484 | 0.00763 | 0.01116 | 0.00518 | 0.00674 | 0.00276 | 0.00199 | 0.00032 | 0.01464 | 0.00240 | 0.00183 | 0.01068 | 0.00188 | 0.01613 | -0.0501 | 0.9036 | 0.9529 | 0.8358 | 0.6795 | 0.7763 | 0.5833 | 3 |
| PRC | 'Precommissural nucleus' | 0.00987 | 0.01800 | 0.02155 | 0.01151 | 0.00489 | 0.01678 | 0.00632 | 0.01295 | 0.02932 | 0.01685 | 0.01289 | 0.02043 | 0.02699 | 0.02293 | 0.01584 | -0.1312 | 0.2823 | 0.0403 | 0.7293 | 0.1369 | 0.0171 | 0.4634 | 3 |
| APN | 'Anterior pretectal nucleus' | 0.00044 | 0.00071 | 0.00223 | 0.00129 | 0.00016 | 0.00014 | 0.00018 | 0.00020 | 0.00018 | 0.00183 | 0.00063 | 0.00024 | 0.00536 | 0.00021 | 0.00312 | -0.0295 | 0.8469 | 0.436 | 0.5352 | 0.5912 | 0.2104 | 0.2886 | 3 |
| MPT | 'Medial pretectal area' | 0.00971 | 0.01661 | 0.01500 | 0.02077 | 0.01386 | 0.00153 | 0.00269 | 0.00807 | 0.00866 | 0.02371 | 0.02601 | 0.02722 | 0.02059 | 0.00372 | 0.00846 | -0.0780 | 0.5826 | 0.7486 | 0.9729 | 0.3394 | 0.4852 | 0.8305 | 3 |
| SNc | 'Substantia nigra, compact part' | 0.00215 | 0.00263 | 0.00455 | 0.00224 | 0.00281 | 0.00108 | 0.00246 | 0.00248 | 0.00265 | 0.00446 | 0.00092 | 0.00531 | 0.00743 | 0.00421 | 0.00125 | -0.0317 | 0.8326 | 0.2954 | 0.4562 | 0.5777 | 0.1230 | 0.2403 | 2 |
| PPN | 'Pedunculopontine nucleus' | 0.00224 | 0.00251 | 0.00705 | 0.00464 | 0.00177 | 0.00141 | 0.00113 | 0.00142 | 0.00093 | 0.00654 | 0.00115 | 0.01054 | 0.01128 | 0.00376 | 0.00263 | -0.0679 | 0.9836 | 0.2293 | 0.2827 | 0.8687 | 0.0902 | 0.1333 | 3 |
| IF | 'Intertascicular nucleus raphe' | 0.00119 | 0.00221 | 0.00231 | 0.00144 | 0.00020 | 0.00131 | 0.00180 | 0.00420 | 0.00170 | 0.00253 | 0.00041 | 0.00198 | 0.00300 | 0.00111 | 0.00313 | -0.0186 | 0.5213 | 0.2239 | 0.9942 | 0.2824 | 0.1056 | 0.9215 | 2 |
| IPN | 'Interpeduncular nucleus' | 0.00201 | 0.00155 | 0.00160 | 0.00344 | 0.00069 | 0.00250 | 0.00143 | 0.00177 | 0.00167 | 0.00202 | 0.00073 | 0.00071 | 0.00231 | 0.00029 | 0.00311 | -0.0030 | 0.7746 | 0.8932 | 0.9965 | 0.5117 | 0.6639 | 0.9421 | 2 |
| RL | 'Rostral linear nucleus raphe' | 0.00317 | 0.00707 | 0.01023 | 0.00433 | 0.00046 | 0.00411 | 0.00376 | 0.01370 | 0.00146 | 0.01056 | 0.00256 | 0.00346 | 0.01407 | 0.00130 | 0.02209 | -0.1171 | 0.4983 | 0.4249 | 0.8486 | 0.2596 | 0.2014 | 0.5991 | 3 |
| CLI | 'Central linear nucleus raphe' | 0.00369 | 0.00449 | 0.01023 | 0.00380 | 0.00002 | 0.00672 | 0.00421 | 0.00749 | 0.00047 | 0.00980 | 0.00019 | 0.00012 | 0.01665 | 0.00087 | 0.02327 | -0.1148 | 0.9567 | 0.5711 | 0.6607 | 0.7845 | 0.3134 | 0.3960 | 3 |
| DR | 'Dorsal nucleus raphe' | 0.00474 | 0.00492 | 0.00566 | 0.00466 | 0.00079 | 0.01234 | 0.00118 | 0.00200 | 0.00000 | 0.01756 | 0.00012 | 0.00057 | 0.00854 | 0.00026 | 0.01326 | -0.0385 | 0.9993 | 0.9737 | 0.9894 | 0.9729 | 0.8332 | 0.8940 | 3 |
| P | 'Pons' | 0.00734 | 0.00620 | 0.00703 | 0.00926 | 0.00952 | 0.00497 | 0.00563 | 0.00578 | 0.00595 | 0.00590 | 0.00715 | 0.00873 | 0.00873 | 0.00597 | 0.00470 | -0.0009 | 0.6552 | 0.9783 | 0.7315 | 0.3870 | 0.8485 | 0.4612 | 3 |
| P-sen | 'Pons, sensory related' | 0.00890 | 0.00705 | 0.00744 | 0.01356 | 0.00800 | 0.00303 | 0.01099 | 0.00570 | 0.00875 | 0.00631 | 0.00745 | 0.01043 | 0.00948 | 0.00548 | 0.00438 | 0.0079 | 0.7354 | 0.8569 | 0.969 | 0.4641 | 0.6099 | 0.8187 | 3 |
| NLL | 'Nucleus of the lateral lemniscus' | 0.00120 | 0.00205 | 0.00482 | 0.00154 | 0.00178 | 0.00104 | 0.00043 | 0.00220 | 0.00190 | 0.00173 | 0.00236 | 0.00674 | 0.00472 | 0.00561 | 0.00219 | -0.0385 | 0.1202 | 0.0585 | 0.1237 | 0.0424 | 0.0117 | 0.0299 | 1 |
| P-mot | 'Pons, motor related' | 0.00715 | 0.00560 | 0.00601 | 0.01348 | 0.00549 | 0.00578 | 0.00385 | 0.00526 | 0.00377 | 0.00721 | 0.00616 | 0.00775 | 0.00945 | 0.00317 | 0.00365 | -0.0053 | 0.7867 | 0.9038 | 0.9696 | 0.5205 | 0.6809 | 0.8190 | 3 |
| P-sat | 'Pons, behavioral state related' | 0.00428 | 0.00472 | 0.00696 | 0.00940 | 0.00289 | 0.00294 | 0.00189 | 0.00449 | 0.00312 | 0.00840 | 0.00289 | 0.00424 | 0.01323 | 0.00216 | 0.00820 | -0.0588 | 0.9766 | 0.6637 | 0.7144 | 0.8429 | 0.4045 | 0.4482 | 3 |
| MY | 'Medulla' | 0.00609 | 0.00642 | 0.00532 | 0.00421 | 0.00824 | 0.00342 | 0.00849 | 0.00608 | 0.00614 | 0.00492 | 0.00853 | 0.00724 | 0.00648 | 0.00408 | 0.00350 | 0.0159 | 0.9751 | 0.8849 | 0.647 | 0.8367 | 0.6502 | 0.3903 | 3 |
| MY-sen | 'Medulla, sensory related' | 0.00628 | 0.00647 | 0.00541 | 0.01334 | 0.00524 | 0.00190 | 0.00463 | 0.00744 | 0.00615 | 0.00726 | 0.00503 | 0.00779 | 0.00843 | 0.00366 | 0.00177 | -0.0081 | 0.9968 | 0.9539 | 0.818 | 0.9420 | 0.7791 | 0.5588 | 3 |
| MY-mot | 'Medulla, motor related' | 0.00330 | 0.00248 | 0.00307 | 0.00986 | 0.00159 | 0.00029 | 0.00144 | 0.00293 | 0.00137 | 0.00321 | 0.00240 | 0.00469 | 0.00449 | 0.00230 | 0.00080 | -0.0122 | 0.931 | 0.9952 | 0.8356 | 0.7277 | 0.9284 | 0.5797 | 3 |
| MY-sat | 'Medulla, behavioral state related' | 0.00477 | 0.00930 | 0.00784 | 0.00644 | 0.00638 | 0.00251 | 0.00376 | 0.01313 | 0.00412 | 0.00798 | 0.01197 | 0.01671 | 0.00644 | 0.00818 | 0.00003 | -0.0251 | 0.222 | 0.6946 | 0.9303 | 0.0931 | 0.4230 | 0.7280 | 3 |
