## Supplementary material for "Brain-wide projections and differential encoding of prefrontal neuronal classes underlying learned and innate threat avoidance": Table 3

**Table 3.** ANOVA Tables for Regional Axon Density Comparisons. Related to Figures 1–3.

**Figure 1E (Distinction Score)**

| **Repeated measures ANOVA summary** | | | |
|---|---|---|---|
| F | 14.46 | | |
| P value | <0.0001 | | |
| **Multiple comparisons (Benjamini, Krieger and Yekutieli)** | Summary | q value | Individual P value |
| DeepTraCE vs untrained TRAILMAP mod | **** | <0.0001 | <0.0001 |
| DeepTraCE vs Best Single Mod | ** | 0.0057 | 0.0039 |
| DeepTraCE vs Max Probability of Model | *** | 0.0004 | 0.0002 |
| DeepTraCE vs Sum of Model | ** | 0.0069 | 0.0063 |

**Whole-Brain 2-Way ANOVA (Figures 2, 3, S6)**

| **ANOVA table** | F (DFn, DFd | P value |
|---|---|---|
| Cell Type x Brain Region | F (278, 1251) = 3.959 | P<0.0001 |
| Cell Type | F (2, 9) = 5.690 | P=0.0253 |
| Brain Region | F (139.0, 1251) = 20.28 | P<0.0001 |

**Figure 3C: ORB Layer-Specific Innervation**

| **ANOVA table** | F (DFn, DFd | P value |
|---|---|---|
| Layer x Cell Type | F (6, 27) = 4.225 | P=0.0040 |
| Layer | F (1.539, 13.85) = 21.45 | P=0.0001 |
| Cell Type | F (2, 9) = 39.86 | P<0.0001 |

**Figure 3C: AUDs Layer-Specific Innervation**

| **ANOVA table** | F (DFn, DFd | P value |
|---|---|---|
| Layer x Cell Type | F (8, 36) = 1.635 | P=0.1493 |
| Layer | F (1.215, 10.93) = 7.283 | P=0.0172 |
| Cell Type | F (2, 9) = 44.48 | P<0.0001 |

**Figure 3C: TEa Layer-Specific Innervation**

| **ANOVA table** | F (DFn, DFd | P value |
|---|---|---|
| Layer x Cell Type | F (8, 36) = 2.466 | P=0.0305 |
| Layer | F (1.324, 11.92) = 4.447 | P=0.0484 |
| Cell Type | F (2, 9) = 45.49 | P<0.0001 |

**Figure 3C: ECT/PERI Layer-Specific Innervation**

| **ANOVA table** | F (DFn, DFd | P value |
|---|---|---|
| Layer x Cell Type | F (6, 27) = 3.490 | P=0.0110 |
| Layer | F (1.118, 10.06) = 2.843 | P=0.1205 |
| Cell Type | F (2, 9) = 27.44 | P=0.0001 |
