## Supplementary material for "Brain-wide projections and differential encoding of prefrontal neuronal classes underlying learned and innate threat avoidance": Table 4

**Table 4.** Quantification of TRAPed cell density by brain region. Related to Figure 4.

| Abbrev. | Allen Brain Atlas ID | PMA1 | PMA2 | PMA3 | PMA4 | Control1 | Control2 | Control3 | Control4 | Control4 | PC1 Weight |
|---|---|---|---|---|---|---|---|---|---|---|---|
| FRP | 'Frontal pole, cerebral cortex' | 0.00317 | 0.00341 | 0.00707 | 0.00441 | 0.00182 | 0.00254 | 0.00861 | 0.01356 | 0.00211 | -0.023057434 |
| MOp | 'Primary motor area' | 0.00579 | 0.00571 | 0.00404 | 0.00942 | 0.00444 | 0.00148 | 0.00253 | 0.00397 | 0.00385 | 0.11882221 |
| MOs | 'Secondary motor area' | 0.00501 | 0.00445 | 0.00317 | 0.00643 | 0.00334 | 0.00145 | 0.00212 | 0.00368 | 0.00307 | 0.077040964 |
| SSp | 'Primary somatosensory area' | 0.00750 | 0.00648 | 0.00579 | 0.01118 | 0.00569 | 0.00213 | 0.00446 | 0.00576 | 0.00548 | 0.123692076 |
| SSs | 'Supplemental somatosensory area' | 0.00678 | 0.00625 | 0.00566 | 0.00891 | 0.00515 | 0.00240 | 0.00444 | 0.00484 | 0.00454 | 0.09431028 |
| GU | 'Gustatory areas' | 0.00417 | 0.00473 | 0.00425 | 0.00525 | 0.00607 | 0.00281 | 0.00242 | 0.00321 | 0.00438 | 0.034970066 |
| VISC | 'Visceral area' | 0.00448 | 0.00404 | 0.00392 | 0.00523 | 0.00397 | 0.00322 | 0.00493 | 0.00315 | 0.00364 | 0.022171316 |
| AUDd | 'Dorsal auditory area' | 0.00712 | 0.00769 | 0.00783 | 0.01131 | 0.00500 | 0.00266 | 0.00591 | 0.00429 | 0.00577 | 0.129047686 |
| AUDp | 'Primary auditory area' | 0.00988 | 0.00953 | 0.00908 | 0.01336 | 0.00534 | 0.00552 | 0.00582 | 0.00603 | 0.00687 | 0.144700808 |
| AUDpo | 'Posterior auditory area' | 0.00886 | 0.00758 | 0.00819 | 0.01156 | 0.00492 | 0.00412 | 0.00742 | 0.00558 | 0.00590 | 0.111094333 |
| AUDv | 'Ventral auditory area' | 0.01109 | 0.01138 | 0.01009 | 0.01375 | 0.00647 | 0.00500 | 0.00708 | 0.00727 | 0.00711 | 0.155184172 |
| VISal | 'Anterolateral visual area' | 0.00930 | 0.00984 | 0.00803 | 0.01346 | 0.00479 | 0.00503 | 0.00870 | 0.00654 | 0.00822 | 0.12931079 |
| VISam | 'Anteromedial visual area' | 0.00971 | 0.01198 | 0.00858 | 0.01379 | 0.00619 | 0.00459 | 0.00691 | 0.00773 | 0.00922 | 0.153587341 |
| VISl | 'Lateral visual area' | 0.01161 | 0.00964 | 0.00928 | 0.01504 | 0.00889 | 0.00659 | 0.00988 | 0.00993 | 0.00859 | 0.107482394 |
| VISp | 'Primary visual area' | 0.01433 | 0.01262 | 0.01260 | 0.01688 | 0.00983 | 0.00801 | 0.01003 | 0.01162 | 0.01049 | 0.139084682 |
| VISpl | 'Posterolateral visual area' | 0.01070 | 0.01049 | 0.00706 | 0.00763 | 0.00919 | 0.00865 | 0.01180 | 0.01075 | 0.00804 | -0.025695417 |
| VISpm | 'posteromedial visual area' | 0.01098 | 0.01297 | 0.00928 | 0.01324 | 0.00880 | 0.00685 | 0.00840 | 0.00916 | 0.00981 | 0.110772464 |
| ACA | 'Anterior cingulate area' | 0.00968 | 0.01204 | 0.00740 | 0.01168 | 0.00897 | 0.00436 | 0.00720 | 0.00914 | 0.00808 | 0.109385742 |
| PL | 'Prelimbic area' | 0.00736 | 0.00764 | 0.00698 | 0.00949 | 0.00557 | 0.00302 | 0.00505 | 0.00642 | 0.00568 | 0.097274786 |
| ILA | 'Infralimbic area' | 0.00754 | 0.00986 | 0.00787 | 0.01026 | 0.00912 | 0.00600 | 0.00538 | 0.00591 | 0.00813 | 0.075655386 |
| ORBl | 'Orbital area, lateral part' | 0.00716 | 0.00687 | 0.00627 | 0.00930 | 0.00391 | 0.00327 | 0.00568 | 0.00690 | 0.00539 | 0.088445623 |
| ORBm | 'Orbital area, medial part' | 0.00664 | 0.00733 | 0.00672 | 0.00734 | 0.00637 | 0.00713 | 0.00683 | 0.00428 | 0.00602 | 0.017216403 |
| ORBvl | 'Orbital area, ventrolateral part' | 0.00846 | 0.01005 | 0.00878 | 0.01172 | 0.00618 | 0.00484 | 0.00869 | 0.01056 | 0.00694 | 0.093498991 |
| AId | 'Agranular insular area, dorsal part' | 0.00271 | 0.00300 | 0.00190 | 0.00382 | 0.00324 | 0.00135 | 0.00291 | 0.00270 | 0.00313 | 0.022942893 |
| AIp | 'Agranular insular area, posterior part' | 0.00290 | 0.00247 | 0.00295 | 0.00238 | 0.00415 | 0.00328 | 0.00372 | 0.00327 | 0.00308 | -0.025218962 |
| AIv | 'Agranular insular area, ventral part' | 0.00432 | 0.00570 | 0.00491 | 0.00595 | 0.00467 | 0.00418 | 0.00461 | 0.00345 | 0.00490 | 0.031055217 |
| RSP | 'Retrosplenial area' | 0.01019 | 0.01229 | 0.00935 | 0.00842 | 0.00701 | 0.00473 | 0.00693 | 0.00767 | 0.00868 | 0.08313066 |
| VISrl | 'Rostrolateral visual area' | 0.01460 | 0.00910 | 0.01057 | 0.01618 | 0.00671 | 0.00665 | 0.00971 | 0.00986 | 0.00844 | 0.141023559 |
| TEa | 'Temporal association areas' | 0.00870 | 0.00945 | 0.00755 | 0.01154 | 0.00544 | 0.00506 | 0.00722 | 0.00642 | 0.00664 | 0.107962164 |
| PERI | 'Perirhinal area' | 0.00320 | 0.00366 | 0.00326 | 0.00222 | 0.00545 | 0.00815 | 0.00725 | 0.00649 | 0.00213 | -0.09768343 |
| ECT | 'Ectorhinal area' | 0.00795 | 0.00931 | 0.00745 | 0.00857 | 0.00612 | 0.00769 | 0.00795 | 0.00586 | 0.00671 | 0.035101929 |
| OLF | 'Olfactory areas' | 0.00492 | 0.00575 | 0.00599 | 0.00556 | 0.00532 | 0.00763 | 0.00842 | 0.00863 | 0.00537 | -0.043490928 |
| AOB | 'Accessory olfactory bulb' | 0.01089 | 0.00569 | 0.00881 | 0.00171 | 0.01672 | 0.01461 | 0.01538 | 0.01831 | 0.00846 | -0.264757864 |
| AON | 'Anterior olfactory nucleus' | 0.00686 | 0.00813 | 0.00681 | 0.00470 | 0.00820 | 0.00641 | 0.00781 | 0.00829 | 0.00751 | -0.033765161 |
| TT | 'Taenia tecta' | 0.00506 | 0.00578 | 0.00542 | 0.00713 | 0.00679 | 0.00427 | 0.00657 | 0.00366 | 0.00475 | 0.027183022 |
| DP | 'Dorsal peduncular area' | 0.00539 | 0.00709 | 0.00514 | 0.00732 | 0.00504 | 0.00280 | 0.00348 | 0.00452 | 0.00497 | 0.076644074 |
| PIR | 'Piriform area' | 0.00570 | 0.00659 | 0.00568 | 0.00681 | 0.00724 | 0.00842 | 0.00684 | 0.00783 | 0.00645 | -0.025705975 |
| NLOT | 'Nucleus of the lateral olfactory tract' | 0.00714 | 0.01210 | 0.00798 | 0.01244 | 0.00686 | 0.01035 | 0.01906 | 0.01298 | 0.00120 | -0.002643787 |
| COAa | 'Cortical amygdalar area, anterior part' | 0.01639 | 0.01336 | 0.01091 | 0.01295 | 0.01684 | 0.01583 | 0.01346 | 0.01490 | 0.00913 | -0.036940496 |
| COAp | 'Cortical amygdalar area, posterior part' | 0.00404 | 0.00242 | 0.00245 | 0.00338 | 0.00700 | 0.00756 | 0.00731 | 0.00361 | 0.00393 | -0.087207555 |
| PAA | 'Piriform-amygdalar area' | 0.00508 | 0.00361 | 0.00247 | 0.00357 | 0.00471 | 0.00508 | 0.00541 | 0.00446 | 0.00240 | -0.028309203 |
| TR | 'Postpiriform transition area' | 0.00213 | 0.00205 | 0.00190 | 0.00208 | 0.00250 | 0.00337 | 0.00375 | 0.00322 | 0.00205 | -0.027672461 |
| HPF | 'Hippocampal formation' | 0.00396 | 0.00541 | 0.00573 | 0.00484 | 0.00599 | 0.00474 | 0.00726 | 0.00498 | 0.00450 | -0.016172825 |
| CA1 | 'Field CA1' | 0.00134 | 0.00221 | 0.00254 | 0.00149 | 0.00281 | 0.00084 | 0.00169 | 0.00214 | 0.00194 | 0.002239642 |
| CA2 | 'Field CA2' | 0.00175 | 0.00130 | 0.00250 | 0.00029 | 0.00395 | 0.00053 | 0.00070 | 0.00308 | 0.00281 | -0.019062821 |
| CA3 | 'Field CA3' | 0.00241 | 0.00191 | 0.00348 | 0.00154 | 0.00386 | 0.00134 | 0.00233 | 0.00376 | 0.00305 | -0.014178478 |
| DG | 'Dentate gyrus' | 0.00226 | 0.00362 | 0.00414 | 0.00179 | 0.00368 | 0.00210 | 0.00325 | 0.00283 | 0.00312 | -0.008124236 |
| ENTl | 'Entorhinal area, lateral part' | 0.00401 | 0.00483 | 0.00410 | 0.00379 | 0.00475 | 0.00410 | 0.00667 | 0.00509 | 0.00423 | -0.021597379 |
| ENTm | 'Entorhinal area, medial part, dorsal zone' | 0.00584 | 0.00466 | 0.00363 | 0.00386 | 0.00671 | 0.00479 | 0.00650 | 0.00624 | 0.00456 | -0.035671938 |
| PAR | 'Parasubiculum' | 0.00644 | 0.00538 | 0.00340 | 0.00274 | 0.00647 | 0.00375 | 0.00632 | 0.00793 | 0.00333 | -0.035135568 |
| POST | 'Postsubiculum' | 0.01324 | 0.01417 | 0.01642 | 0.01697 | 0.00478 | 0.00554 | 0.00838 | 0.00736 | 0.01051 | 0.222701275 |
| PRE | 'Presubiculum' | 0.00383 | 0.00671 | 0.00743 | 0.00545 | 0.00214 | 0.00397 | 0.00571 | 0.00371 | 0.00428 | 0.041685722 |
| SUB | 'Subiculum' | 0.00240 | 0.00575 | 0.00532 | 0.00663 | 0.00227 | 0.00139 | 0.00299 | 0.00235 | 0.00345 | 0.088154077 |
| CTXsp | 'Cortical subplate' | 0.00533 | 0.00589 | 0.00465 | 0.00498 | 0.00616 | 0.00363 | 0.00443 | 0.00675 | 0.00626 | 0.009272606 |
| CLA | 'Claustrum' | 0.00658 | 0.00664 | 0.00662 | 0.01286 | 0.00539 | 0.00393 | 0.00476 | 0.00741 | 0.00773 | 0.120423187 |
| EP | 'Endopiriform nucleus' | 0.00505 | 0.00524 | 0.00450 | 0.00593 | 0.00547 | 0.00280 | 0.00450 | 0.00473 | 0.00471 | 0.035302471 |
| LA | 'Lateral amygdalar nucleus' | 0.00565 | 0.00555 | 0.00542 | 0.00690 | 0.00419 | 0.00330 | 0.00466 | 0.00408 | 0.00419 | 0.05684276 |
| BLAa | 'Basolateral amygdalar nucleus, anterior pa | 0.00645 | 0.00773 | 0.00673 | 0.00950 | 0.00579 | 0.00438 | 0.00461 | 0.00607 | 0.00636 | 0.084075243 |
| BLAp | 'Basolateral amygdalar nucleus, posterior p | 0.00356 | 0.00367 | 0.00300 | 0.00449 | 0.00326 | 0.00235 | 0.00339 | 0.00388 | 0.00365 | 0.024362047 |
| BLAv | 'Basolateral amygdalar nucleus, ventral pa | 0.00432 | 0.00434 | 0.00335 | 0.00359 | 0.00315 | 0.00141 | 0.00222 | 0.00434 | 0.00458 | 0.032966479 |
| BMAa | 'Basomedial amygdalar nucleus, anterior p | 0.01355 | 0.01414 | 0.01157 | 0.01650 | 0.01553 | 0.01445 | 0.00892 | 0.01145 | 0.01030 | 0.062785232 |
| BMAp | 'Basomedial amygdalar nucleus, posterior | 0.00611 | 0.00560 | 0.00480 | 0.00618 | 0.00807 | 0.00590 | 0.00615 | 0.00628 | 0.00523 | -0.011103236 |
| PA | 'Posterior amygdalar nucleus' | 0.00535 | 0.00515 | 0.00525 | 0.00364 | 0.01016 | 0.00656 | 0.00698 | 0.00609 | 0.00536 | -0.069592379 |
| STR | 'Striatum' | 0.00664 | 0.00685 | 0.00674 | 0.00767 | 0.00619 | 0.00583 | 0.00541 | 0.00565 | 0.00679 | 0.034896512 |
| CP | 'Caudoputamen' | 0.00167 | 0.00200 | 0.00235 | 0.00132 | 0.00264 | 0.00054 | 0.00106 | 0.00181 | 0.00199 | 0.007421866 |
| STRv | 'Striatum ventral region' | 0.00476 | 0.00462 | 0.00411 | 0.00445 | 0.00332 | 0.00415 | 0.00271 | 0.00393 | 0.00485 | 0.020843718 |
| ACB | 'Nucleus accumbens' | 0.00322 | 0.00284 | 0.00301 | 0.00245 | 0.00472 | 0.00205 | 0.00281 | 0.00358 | 0.00346 | -0.008432722 |
| OT | 'Olfactory tubercle' | 0.00314 | 0.00298 | 0.00293 | 0.00303 | 0.00274 | 0.00394 | 0.00355 | 0.00280 | 0.00211 | -0.007936449 |
| LSX | 'Lateral septal complex' | 0.00930 | 0.00893 | 0.00836 | 0.01126 | 0.00809 | 0.00740 | 0.00941 | 0.00883 | 0.01014 | 0.043668516 |
| sAMY | 'Striatum-like amygdalar nuclei' | 0.00989 | 0.01227 | 0.01010 | 0.01406 | 0.01146 | 0.01028 | 0.00892 | 0.01123 | 0.00888 | 0.064916817 |
| CEA | 'Central amygdalar nucleus' | 0.00527 | 0.00833 | 0.00626 | 0.00894 | 0.00418 | 0.00451 | 0.00482 | 0.00564 | 0.00696 | 0.08130338 |
| MEA | 'Medial amygdalar nucleus' | 0.00940 | 0.01097 | 0.00998 | 0.01304 | 0.02121 | 0.01647 | 0.01718 | 0.01298 | 0.00655 | -0.112017212 |
| PAL | 'Pallidum' | 0.00384 | 0.00363 | 0.00355 | 0.00307 | 0.00247 | 0.00315 | 0.00253 | 0.00349 | 0.00331 | 0.010563305 |
| GPe | 'Globus pallidus, external segment' | 0.00214 | 0.00332 | 0.00310 | 0.00287 | 0.00145 | 0.00094 | 0.00119 | 0.00185 | 0.00266 | 0.039231967 |
| GPi | 'Globus pallidus, internal segment' | 0.00192 | 0.00090 | 0.00111 | 0.00162 | 0.00097 | 0.00031 | 0.00088 | 0.00279 | 0.00299 | 0.008742267 |

| | | | | | | | | | | | |
|---|---|---|---|---|---|---|---|---|---|---|---|
| SI | 'Substantia innominata' | 0.00347 | 0.00299 | 0.00265 | 0.00115 | 0.00216 | 0.00195 | 0.00279 | 0.00333 | 0.00276 | -0.009632939 |
| MA | 'Magnocellular nucleus' | 0.00367 | 0.00602 | 0.00490 | 0.00348 | 0.00277 | 0.00337 | 0.00552 | 0.00499 | 0.00360 | 0.00716399 |
| MS | 'Medial septal nucleus' | 0.00694 | 0.00339 | 0.01084 | 0.00550 | 0.00426 | 0.00476 | 0.00528 | 0.00608 | 0.00812 | 0.007829488 |
| NDB | 'Diagonal band nucleus' | 0.00501 | 0.00455 | 0.00526 | 0.00173 | 0.00412 | 0.00717 | 0.00604 | 0.00675 | 0.00448 | -0.071880261 |
| TRS | 'Triangular nucleus of septum' | 0.00998 | 0.00868 | 0.00923 | 0.00689 | 0.01098 | 0.00460 | 0.00585 | 0.00739 | 0.00833 | 0.029297242 |
| BST | 'Bed nuclei of the stria terminalis' | 0.00875 | 0.00725 | 0.00832 | 0.00894 | 0.01193 | 0.00882 | 0.00984 | 0.01120 | 0.00822 | -0.036085714 |
| TH | 'Thalamus' | 0.00700 | 0.00644 | 0.00708 | 0.00737 | 0.00684 | 0.00642 | 0.01036 | 0.00746 | 0.00792 | -0.016620487 |
| VENT | 'Ventral group of the dorsal thalamus' | 0.00451 | 0.00480 | 0.00423 | 0.00280 | 0.00213 | 0.00128 | 0.00249 | 0.00282 | 0.00337 | 0.038456036 |
| SPF | 'Subparafascicular nucleus' | 0.01247 | 0.01320 | 0.01130 | 0.00846 | 0.00654 | 0.01845 | 0.01824 | 0.01183 | 0.01542 | -0.11762581 |
| SPA | 'Subparafascicular area' | 0.01567 | 0.00236 | 0.01387 | 0.00589 | 0.00828 | 0.01466 | 0.01564 | 0.01278 | 0.01497 | -0.168073951 |
| PP | 'Peripeduncular nucleus' | 0.01063 | 0.00597 | 0.00591 | 0.01566 | 0.01034 | 0.00721 | 0.00911 | 0.00686 | 0.00588 | 0.088341865 |
| GENd | 'Geniculate group, dorsal thalamus' | 0.00242 | 0.00415 | 0.00375 | 0.00334 | 0.00237 | 0.00077 | 0.00203 | 0.00176 | 0.00316 | 0.044172079 |
| LAT | 'Lateral group of the dorsal thalamus' | 0.00250 | 0.00284 | 0.00341 | 0.00309 | 0.00177 | 0.00141 | 0.00282 | 0.00244 | 0.00259 | 0.023148429 |
| ATN | 'Anterior group of the dorsal thalamus' | 0.00519 | 0.00424 | 0.00420 | 0.00530 | 0.00352 | 0.00181 | 0.00305 | 0.00558 | 0.00393 | 0.046471398 |
| MED | 'Medial group of the dorsal thalamus' | 0.00643 | 0.00539 | 0.00580 | 0.00567 | 0.00590 | 0.00269 | 0.00617 | 0.00962 | 0.00611 | 0.011774784 |
| MTN | 'Midline group of the dorsal thalamus' | 0.01531 | 0.00642 | 0.01415 | 0.01983 | 0.01429 | 0.00987 | 0.01604 | 0.01710 | 0.01310 | 0.048278267 |
| ILM | 'Intralaminar nuclei of the dorsal thalamus | 0.00884 | 0.00365 | 0.00683 | 0.00990 | 0.00480 | 0.00348 | 0.00468 | 0.00800 | 0.00648 | 0.076175985 |
| RT | 'Reticular nucleus of the thalamus' | 0.00132 | 0.00201 | 0.00237 | 0.00046 | 0.00163 | 0.00080 | 0.00069 | 0.00210 | 0.00274 | -0.002487358 |
| IGL | 'Intergeniculate leaflet of the lateral genicu | 0.01314 | 0.03115 | 0.01187 | 0.02332 | 0.01494 | 0.01288 | 0.00119 | 0.01039 | 0.00952 | 0.334251272 |
| LGv | 'Ventral part of the lateral geniculate comp | 0.01555 | 0.02715 | 0.02302 | 0.01775 | 0.01212 | 0.02523 | 0.02195 | 0.00559 | 0.01881 | 0.028825121 |
| SubG | 'Subgeniculate nucleus' | 0.00190 | 0.00309 | 0.00071 | 0.00000 | 0.00000 | 0.00362 | 0.00336 | 0.00158 | 0.00287 | -0.036710927 |
| MH | 'Medial habenula' | 0.00368 | 0.01128 | 0.00696 | 0.00727 | 0.01706 | 0.00959 | 0.01703 | 0.01421 | 0.01459 | -0.144843425 |
| LH | 'Lateral habenula' | 0.00616 | 0.00774 | 0.00748 | 0.00795 | 0.00789 | 0.00729 | 0.01676 | 0.01059 | 0.00692 | -0.061418169 |
| HY | 'Hypothalamus' | 0.01377 | 0.01129 | 0.01065 | 0.01031 | 0.01217 | 0.01384 | 0.01229 | 0.01174 | 0.01179 | -0.042971439 |
| PVZ | 'Periventricular zone' | 0.01060 | 0.00433 | 0.00910 | 0.01273 | 0.02051 | 0.02009 | 0.00928 | 0.01687 | 0.01777 | -0.170407324 |
| PVR | 'Periventricular region' | 0.01402 | 0.01037 | 0.01319 | 0.01096 | 0.01651 | 0.01710 | 0.01279 | 0.01596 | 0.01767 | -0.110392965 |
| AHN | 'Anterior hypothalamic nucleus' | 0.01557 | 0.01621 | 0.01273 | 0.00789 | 0.00888 | 0.01082 | 0.01132 | 0.00994 | 0.01038 | 0.02268004 |
| MBO | 'Mammillary body' | 0.01193 | 0.00563 | 0.00944 | 0.01050 | 0.01058 | 0.01124 | 0.00607 | 0.00677 | 0.01010 | 0.001525472 |
| MPN | 'Medial preoptic nucleus' | 0.01127 | 0.00926 | 0.00839 | 0.00899 | 0.01433 | 0.01846 | 0.01633 | 0.01306 | 0.01355 | -0.169420536 |
| PMd | 'Dorsal premammillary nucleus' | 0.01809 | 0.00974 | 0.01039 | 0.00847 | 0.00900 | 0.01067 | 0.00838 | 0.01479 | 0.00802 | 0.000794977 |
| PMv | 'Ventral premammillary nucleus' | 0.01750 | 0.01038 | 0.01153 | 0.01234 | 0.01618 | 0.01865 | 0.00808 | 0.01016 | 0.00770 | -0.033526476 |
| PVHd | 'Paraventricular hypothalamic nucleus, des | 0.01828 | 0.01887 | 0.02188 | 0.01674 | 0.02752 | 0.02877 | 0.01604 | 0.01753 | 0.01998 | -0.134831884 |
| VMH | 'Ventromedial hypothalamic nucleus' | 0.00870 | 0.00783 | 0.00642 | 0.00425 | 0.01463 | 0.01064 | 0.01282 | 0.00716 | 0.00558 | -0.129787319 |
| PH | 'Posterior hypothalamic nucleus' | 0.02023 | 0.01330 | 0.01949 | 0.01719 | 0.02042 | 0.01561 | 0.02334 | 0.02387 | 0.02517 | -0.085038582 |
| LZ | 'Hypothalamic lateral zone' | 0.01105 | 0.01090 | 0.00820 | 0.00934 | 0.00835 | 0.00999 | 0.00790 | 0.00919 | 0.00832 | 0.022938553 |
| ZI | 'Zona incerta' | 0.00710 | 0.00831 | 0.00769 | 0.00768 | 0.00531 | 0.00723 | 0.00736 | 0.00662 | 0.00581 | 0.026064853 |
| ME | 'Median eminence' | 0.00339 | 0.00000 | 0.00069 | 0.01314 | 0.00655 | 0.00588 | 0.00000 | 0.00179 | 0.00621 | 0.085033276 |
| MB | 'Midbrain' | 0.00819 | 0.00803 | 0.00853 | 0.00534 | 0.00711 | 0.00767 | 0.01050 | 0.00803 | 0.00768 | -0.041608705 |
| MBsen | 'Midbrain, sensory related' | 0.00453 | 0.00667 | 0.00523 | 0.00403 | 0.00418 | 0.00289 | 0.00501 | 0.00511 | 0.00523 | 0.019351723 |
| SCs | 'Superior colliculus, sensory related' | 0.01871 | 0.02157 | 0.01729 | 0.00805 | 0.01617 | 0.02364 | 0.02120 | 0.01231 | 0.01797 | -0.155228802 |
| IC | 'Inferior colliculus' | 0.00942 | 0.01409 | 0.01171 | 0.00469 | 0.00842 | 0.01516 | 0.01726 | 0.00796 | 0.01318 | -0.127422218 |
| MBmot | 'Midbrain, motor related' | 0.00636 | 0.00717 | 0.00599 | 0.00276 | 0.00499 | 0.00450 | 0.00496 | 0.00594 | 0.00517 | -0.007770485 |
| SNr | 'Substantia nigra, reticular part' | 0.00202 | 0.00175 | 0.00250 | 0.00006 | 0.00066 | 0.00094 | 0.00191 | 0.00122 | 0.00115 | -0.006237696 |
| VTA | 'Ventral tegmental area' | 0.00532 | 0.00523 | 0.00604 | 0.00215 | 0.00382 | 0.00349 | 0.00433 | 0.00480 | 0.00530 | -0.00998256 |
| SCm | 'Superior colliculus, motor related' | 0.01119 | 0.01217 | 0.00964 | 0.00626 | 0.01214 | 0.00914 | 0.01416 | 0.01191 | 0.01042 | -0.066863714 |
| PAG | 'Periaqueductal gray' | 0.01000 | 0.01097 | 0.01116 | 0.00303 | 0.00830 | 0.00861 | 0.01019 | 0.01067 | 0.01087 | -0.068273086 |
| PRC | 'Precommissural nucleus' | 0.01884 | 0.01606 | 0.01362 | 0.01796 | 0.01779 | 0.02228 | 0.01704 | 0.02427 | 0.01531 | -0.063474231 |
| APN | 'Anterior pretectal nucleus' | 0.01000 | 0.01157 | 0.00756 | 0.00741 | 0.00604 | 0.00455 | 0.00682 | 0.00725 | 0.00741 | 0.071463487 |
| MPT | 'Medial pretectal area' | 0.01774 | 0.01720 | 0.00719 | 0.00494 | 0.01105 | 0.02291 | 0.00679 | 0.00559 | 0.01402 | -0.101262366 |
| SNc | 'Substantia nigra, compact part' | 0.00273 | 0.00230 | 0.00396 | 0.00147 | 0.00316 | 0.00170 | 0.00138 | 0.00306 | 0.00146 | 0.000219909 |
| PPN | 'Pedunculopontine nucleus' | 0.00164 | 0.00488 | 0.00336 | 0.00000 | 0.00161 | 0.00231 | 0.00241 | 0.00294 | 0.00328 | -0.014663994 |
| IF | 'Interfascicular nucleus raphe' | 0.00160 | 0.00000 | 0.01418 | 0.00353 | 0.00945 | 0.00662 | 0.00472 | 0.00422 | 0.00914 | -0.083311538 |
| P | 'Pons' | 0.00359 | 0.00649 | 0.00703 | 0.00079 | 0.00698 | 0.01243 | 0.00629 | 0.00517 | 0.00818 | -0.137515294 |
| P-sen | 'Pons, sensory related' | 0.00323 | 0.00619 | 0.00467 | 0.00019 | 0.00563 | 0.00667 | 0.00587 | 0.00491 | 0.00630 | -0.087427607 |
| NLL | 'Nucleus of the lateral lemniscus' | 0.00323 | 0.00311 | 0.00431 | 0.00009 | 0.00320 | 0.00594 | 0.00578 | 0.00135 | 0.00486 | -0.078761616 |
| P-mot | 'Pons, motor related' | 0.00635 | 0.00735 | 0.00841 | 0.00246 | 0.00877 | 0.01150 | 0.00862 | 0.01063 | 0.00995 | -0.131946985 |
| P-sat | 'Pons, behavioral state related' | 0.00303 | 0.00373 | 0.00511 | 0.00074 | 0.00237 | 0.00384 | 0.00369 | 0.00301 | 0.00428 | -0.032647357 |
| MY | 'Medulla' | 0.00065 | 0.00164 | 0.00124 | 0.00019 | 0.00230 | 0.00412 | 0.00244 | 0.00091 | 0.00129 | -0.050690308 |
| MY-sen | 'Medulla, sensory related' | 0.00098 | 0.00270 | 0.00166 | 0.00007 | 0.00556 | 0.00717 | 0.00601 | 0.00299 | 0.00180 | -0.115454626 |
| MY-mot | 'Medulla, motor related' | 0.00021 | 0.00188 | 0.00101 | 0.00003 | 0.00119 | 0.00123 | 0.00130 | 0.00063 | 0.00116 | -0.013426442 |
